# Evolution and functional dynamics of dehydrins in model *Brachypodium* grasses

**DOI:** 10.1101/2021.09.03.458816

**Authors:** M.A. Decena, S. Galvez-Rojas, F. Agostini, R. Sancho, B. Contreras-Moreira, D. L. Des Marais, P. Hernández, P. Catalán

## Abstract

Dehydration proteins (dehydrins, DHNs) confer tolerance to water-stress deficit to plants, thus playing a fundamental role in plant response and adaptation to water-deprivation stressful environments. We have performed a comparative genomics and evolutionary study of DHN genes in four model *Brachypodium* grass species, and a drought-induced functional analysis in 32 ecotypes of the flagship species *B. distachyon,* to gain insight into the origins and dynamics of these proteins and the correlated drought-mediated phenotypic responses in ecotypes showing different hydric requirements. Genomic sequence analysis detected 10 types of dehydrin genes (*Bdhn*) across the *Brachypodium* species, totalling 47 genes. Domain and conserved motif contents of peptides encoded by *Bdhn* genes revealed eight protein architectures, YSɸK_2_ being the most common architecture. *Bdhn* genes were spread across several chromosomes and more frequent in syntenic chromosomes 3 and 4 of *B. distachyon*, 4 and 5 of *B. stacei* and 4 of *B. sylvaticum*. Tandem and segmental duplication events were detected for four *Bdhn* genes. Selection analysis indicated that all the *Bdhn* genes were constrained by purifying selection. Three upstream *cis*-regulatory motifs (BES1, MYB124, ZAT) were consistently detected in several *Bdhn* genes. Functional analysis in 32 natural accessions of *B. distachyon* demonstrated that only four *Bdhn* genes (*Bdhn*1, *Bdhn*2, *Bdhn*3, *Bdhn*7) were expressed in mature leaves and that all of them were significantly more highly expressed in plants under drought conditions. These genes corresponded to wheat orthologs that were also significantly more expressed under drought stress. *Brachypodium* dehydrin expression was significantly correlated with drought-response phenotypic traits (plant biomass, leaf carbon and proline contents and WUE increases, leaf water and nitrogen content changes) being more pronounced in drought-tolerant ecotypes. *Bdhn* expression, associated phenotypic trait changes and climate niche variation did not show significant phylogenetic signal when tested in the *B. distachyon* genealogical-species tree. By contrast, some of them showed low or marginal significant phylogenetic signal when tested in the *B. distachyon Bdhn* tree, suggesting that *Bdhn* gene evolution is partially related to adaptation to drought in this species. Our results demonstrate that dehydrin composition and regulation is a key factor determining the acquisition of water-stress tolerance in grasses.

## Introduction

Water deprivation is one of the main abiotic stresses that affect plant development and fitness (Hossain, et al, 2016; Des Marais & Juenger, 2017). Water deficit stress in plants is mostly caused by low soil water content but also by other abiotic stresses such as salinity and extreme temperature (Mahajan & Tuteja, 2005; Hanin et al., 2011) which sometimes interact (Des Marais et al. 2017), forcing the plants to adapt and survive in a wide range of environmental conditions (Tommasini et al., 2008; Hanin et al., 2011). Plant species adapted to dry environments have developed mechanisms to protect their cells from water stress deficit. Among the several physiological and genomic regulatory mechanisms triggered by water limitation in plants, there is an almost universal response in the upregulation of dehydrins (Reddy et al., 2004; Hanin et al., 2011; Hossain, 2016). Dehydrins (DHN) belong to group 2 LEA (Late-Embryogenesis-Abundant) proteins (Wang et al., 2014), and are intrinsically disordered hydrophilic proteins that acquire structure when bound to ligands, such as membranes, acting as chaperones that impede the aggregation or inactivation of other proteins under desiccation to maintain the biological activity of the cell (Graether & Boddington, 2014; Verma et al., 2017). They show a high hydration capacity and can also bind large quantities of cations, retaining water in the drying cells and preventing ionic unbalance and protein denaturation. DHNs are also called RAB proteins because they are usually responsive to abscisic acid (Verma et al., 2017). DHNs accumulate in all vegetative tissues under water stress though with different specificities (Hanin et al., 2011), and also during seed development (Graether & Boddington, 2014).

The dehydrin protein family is characterized by a modular sequence domain (Dehydrin) that contains three conserved segments (Y, S, K; Graether & Boddington, 2014; Riley et al., 2019). The dehydrin identifier K-segment motif is present in all plant DHNs. It is a Lys-rich segment with a defined though variable sequence (XKXGXX(D/E)KIK(D/E)KXPG) that is located towards the C-terminus of the protein coding region and can have different copies (Graether & Boddington, 2014). The S-segment motif is a track of 3 to 8 serine residues (LHR(S/T)GS4-6(S/D/E)(D/E)3), that could be interrupted by an intron, and is present in a unique copy (Graether & Boddington, 2014). The S-segment is located between the K and Y segments and, when phosphorylated, transfers dehydrins from the cytosol to the nucleus (Goday et al., 1994; Riley et al., 2019). The Y-segment motif is located towards the N-terminus of the protein coding region and shows a consensus sequence of seven residues ((V/T)DEYGNP); when present, it can be in one or more copies. It has been suggested that the Y-segment may act as a chaperone binding site though its function is not well known (Riley et al., 2019). The Y, S and K segments are interspersed by other less conserved segments, named interpatter or ɸ-segments that mostly contain small, polar, and charged amino acids (Rosales et al., 2014). Recent studies have described another conserved F-segment motif across the full spectrum of seed plant DHNs (Perdiguero et al., 2014; Strimbeck, 2017). The F-segment comprises 11 amino acids and is characterized by a short palindromic sequence with hydrophobic phenylalanine residues (DRGLFDKFIGKK). This segment, when present, has a unique copy and is located between the N-terminus of the protein coding region and the S-segment. An additional NLS (Nuclear localization signal)-segment motif was found in some dehydrin families of maize (Goday et al., 1994). Different combinations of conserved motifs have been used to classify the dehydrin domain into major architectures, with five of them (K_n_, SK_n_, K_n_S, Y_n_K_n_, Y_n_SK_n_) being common across angiosperms (Riley et al., 2019). The dehydrin domain is occasionally fused with upstream DNAJ-X and DNA-J containing gene domains in some DHN genes (Riley et al., 2019). DNA-J is a heat shock protein (Hsp40) that prevents the aggregation of unfolded proteins and functions as a chaperone folding protein when associated with Hsp70 under water-stress (Henessy et al. 2005; Riley et al., 2019).

DHNs have been extensively studied in grasses due to their key role as agents conferring water-stress tolerance in cereal and forage crops (Kosová et al., 2014; Suprunova et al., 2004; Verma et al., 2017; Wang et al., 2014). Expression of dehydrins under drought stress has been positively associated with plant biomass and grain yield (Karami et al., 2013). Several *in vitro* studies have demonstrated that DHN expression enhances plant stress tolerance (Yu et al., 2018; Lv et al., 2018). Dehydrins maintain the osmotic balance of cells and their chlorophyll contents, bind metals to scavenge ROS, and bind to DNA and phospholipids (Liu et al., 2017; Yu et al. 2018). Despite these advances, some of the biological functions of DHNs have not been fully established yet (Riley et al., 2019). *Brachypodium* is a model system for grasses due to its intermediate evolutionary position between the temperate cereals and the tropical biofuel crops. Its three annual species have been selected as a model complex for polyploidy (diploids *B. distachyon* and *B. stacei* and derived allotetraploid *B. hybridum*) and one of its perennial species has been selected as a model species for perenniality (diploid *B. sylvaticum*) (Scholthof et al., 2018). Transgenic plants of the flagship species *B. distachyon* have been analyzed to identify candidate genes that enhance drought stress tolerance in plants (Lee & Kang, 2016; Ryu & Cho, 2015; Yoon et al., 2019) and inspection of an early version of the *B. distachyon* reference genome Bd21 detected several LEA2 encoding genes (Filiz et al., 2013). However, the dehydrin gene content, structure, evolution, and expression in response to drought, among species and accessions of *Brachypodium* has not been investigated yet.

Given the significance of DHNs in water stress response of grasses, we have analyzed the members of the dehydrin gene family in the four *Brachypodium* model species and in 54 *B. distachyon* ecotypes showing different hydric requirements and drought tolerances using *in silico* analysis of genome annotations. We have also performed DHN expression analysis in 32 *B. distachyon* ecotypes under different drought and watered conditions. The aims of our study were to: i) identify and characterise the *Brachypodium* dehydrin genes (*Bdhn*) and the structure and biochemical properties of the encoded proteins, comparing them with those of the close cereal crops, ii) evaluate the potential regulatory effects of their cis-regulatory motifs, iii) analyse their syntenic distributions and origins, iv) identify gene duplication events and test the functionality of paralogs, v) analyse their expression profiles under control vs dry conditions and compare them to those of close cereals, vi) correlate the protein load and composition with the phenotypic and environmental traits of the plants, and vii) test their potential phylogenetic signal.

## Material and Methods

### Identification of dehydrin sequences

Dehydrins of *B. distachyon*, *B. stacei*, *B. hybridum*, *B. sylvaticum* and other grass outgroups were identified using three searching approaches. First, the Phytozome v.12.1 database (Goodstein et al., 2012) was searched for DHN gene sequences of *B. distachyon, B.stacei, B.hybridum* and *B. sylvaticum* (Table 1). Phytozome dehydrin sequences were retrieved using BioMart to filter sequences having DHN Pfam code (Pfam00257). This search was repeated in the Ensembl (Howe et al., 2020) and Genbank (Agarwala et al., 2018) databases, aiming to retrieve all dehydrin genes present in *Brachypodium*. Redundant sequences and incomplete transcripts were deleted. Second, a consensus K-segment was used as a query sequence to search for complete DHN genes within the retrieved sequences using the BlastP tool (Altschul et al., 1990). The presence of a K-segment with a maximum threshold of 4 mismatches with respect to the query was used to characterize a protein sequence as a dehydrin. Third, orthologous dehydrin genes from five additional grass species with complete sequenced genomes were searched in Ensembl Plants, Phytozome, Genbank, and Panther (http://pantherdb.org/data/) databases using BioMart and used as reference outgroups. The Pfam00257 code was used to find DHN orthologous sequences from *Aegilops tauschii*, *Hordeum vulgare, Oryza sativa, Sorghum bicolor,* and *Zea mays*, discarding also redundant sequences or sequences without the K-segment.

**Table 1.**
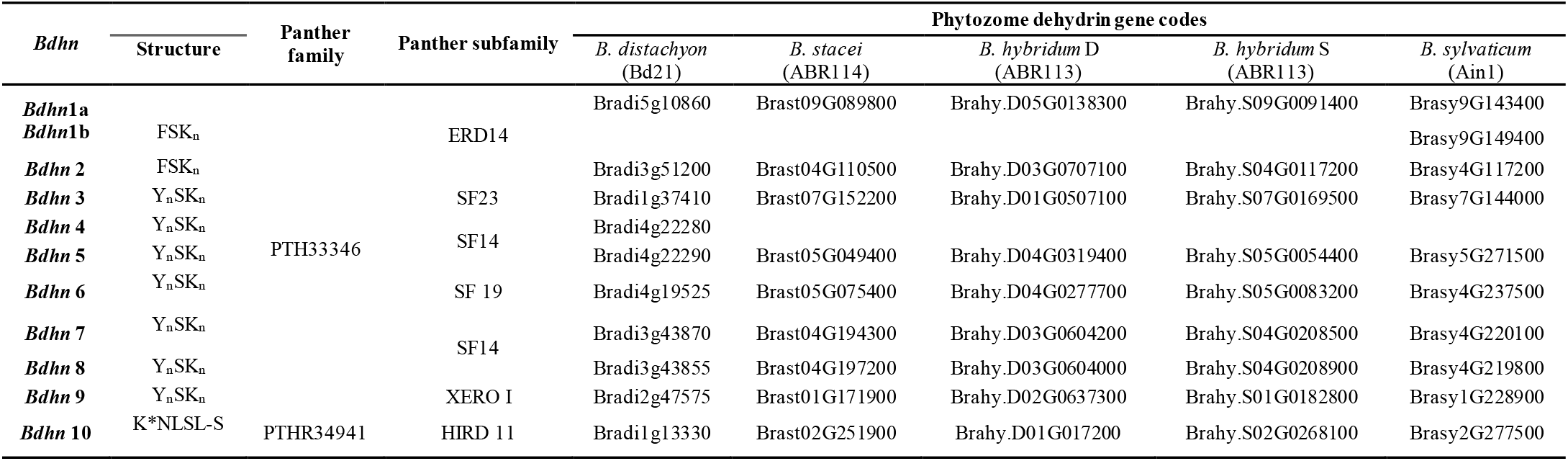
Dehydrin genes found in the studied *Brachypodium distachyon*, *B. stacei*, *B. hybridum* (subgenomes D and S) and *B. sylvaticum* species. *Brachypodium* dehydrin gene codes (*Bdhn*1-*Bdhn*10) given in this study, the protein structure and their corresponding Panther family and subfamilies gene codes. Phytozome dehydrin gene codes correspond to the respective gene numbers in the reference genome of each species deposited in Phytozome.

The theoretical molecular weight (mol. wt), isoelectric point (pI), instability index, and grand hydrophathicity average index (GRAVY) values of the different dehydrins were predicted using the ProtParam (http://web.expasy.org/protparam) tool. *In silico* dehydrin structures were modelled using the web server version of RaptorX (http://raptorx.uchicago.edu/BindingSite/; Källberg et al., 2012; Xu et al., 2020) and their structural properties were analysed using ICn3D 3D structure viewing tool (Wang et al., 2020).

### Structural analysis, conserved motifs and cis-regulatory elements

The inferred DHN polypeptide sequences were used to analyse the presence of conserved motifs and to characterize the structure of the dehydrins. A custom search tool (Supplementary Data, https://github.com/Bioflora/Brachypodium_dehydrins) was designed to find the conserved K (EKKGIMDKIKEKLPG), Y (VDEYGNP), S (SSSSS+), ɸ (EDDGQGR), F (DRGLFDKFIGKK), and NLS (KKDKKKKKEKK) motifs present in the dehydrin domain. A consensus sequence for each segment was retrieved and used as a query in a BLASTP search (https://blast.ncbi.nlm.nih.gov/Blast.cgi?PAGE=Proteins), allowing a maximum threshold of 4 mismatches with respect to the query. The dehydrin architectures were established according to the presence and distribution of their conserved motifs.

*De novo* discovery of cis-regulatory elements (CREs) was performed on *Bdhn* promoter regions, producing a set of enriched DNA motifs represented as weight matrices. CREs are binding sites for transcription factors (TFs) that accumulate around the transcriptional start site (TSS) and control gene expression. A window of −500-to-+200 bp both sides of the TSS was selected for DHN genes in all the studied *Brachypodium* species and ecotypes. We searched for over-represented motifs using RSAT::Plants (Nguyen et al., 2018) tool peak-motifs, as described in Ksouri et al. (2021) This analysis was run four times, using as genome background model the respective reference genome of each *Brachypodium* species under study (*B. distachyon* Bd21 v3.0.46, *B. stacei* ABR114 v1.1.JGI, *B. hybridum* ABR113 v1.1.JGI, *B. sylvaticum* Ain1v1.1.JGI). Significant enrichment of the discovered motifs was assessed using as negative controls promoters from the same number of randomly picked genes (Ksouri et al., 2021). Candidate motifs were chosen based on their k-mer significance and number of sites and subsequently clustered to avoid redundancies using the matrix-clustering tool (Castro-Mondragon et al., 2017). Selected motifs were finally scanned along each *Bdhn* promoter to locate potential CREs using a matrix-scan and a maximum threshold of 9 based on the median length of the 3 motif logos.

### Multiple alignments and phylogenetic analysis

Multiple sequence alignment (MSA) of the nucleotide coding sequences of all the *Brachypodium* species and other grass outgroups’ dehydrin genes was performed with ClustalW in MEGA v.5 (Tamura et al., 2011) using default settings. The start codon of each dehydrin gene was set using the Phytozome annotations and the sequences were adjusted manually to fit the reading frames. Alignments of dehydrin sequences, including exons and introns were performed with MAFFT v.7.215 (Katoh & Standley, 2013) in Geneious Prime 2021 (https://www.geneious.com/prime/). Maximum likelihood (ML) phylogenetic trees were constructed with IQTREE 1.6.12 (Trifinopoulos et al., 2016) imposing the best-fit nucleotide substitution model of each data set according to the Bayesian Information Criterion (BIC). Branch support for the best tree was estimated through 1,000 ultrafast bootstrap replicates.

### Chromosomal location, gene duplication, and selection analysis

Physical locations of the *Brachypodium* dehydrin genes in the 5 chromosomes of *B. distachyon*, 10 of *B. stacei*, and 9 of *B. sylvaticum* were obtained from Phytozome and Ensembl. They were mapped to their respective chromosomes using gff3 annotation coordinates for each dehydrin gene. Tandemly and segmentally duplicated genes were identified on the chromosomes; tandemly duplicated dehydrin genes were those distributed adjacent to an homologous dehydrin gene on the same chromosome or within a sequence distance of 50 kb (Rehman et al., 2020).

Selection rates, a measure to examine selection pressure among duplicate gene pairs, were estimated based on the dN/dS ratio (ω) over the codon alignment of the coding sequences (CDSs) of the *Brachypodium* dehydrin genes. Codon alignments were performed with ClustalW in Mega5, following the protocol indicated in https://github.com/hyphaltip/subopt-kaks. Pairwise ω comparisons were computed following Yang & Nielsen (2000); ω <1 values indicated strong purifying selection (negative selection), ω=1 neutral selection, and ω >1 accelerate evolution (positive selection).

### Clustering and phylogeny of dehydrin genes in *B. distachyon* ecotypes

Annotations of the locations of dehydrin genes in the genomes of 54 *B. distachyon* ecotypes distributed across the Mediterranean region (Supplementary Table S1) were used to map them into the five *B. distachyon* chromosomes using a custom tool (Supplementary Data, https://github.com/Bioflora/Brachypodium_dehydrins). Protein sequences were obtained from primary transcript files and specific dehydrin genes were extracted from pseudomolecules using coordinates from gff3 annotation files. The dehydrin protein sequences were aligned using BLOSUM62. Dehydrin genes of all ecotypes (including the *B. distachyon* reference genome Bd21) were classified into different clusters; each cluster contained the set of all dehydrins with a similarity of 95% or higher. To avoid potential assembly or annotation errors, only clusters containing three or more sequences were selected. Unclassified sequences were iteratively compared to the previous blocks and classified into new clusters following the procedures of the first analysis. The remaining unclassified sequences were identified manually and classified as dehydrins whenever possible. Dehydrin *Bdhn*4 and *Bdhn*5 were annotated together in certain *B. distachyon* lines. Those sequences were manually curated and its presence in the genomes corroborated using BLASTN (Supplementary Materials 1) at Brachypan database (Gordon et al., 2017). Maximum-likelihood (ML) phylogenetic analysis was performed with coding and non-coding sequences of the ten *Brachypodium* dehydrin genes across the 54 ecotypes of *B. distachyon* using IQTREE and the procedures indicated above.

### Expression analysis of dehydrin genes in *B. distachyon*

Development and tissue specific expression analysis of dehydrin genes was performed from a wide transcriptome study of 32 out of the 54 genomically sequenced ecotypes of *B. distachyon* (Sancho et al., 2021) (Supplementary Table S1). Grown plants (21 days from initial pot emergence) were subjected to watered (W) *vs* dry (D) conditions, following the experimental design described in Des Marais et al. (2017). Irrigated plants were watered to field capacity every second day whereas soil water content was reduced by ∼5% each day in dry plants. The plants under both treatments were simultaneously exposed to cool (C, daytime ∼25°C) or hot (H, daytime ∼35°C) conditions although the temperature stress conditions did not affect substantially the expression of dehydrin genes (see Results). Fully expanded leaves from up to four individuals (replicas) per ecotype and treatment were excised below the lamina, flash-frozen on liquid nitrogen and then stored at −80°C until RNA extraction. RNA isolation, sequencing and data processing followed the procedures indicated in Sancho et al. (2021). Filtered transcripts per million (TPM) values of annotated dehydrins of plants under the combined WC, WH, DC and DH treatments were extracted from the large TPM abundance database. Total TPM values were quantified with Kallisto and normalized with Sleuth according to Sancho et al. (2021) (Supplementary Table S2). The annotation of dehydrin transcripts was conducted using the *B. distachyon* Bd21 v.3.1 reference genome (http://phytozome.jgi.doe.gov/; IBI 2010). Annotated transcripts from full transcriptomes have been deposited in the European Nucleotide Archive (ENA;https://www.ebi.ac.uk/ena) under accession codes ERR6133302 to ERR6133575 (project PRJEB45871) and those of *Bdhn* genes in Github (Supplementary Data, https://github.com/Bioflora/Brachypodium_dehydrins).

Summary statistic (mean, median, SD, range) values, boxplots and whisker plots of differentially expressed (DE) dehydrin (TPM) data were computed for each ecotype and expressed dehydrin gene using the base, dplyr, ggplot2, ggpubrr, ggsignif, readxl and stats packages in R. Statistically significant differences between median values of samples under drought (W vs D) and temperature (C vs H) stresses, and within each of the W and D treatments of the drought experiment were tested. Wilcoxon pairwise difference tests for all pairs of compared samples with p-values adjusted with the Benjamini–Hochberg procedure to correct for multiple comparisons, Kruskal–Wallis rank tests for the whole group of samples within each group, and posthoc Tukey tests for among ecotypes differences were computed using the lm package and other options of R.

### Drought-induced changes in dehydrin expressions, phenotypic, physiological, and climatic niche traits, and phylogenetic signal in *Brachypodium distachyon*

The potential effect of drought on dehydrin expression levels and on correlated changes of phenotypic and physiological drought-response traits of the plants was evaluated in 32 *B. distachyon* ecotypes (Supplementary Table S1). Values of 12 drought-response traits under W and D treatments (leaf_rwc: relative water content in leaf; leaf_wc: water content in leaf; lma: leaf mass per area; pro: leaf proline content; abvrgd: above ground biomass; blwgrd: below ground biomass; ttlmass: total plant mass; rmr: root mass ratio; delta13c: carbon isotope, a proxy for lifetime integrated WUE; leafc: leaf carbon content; leafn: leaf nitrogen content; cn: leaf carbon/nitrogen ratio) were measured in the same individual samples (replicates) used in the transcriptomic analyses (Supplementary Materials 2); these phenotypic characters corresponded to those studied by Des Marais et al. (2017). Summary statistics and significance tests were computed for the 12 traits under W and D treatments following the same procedures mentioned above. Differential expression levels of the *B. distachyon* dehydrin genes in leaves of plants under W and D conditions were compared to those of wheat dehydrin genes in the available transcriptomes of wheat, a close temperate cereal for which *B. distachyon* is a model genomic system (Scholthof et al. 2018). The wheat RNAseq analyses were carried out in flag leaves of individuals under field drought stress (Galvez et al. 2019, Lv et al. 2018) and field rain shelter and greenhouse (Reddy et al. 2014, Chu et al. 2021, Konstantinov et al. 2021) experiments. Three of these wheat studies reported differentially expressed wheat dehydrins under water stress (Reddy et al. 2014, Galvez et al. 2019, Chu et al. 2021) and were used to perform the comparisons. Sixty wheat dehydrin gene sequences (IWGSC 2018) were retrieved through Blast analysis from the Wheat@URGI portal https://wheat-urgi.versailles.inrae.fr (Alaux et al. 2018). Orthology of the expressed *B. distachyon* and *Triticum aestivum* dehydrin genes was retrieved from Ensembl Plants BioMart, from Galvez et al. (2019) and from BLAST analyses performed in this study.

Environmental climate data was retrieved for the studied *B. distachyon* ecotypes from worldclim (19 temperature and precipitation variables; Supplementary Table 3). Climatic niche optima were constructed for each ecotype based on occurrence data and the first axis of the ordination of the climatic variables (PCA1) was computed with the dudi.pca function of the ade4 package (Dray & Dufour, 2007) in R. The climatic niches of the *B. distachyon* ecotypes were classified in climatic classes warm, mesic or cold according to their PCA1 values (see Results; Supplementary Table 3).

To address potential correlations between the dehydrin gene expressions and the changes in drought-response phenotypic traits, linear regression model analyses were performed for testing the effect of particular *Bdhn* gene expressions on phenotypic changes using the lm function of the R stats package.

A consensus ML phylogenetic tree of 30 *B. distachyon* ecotypes based on the expressed dehydrin genes (*Bdhn* tree) was topologically contrasted to that of the *B. distachyon* nuclear species tree based on genome-wide >3.9 million syntenic SNPs (Gordon et al., 2017) using the KH and SH tests with resampling estimated log-likelihood (RELL) optimization and 1 million bootstrap replicates in PAUP* (Swofford, 2002). We also tested for topological congruence of the *Bdhn* tree and the *B. distachyon* plastome tree based on full plastome sequences of these ecotypes (Sancho et al., 2018) using the KH and SH test approach.

Dehydrin expression level, drought-response phenotypic change and climatic niche (PCA1) variation traits were tested for phylogenetic signal using Blomberg’s K (Blomberg et al., 2003) and Pagel’s lambda (Pagel, 1999) with the *phylosig* function of the package *phytools* (Revell, 2012) in R. For both tests, values close to 1 indicate that trait values are consistent with the tree topology (phylogenetic signal) and those close to 0 that there is no influence of shared ancestry on trait values (phylogenetic independence). Phylogenetic signal was assessed on both the *B. distachyon* nuclear species tree and the *B. distachyon Bdhn* tree. Phyloheatmaps were generated for the standardized values of these continuous characters with *phytools*.

## Results

### Dehydrin genes of *Brachypodium* species and outgroup grasses

Genome searches in Phytozome and Ensembl Plants retrieved 47 dehydrin gene sequences collected from the reference genomes of the four sequenced *Brachypodium* species [*B. distachyon* (10), *B. sylvaticum* (10), *B. stacei* (9), *B. hybridum* (18; 9 from its *B. distachyon*-type D subgenome and 9 from its *B. stacei*-type S subgenome)] (Table 1; Fig. 1a). A total of 35 orthologous DHN sequences were retrieved from the reference genomes of five outgroup grass species [ *Aegilops tauschii* (9), *Hordeum vulgare* (8), *Zea mays* (7), *Oryza sativa* (6), *Sorghum bicolor* (5); Supplementary Table 4]. A new nomenclature was created for the *Brachypodium* dehydrin genes (*Bdhn*1 to *Bdhn*10) (Table 1; Fig. 1). In several instances we used the same numbers as those of the orthologous *Hordeum vulgare* DHN genes (*Bdhn* 6-7, and *Bdhn* 9-10) (ENA database: AF043086, AF043092; AF043086 and Genbank database: AY681974) and the orthologous *Oryza sativa* DHN genes (*Bdhn* 1-2 and *Bdhn* 8) (RAP database: Os02g0669100, Os11g0454300). *Bdhn* 3 was numbered according to prior annotation in *Brachypodium* (GeneBank: XM_010229280), whereas the remaining *Bdhn* genes were numbered consecutively as *Bdhn* 4 and *Bdhn* 5. The *Bdhn* genes were also classified according to Panther protein gene families (PTH33346 and PTH34941) and subfamilies (ERD14, XERO1, SF14, SF19, SF23, HIRD11) (Table 1). All *Bdhn*10 genes found in the studied species of *Brachypodium* lacked the K-segment (Fig. 1b). However, those dehydrins showed an extraordinary sequence identity with typical DHNs, including those with a modified K-segment, like DHN-13 from *H. vulgare* (Rodríguez et al., 2005). These *Brachypodium Bdhn*10 dehydrins, with architecture K*(NLS)S, belong to HIRD11 proteins. The absence of the K-segment has been also reported in four dehydrins of rice species (OnDHN6, OrDHN7, OlDHN3, OlDHN6; Verma et al., 2017).

**Figure 1.**
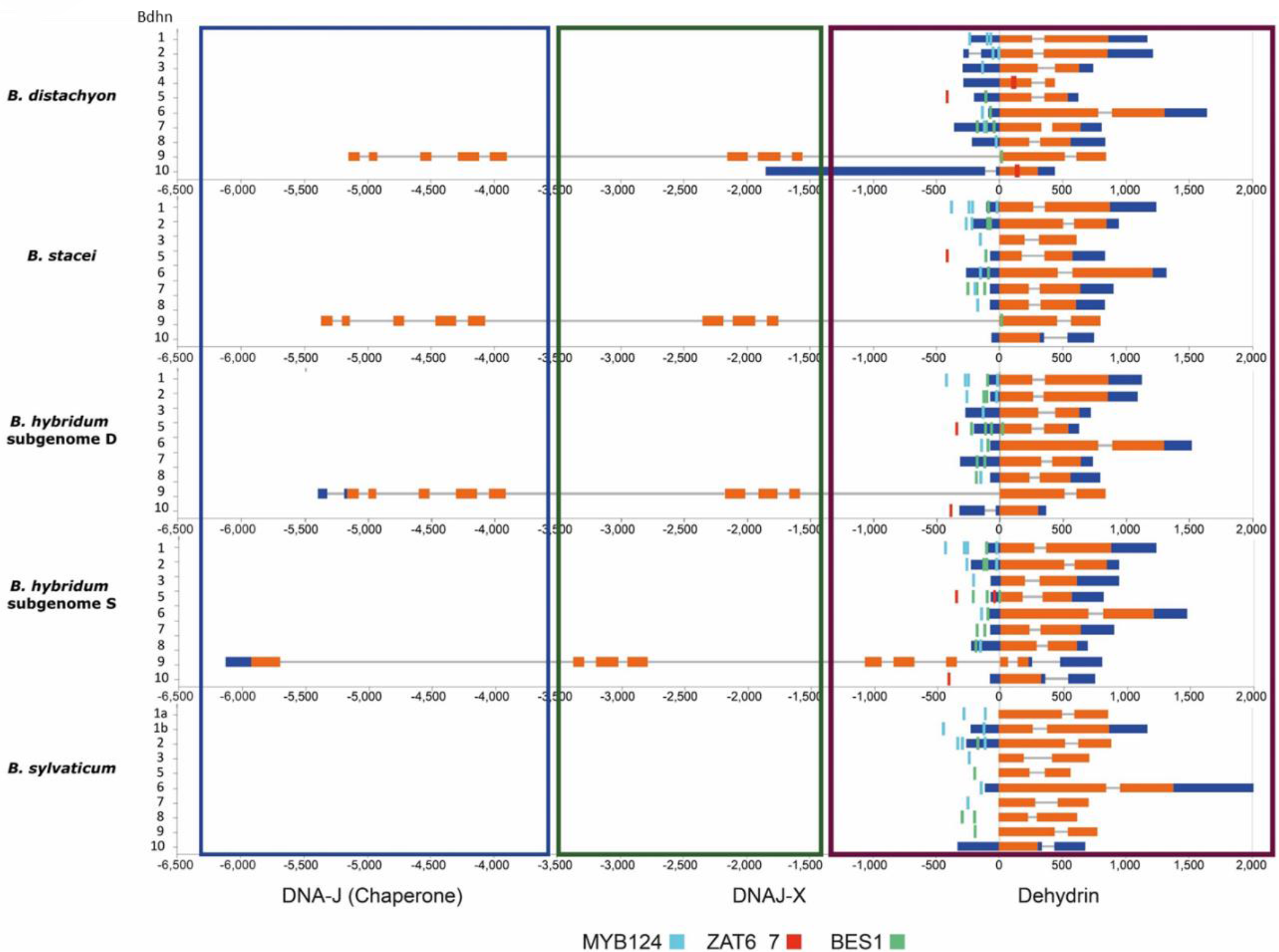
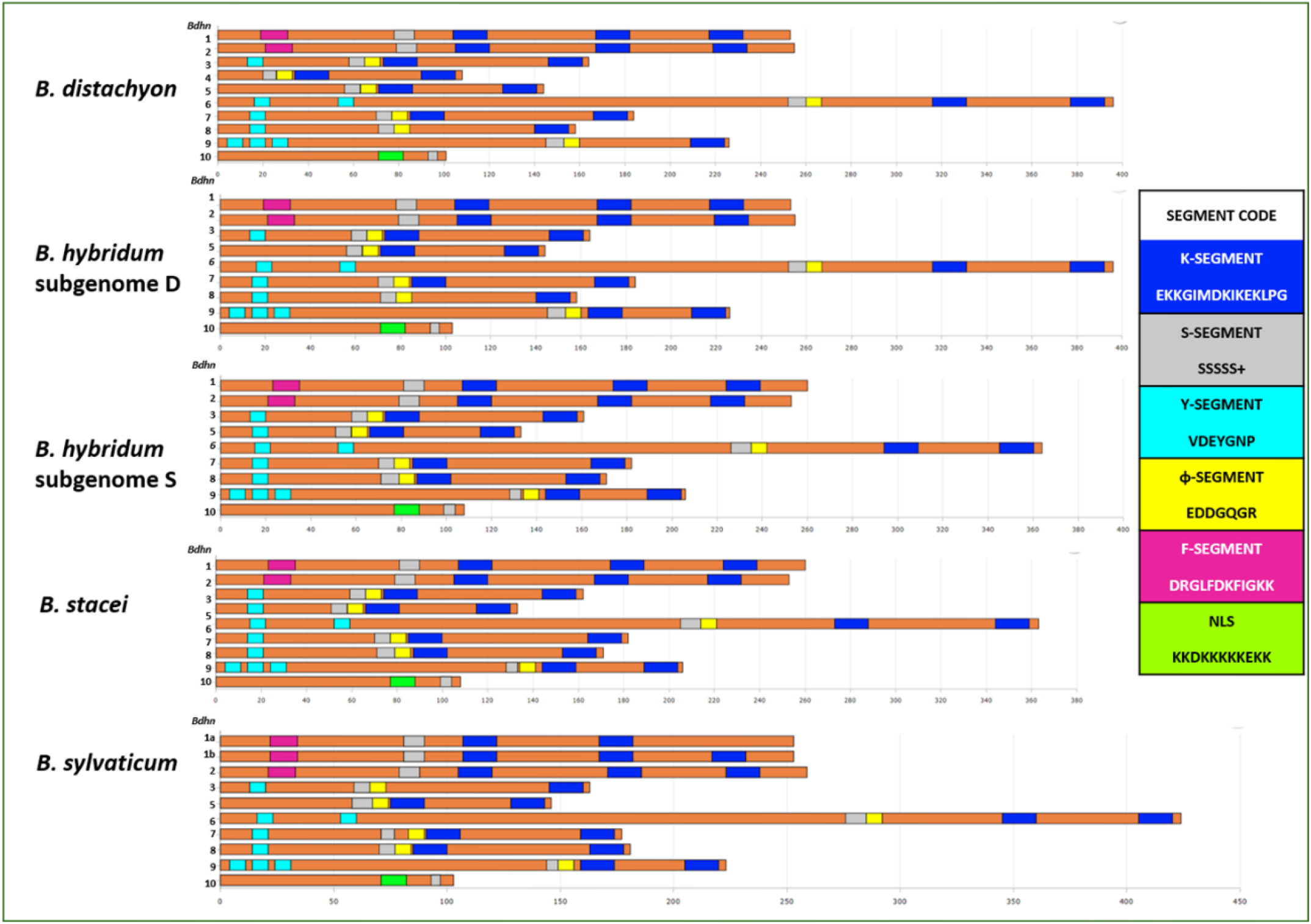
Gene structure of *Brachypodium distachyon, B. stacei, B. hybridum* (D and S subgenomes) and *B. sylvaticum* dehydrins showing: **(a)** the three detected DNA J (purple rectangle), DNA J-X (light green rectangle) and dehydrin (dark green rectangle) domains, and their CDSs (orange bars), introns (grey lines) and 5’ and 3’ UTRs (blue bars); **(b)** the conserved motifs of their CDSs with their respective K (dark blue), S (grey), Y (light blue), ɸ (yellow), F (pink) and NLS (light green) segments. *Brachypodium* dehydrin gene codes (*Bdhn*1-*Bdhn*10) correspond to those indicated in Table 1. *Cis*-regulatory elements BES1, MYB124, ZAT are mapped in the promoters of each gene (see color codes in the chart) (the figure could be also visualized in http://zeta.uma.es/public/journal/brachy/DHN_Brachy_4_varieties.html).

The *Brachypodium* dehydrins showed different lengths, ranging from 86.3 (*Bdhn*10) to 323 amino acid residues (*Bdhn*6), molecular weights from 9762.90 to 30945.54 kDa, and isoelectric points from 4.41 (*Bdhn*2) to 7.6 (*Bdhn*7) [9.4 (*Bdhn*4) in *B. distachyon*] (Table 2). All dehydrins presented a negative GRAVY value, indicating that they are hydrophilic proteins. All *Bdhn* genes except *Bdhn*5 were structurally conserved across the four *Brachypodium* species (Figs. 1a, b). *Bdhn*4 was only present in *B. distachyon*, while *Bdhn*1 was duplicated in *B. sylvaticum* (*Bdhn*1a, *Bdhn*1b) (Table 1; Figs. 1a, b). All *Bdhn* genes but *Bdhn*9 showed a single dehydrin domain. *Bdhn*9 encoded a protein of 508 aa with DHN – DNAJ-X – DNA-J domains in all species except in *B. sylvaticum,* which showed a gene consisting only of the DHN domain (222 aa) (Table 2; Fig. 1a). Eight gene architectures were found along the *Bdhn*1-*Bdhn*10 genes (FSK_2_, FSK_3_, SɸK_2_, YSɸK_2_, YSɸK, Y_3_SɸK, Y_3_SɸK_2_, NLS-K*S), being YSɸK_2_ the most common architecture present in four genes (*Bdhn*3, *Bdhn*6, *Bdhn*7, *Bdhn*8; Table 1, Fig. 1b). The 3D *Bdhn* modelling indicated that the disordered *Brachypodium* dehydrins lacked tertiary structure whereas their secondary structure consisted of different numbers of α-helices and β-sheets (Supplementary Fig. S1). All species’ dehydrins showed α-helices (1 to 4) whereas only some of them presented β-sheets (*Bdhn*1 two in all species except *B. distachyon*, *Bdhn*2 two to three in *B. sylvaticum*, and *Bdhn*6 and *Bdhn*9 three in *B. hybridum*S and *B. sylvaticum*) (Supplementary Fig. S1).

**Table 2.**
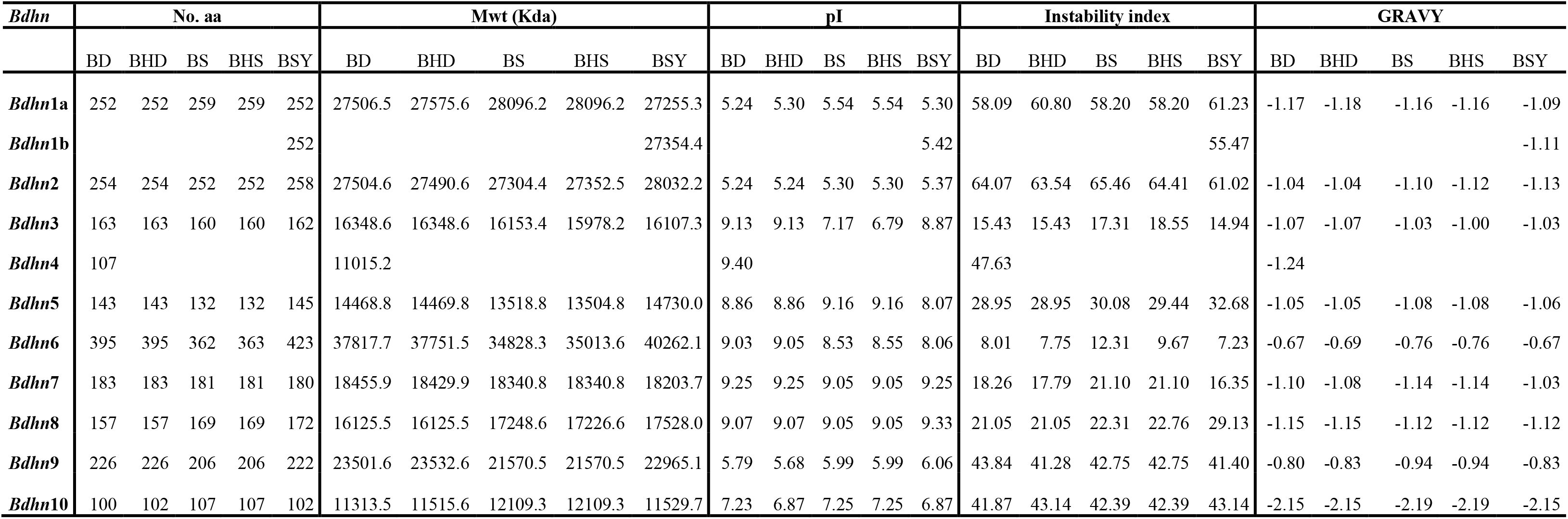
Molecular traits of *Brachypodium Bdhn* proteins. No. aa, number of aminoacids; Mwt, molecular weight; pI, isolectric point; Instability index; GRAVY, grand hydrophathicity average index. Abbreviations of species and reference genomes: BD, *B. distachyon* Bd21; BHD, *B. hybridum* D-subgenome ABR113; BS, *B. stacei* ABR114; BHS, *B. hybridum* S-subgenome ABR113; BSY, *B. sylvaticum* Ain1.

Orthology analysis indicated that *Bdhn*1 and *Bdhn*2 genes were also present in the other surveyed grasses, *Bdhn*3 in *A. tauschii, H. vulgare* and *Z. mays*, *Bdhn*4-5 and *Bdhn*7-8 in *A. tauschii, H. vulgare* and *O. sativa*, *Bdhn*6 in *A. tauschii, O. sativa, S. bicolor* and *Z. mays*, *Bdhn*9 *in A. tauschii, O. sativa, S. bicolor and Z. mays*, and *Bdhn*10 in *H. vulgare, O. sativa, S. bicolor* and *Z. mays* (Supplementary Table 4). By contrast, five DHN genes of *O. sativa* (Os01g0702500, Os11g0453900, Os01g0624400, Os01g0225600, Os03g0655400), four of *H. vulgare* (HORVU6Hr1G083960, HORVU6Hr1G011050, HORVU5Hr1G103460, HORVU3Hr1G089300) and three of *Z. mays* (Zm00001d017547, Zm00001d043730, Zm00001d013647) had no orthologous sequences in *Brachypodium*. Pairwise amino acid sequence similarities indicated that *Bdhn*4 and *Bdhn*5 were the most similar proteins, followed by *Bdhn*1 and *Bdhn*2. *Bdhn*10 was the most dissimilar dehydrin. All the dehydrins with YSɸK_2_ structure were in general highly similar to each other both in *Brachypodium* and in the other grasses.

### Cis-regulatory elements of *Bdhn* genes

A total of 60 potential cis-regulatory motifs were retrieved and subsequently filtered, comparing their k-mer significance and number of sites with the negative controls (Supplementary Fig. S2). Of them, 29 motifs were selected and clustered to avoid redundancies due to different identifications of the same CRE. The analysis consistently identified 3 clusters BES1/BZR1, MYB124 and ZAT binding sites (Table 3; Fig. 2) that are related with different drought-response signaling pathways. BES1/BZR1 and MYB124 motifs were present in all studied promoters, though more abundantly in those of the aridic *B. stacei* and *B. hybridum* species; by contrast, ZAT was predominant in the *Bdhn* promoters of mesic *B. distachyon* though was not found in those of nemoral *B. sylvaticum* (Fig. 2). MYB124 was predominant in the promoters of *Bdhn*1 and *Bdhn*2 of all studied *Brachypodium* genomes, whereas BES1/BZR1 was present in the promoters of all *Bdhn*7 genes and also found in the promoters of the *Bdhn*3, *Bdhn*5, *Bdhn*6, *Bdhn*8 and *Bdhn*9 genes of some species. By contrast, ZAT was only found in the promoters of *Bdhn* 4, *Bdhn*5, and *Bdhn*10 genes in annual species (Table 3; Fig. 2).

**Figure 2.**
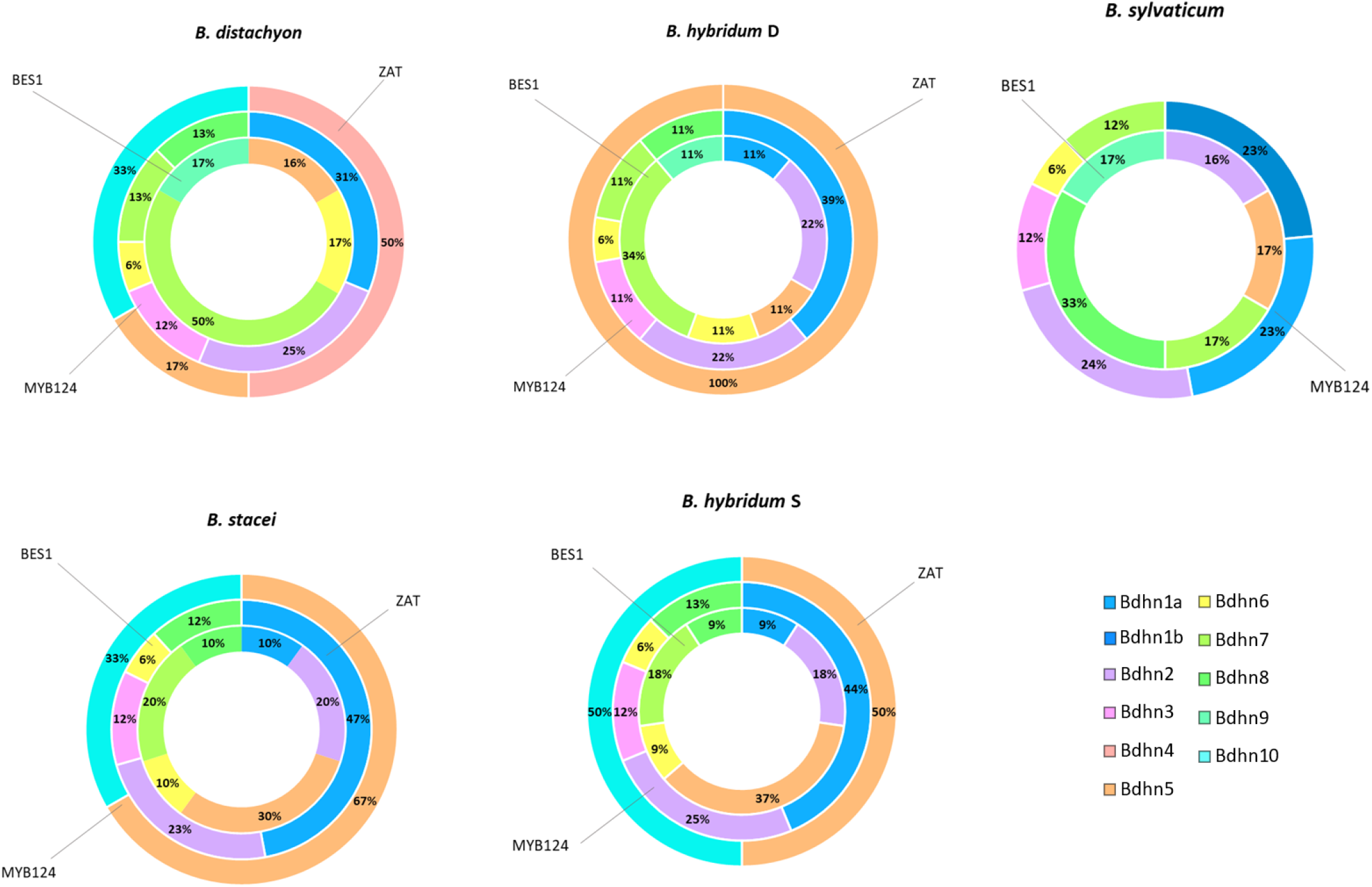
BES1/BZR1 (Basic leucine zipper), MYB124 (Myb gene protein), and ZAT (C2H2 zinc finger) *cis*-regulatory elements found in 5’-upstream promoter region (−500-to-+200 bp) of the *Brachypodium Bdhn* genes. Distributions of cis-motifs per species and reference genomes (BD, *B. distachyon* Bd21; BS, *B. stacei* ABR114; BHD, *B. hybridum* subgenome-D ABR113; BHS, *B. hybridum* subgenome-S ABR113; BS, *B. sylvaticum* Ain1) and per *Bdhn* gene promoter (dehydrin genes *Bdhn*1 to *Bdhn*10).

**Table 3.**
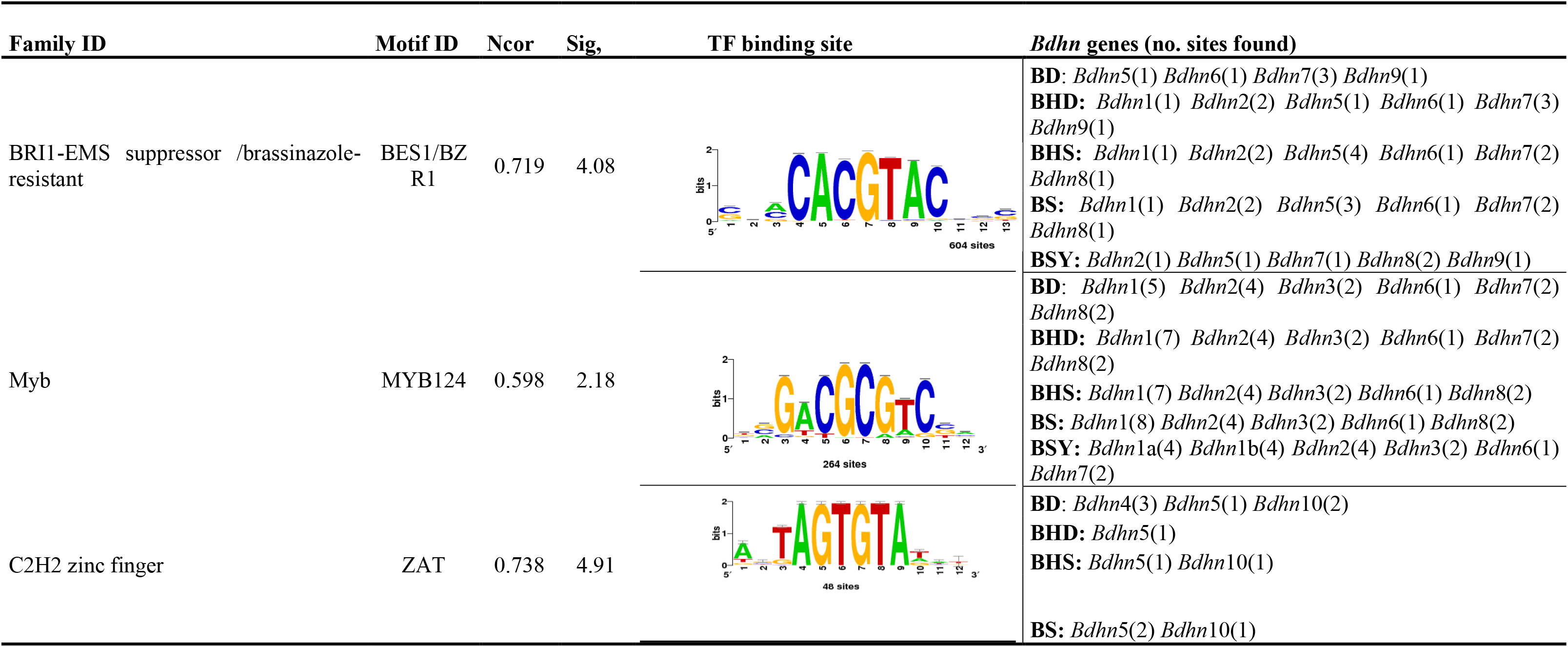
Upstream *cis*-regulatory elements (CREs) found in the promoter region (−500-to-+200 bp) of the *Bdhn* genes of *Brachypodium distachyon*, *B. stacei*, *B. hybridum* (subgenomes D and S) and *B. sylvaticum* using Rsat::plants tools and the corresponding reference genome as background. Family identification, motif code (ID), N-cor (normalized correlation) and Sig (significance value) for the highest hit, matches to transcription factor (TF) binding sites and *Bdhn* genes with the number of sites found within each specie and gene. Species and reference genomes: BD, *B. distachyon* Bd21; BS, *B. stacei* ABR114; BHD, *B. hybridum* subgenome D ABR113; BHS, *B. hybridum* subgenome S ABR113; BS, *B. sylvaticum* Ain1. Mapping positions of these cis-regulatory motifs are indicated in Figure 1a.

| Family ID | Motif ID | Ncor | Sig, | TF binding site | *Bdhn* genes (no. sites found) |
| --- | --- | --- | --- | --- | --- |
| BRI1-EMS suppressor /brassinazole-resistant | BES1/BZR1 | 0.719 | 4.08 |  | **BD:** *Bdhn*5(1) *Bdhn*6(1) *Bdhn*7(3) *Bdhn*9(1)<br>**BHD:** *Bdhn*1(1) *Bdhn*2(2) *Bdhn*5(1) *Bdhn*6(1) *Bdhn*7(3) *Bdhn*9(1)<br>**BHS:** *Bdhn*1(1) *Bdhn*2(2) *Bdhn*5(4) *Bdhn*6(1) *Bdhn*7(2) *Bdhn*8(1)<br>**BS:** *Bdhn*1(1) *Bdhn*2(2) *Bdhn*5(3) *Bdhn*6(1) *Bdhn*7(2) *Bdhn*8(1)<br>**BSY:** *Bdhn*2(1) *Bdhn*5(1) *Bdhn*7(1) *Bdhn*8(2) *Bdhn*9(1) |
| Myb | MYB124 | 0.598 | 2.18 |  | **BD:** *Bdhn*1(5) *Bdhn*2(4) *Bdhn*3(2) *Bdhn*6(1) *Bdhn*7(2) *Bdhn*8(2)<br>**BHD:** *Bdhn*1(7) *Bdhn*2(4) *Bdhn*3(2) *Bdhn*6(1) *Bdhn*7(2) *Bdhn*8(2)<br>**BHS:** *Bdhn*1(7) *Bdhn*2(4) *Bdhn*3(2) *Bdhn*6(1) *Bdhn*8(2)<br>**BS:** *Bdhn*1(8) *Bdhn*2(4) *Bdhn*3(2) *Bdhn*6(1) *Bdhn*8(2)<br>**BSY:** *Bdhn*1a(4) *Bdhn*1b(4) *Bdhn*2(4) *Bdhn*3(2) *Bdhn*6(1) *Bdhn*7(2) |
| C2H2 zinc finger | ZAT | 0.738 | 4.91 |  | **BD:** *Bdhn*4(3) *Bdhn*5(1) *Bdhn*10(2)<br>**BHD:** *Bdhn*5(1)<br>**BHS:** *Bdhn*5(1) *Bdhn*10(1)<br>**BS:** *Bdhn*5(2) *Bdhn*10(1) |

### The *Brachypodium* dehydrin tree

A large ML *Brachypodium* dehydrin tree was constructed from a data set of 47 *Bdhn* protein coding regions present in the four studied *Brachypodium* species and in five outgroup grasses (Fig. 3). Most of the major splits were highly to fully supported (80-100% BS) and only few were moderately supported (BS>60%). The most divergent split separated the duplicated *Bdhn*1-*Bdhn*2 (ERD14) clade from the rest, followed by the isolated *Bdhn*10 (HIRD11) clade. Within the remaining group of Y_n_SK_n_ dehydrin structural genes, there was a divergence of the *Bdhn*9 (XEROI) clade, followed by subsequent divergences of the Y_n_SK_n_ *Bdhn*4-*Bdhn*5, *Bdhn*6, *Bdhn*3 and *Bdhn*7/*Bdhn*8 clades (Fig. 3). All clades included *Brachypodium* and outgroup sequences except *Bdhn*7/*Bdhn*8. *Brachypodium* was resolved as monophyletic in all *Bdhn* clades except *Bdhn*10 in which *H. vulgare* DHNHvul13 was nested within but with low internal support. In all *Bdhn* clades, dehydrin sequences from the *B. hybridum* D and S subgenomes were resolved as sister to those of its diploid progenitor *B. distachyon* and *B. stacei* species, respectively. In six *Bdhn* clades (*Bdhn*1, *Bdhn*3, *Bdhn*(4)5, *Bdhn*7, *Bdhn*8, *Bdhn*9) the *B. stacei*/ *B. hybridum-*S sequences diverged earlier than the ((*B. distachyon*/ *B. hybridum-*D)/ *B. sylvaticum*) sequences (Fig 3), whereas the other clades showed different resolutions.

**Figure 3.**
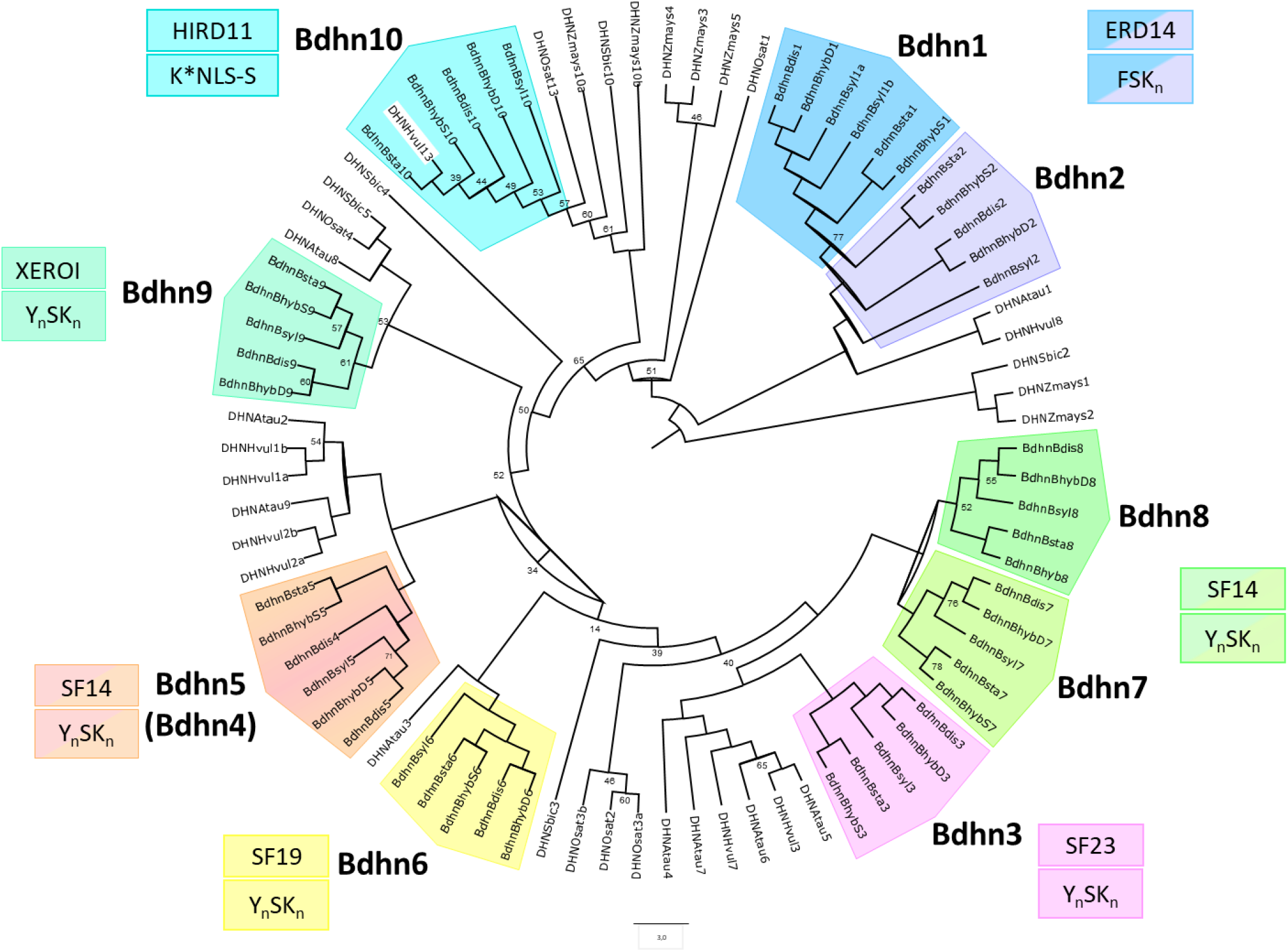
Maximum Likelihood *Brachypodium Bdhn* tree. Unrooted IQtree cladogram showing the relationships among the dehydrin *Bdhn* gene clades and orthologous grass sequences and among *Brachypodium* species and genomes within each clade. *Bdhn* clades are identified by colors, *Bdhn* codes, gene architecture, and Panther subfamily codes (see Table 1). Duplicated *Bdhn* genes form sister clades or fall within the same clade. Ultrafast Bootstrap support (<80%) is shown on branches; the remaining branches are fully supported.

### Chromosome distributions and selection analysis of duplicated *Bdhn* genes

Dehydrin genes were distributed among the five chromosomes of *B. distachyon* and the D subgenome of *B. hybridum* (BhD), in 6 out of 10 chromosomes of *B. stacei* and the S subgenome of *B. hybridum* (BhS), and in 6 out of 9 chromosomes of *B. sylvaticum* (Bsy) (Fig. 4, Supplementary Table 5). The highest density of dehydrin genes were found in the chromosomes Bd3 and Bd4 (*Bdhn*2, *Bdhn*4, *Bdhn*5, *Bdhn*6, *Bdhn*7, *Bdhn*8), Bs4 (*Bdhn*2, *Bdhn*6, *Bdhn*7), the equivalent *B. hybridum* D and S subgenomic chromosomes (except *Bdhn*4), and Bsy4 (*Bdhn*2, *Bdhn*6, *Bdhn*7, *Bdhn*8). In *B. distachyon* and *B. hybridum* D subgenome our analysis detected tandem duplications of *Bdhn*7-*Bdhn*8 in Bd3 and of *Bdhn*4-*Bdhn*5 in Bd4 (only *B. distachyon*), and a segmental duplication of *Bdhn*1-*Bdhn*2 in Bd3 and Bd5 (Fig. 4, Supplementary Table 5). *B. stacei* and *B. hybridum* S subgenome and *B. sylvaticum* showed a tandem duplication of *Bdhn*7-*Bdhn*8 in Bs4 and Bsy4, and a segmental duplication of *Bdhn*1-*Bdhn*2 in Bs4 and Bs9 and in Bsy4 and Bsy9, respectively (Fig. 4). In addition, *B. sylvaticum* showed a tandem duplication of *Bdhn*1a and *Bdhn*1b in Bsy9 not found in the other *Brachypodium* species studied (Fig. 4, Supplementary Table 5). Selection rates values of non-synonymous vs synonymous substitutions (dN/dS) gave values of ω <1 in all *Bdhn* genes and species, and for all duplicated paralogs in each species (Supplementary Tables 6a,b).

**Figure 4.**
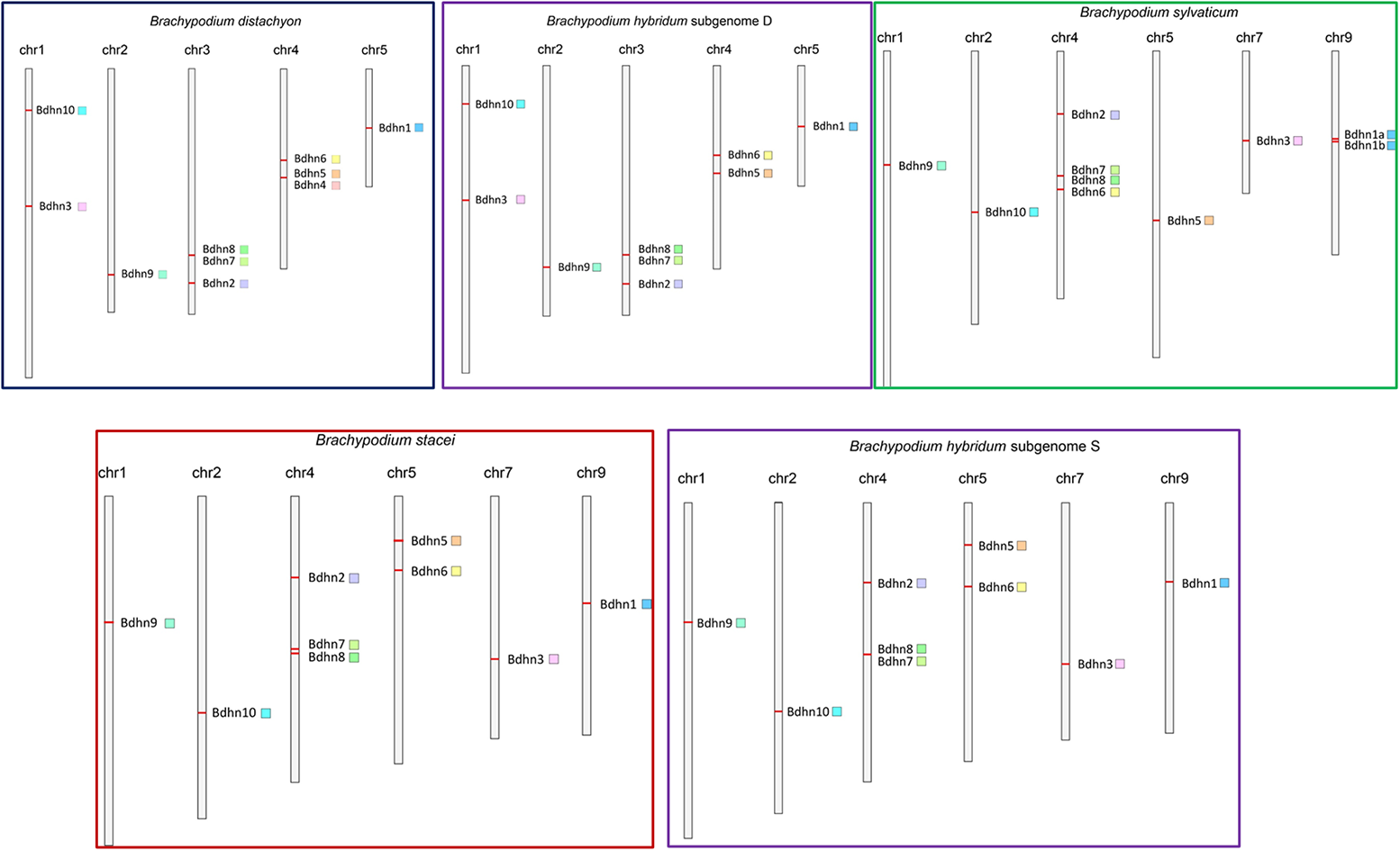
Chromosomal location of *Bdhn* genes in the studied *Brachypodium* species. *B.stacei* (red rectangle), A. *distachyon* (blue), *B. hybridum* subgenomes S and D (purple) and *B. sylvaticum* (green). *Bdhn* genes are mapped on the chromosome with their respective color flags (see Figure 2).

### Dehydrin gene clusters, phylogenetics, and climate niche variation in *B. distachyon*

The genomes of the 54 *B. distachyon* ecotypes contained in most cases (74.07%) all ten *Bdhn* genes (Supplementary Fig. S3). The few genotypes with fewer or more than 10 genes were those with poor genome assemblies (e. g., Bd2-3, Gaz8) and thus likely prone to artefactual inferences of gene losses or gains. Independent ML trees obtained for each of the ten *Bdhn* genes constructed from data sets with exon and intron sequences showed differently resolved topologies (Supplementary Fig. S4). Six of those genes (*Bdhn*1, *Bdhn*2, *Bdhn*3, *Bdhn*6, *Bdhn*7, *Bdhn*8) recovered a congruent topology for some *B. distachyon* groups. A ML tree constructed from their concatenated sequences produced a combined *B. distachyon Bdhn* tree showing a relatively well-supported EDF+ clade (74% BS), and the successive though weakly supported divergences of the S+ and T+ lineages (Supplementary Fig. S5a). A clade of T+ lineages (Bd18-1, Bd21-3, BdTR5i) was resolved as sister to the EDF+ clade. Topological congruence RH and SH tests performed between the *B. distachyon* nuclear tree of Gordon et al., (2017) (Supplementary Fig. S5b), the *B. distachyon* plastome tree of Sancho et al. (2018) (Supplementary Fig. S5c), and the *B. distachyon Bdhn* tree indicated that the topology of the *Bdhn* tree was significantly similar (p<0.001) to the topologies of the nuclear and plastome trees (Supplementary Table 7), however the *Bdhn* tree resembled more the plastome tree than the nuclear tree (Supplementary Fig. S5).

The optimal climate niche of each *B. distachyon* ecotype was inferred from PCA of 19 climate variables (Supplementary Table S3; Supplementary Fig. S6). The main PC1 axis (51.8% of variance) was defined by precipitation (driest quartet and month, warmest quartet) and temperature (annual range, maximum of warmest month) variables. PC1 coordinate values allowed us to classify the *B. distachyon* ecotypes into cold (<-3; ABR2, ABR3, ABR4, ABR5, ABR6, RON2), warm (>3; Bd2-3, Bd21, Bd3-1, Bis1, Kah1, Kah5, Koz1, Koz3) and mesic (−3 to 3, remaining) climate class ecotypes.

### Differential expression of *Bdhn* genes in *Brachypodium distachyon* ecotypes under drought and temperature stress conditions

Only four out of the ten identified *Bdhn* genes were expressed in mature leaves of all 32 studied *B. distachyon* ecotypes [*Bdhn*1a (Bradi5g10860.1), *Bdhn*2 (Bradi3g51200.1), *Bdhn*3 (Bradi1g37410.1), *Bdhn*7 (Bradi3g43870.1); Supplementary Table S2]. These annotated dehydrins showed significant differential expression (DE) levels between the watered (W) and dry (D) conditions, independently of temperature conditions in both separate CD-CW-HD-HW and averaged D vs W comparative tests (Table 4; Supplementary Fig. S7a, b). By contrast, the dehydrin expressions did not show significant differences between cool (C) and hot (H) conditions under drought treatment, and only *Bdhn*3 and *Bdhn*7 showed significant differences in CW vs HW conditions, though none of them did in averaged C vs H comparative tests (Supplementary Fig. S7a, c).

**Table 4.** Summary statistics of dehydrin *Bdhn*1a, *Bdhn*2, *Bdhn*3 and *Bdhn*7 gene expressions under dry (D) vs watered (W) conditions and comparative differential expression (DE) tests in *B. distachyon* ecotypes. (a) Kruskal-Wallis rank tests (D vs W) for each *Bdhn* gene. (b) Wilcoxon pairwise tests of normalized TPM values across ecotypes, p-values were adjusted with the Benjamini–Hochberg (BH) procedure, controlling the false discovery rate, to correct for multiple comparisons; n. s., non significant, *p≤ 0.05*; significant values are highlighted in bold.

| Var | *Bdhn1a* | *Bdhn2* | *Bdhn3* | *Bdhn7* |
| --- | --- | --- | --- | --- |
| t-test | 123.74 | 179.98 | 178.8 | 175.02 |
| df | 63 | 63 | 63 | 63 |
| p-value | 7.69E-06 | 3.38E-13 | 5.10E-15 | 1.75E-12 |

(b)
| Ecotype | *Bdhn1a* |  |  | *Bdhn2* |  |  | *Bdhn3* |  |  | *Bdhn7* |  |  |
| --- | --- | --- | --- | --- | --- | --- | --- | --- | --- | --- | --- | --- |
|  | D | W | W-test | D | W | W-test | D | W | W-test | D | W | W-test |
| **ABR2** | 214.175 | 159.875 | n.s | **167.1** | **46.05** | * | **172.425** | **18.65** | * | **19.4** | **2.3** | * |
| **ABR3** | 263.775 | 215.833 | n.s | 169.05 | 48.467 | n.s | 219.875 | 15.167 | n.s | 29.525 | 2.3 | n.s |
| **ABR4** | 247.15 | 269.375 | n.s | 108.425 | 48.4 | n.s | **62.175** | **5.025** | * | 15.1 | 2.4 | n.s |
| **ABR5** | 252.633 | 272.233 | n.s | 311.233 | 37.6 | n.s | 294.767 | 6.967 | n.s | 51.6 | 2.967 | n.s |
| **ABR6** | 290.175 | 302.233 | n.s | **204.525** | **23.433** | * | 850.55 | 3.8 | n.s | 41.55 | 3 | n.s |
| **ABR8** | 530.05 | 255.333 | n.s | 359.425 | 31.767 | n.s | 231.05 | 4.367 | n.s | 127.775 | 1.333 | n.s |
| **Adi10** | 659.4 | 532.767 | n.s | 935.575 | 40.133 | n.s | 1216.925 | 8.4 | n.s | 332.9 | 4.167 | n.s |
| **Adi12** | 488.65 | 418.675 | n.s | **591.725** | **68.75** | * | **498.525** | **11.75** | * | **139.825** | **4.45** | * |
| **Adi2** | 461.1 | 430 | n.s | 511 | 56.733 | n.s | 437.6 | 3.367 | n.s | 63.567 | 3 | n.s |
| **Bd1_1** | 414.3 | 355.45 | n.s | 152.35 | 32.25 | n.s | 45.95 | 11.25 | n.s | 14.95 | 2.75 | n.s |
| **Bd18-1** | 736.95 | 259.3 | n.s | 548.1 | 19 | n.s | 410.6 | 3.9 | n.s | 172.9 | 0.45 | n.s |
| **Bd21-3** | 632.767 | 290.767 | n.s | 890.1 | 25.8 | n.s | 839.567 | 14.3 | n.s | 360.667 | 1.5 | n.s |
| **Bd21ctrl** | 326.85 | 300.1 | n.s | 113.725 | 44.7 | n.s | 30.575 | 16.375 | n.s | 24.325 | 3.35 | n.s |
| **Bd2-3** | 502.175 | 368.35 | n.s | 202.05 | 55.675 | n.s | 94.825 | 3.925 | n.s | 29.275 | 2.5 | n.s |
| **Bd30-1** | 488.675 | 365.05 | n.s | **761.7** | **33.575** | * | **906.55** | **5.725** | * | **222.7** | **2.3** | * |
| **Bd3-1** | 395.167 | 361.967 | n.s | 465.367 | 41.367 | n.s | 1711.533 | 9.367 | n.s | 329.433 | 1.233 | n.s |
| **BdTR10c** | 410.375 | 200.2 | n.s | 327.65 | 43 | n.s | 924.275 | 4.2 | n.s | 194.1 | 1.75 | n.s |
| **BdTR11g** | 497.15 | 388.925 | n.s | **454.7** | **59.35** | * | **643.075** | **17.6** | * | **95.65** | **7.2** | * |
| **BdTR11i** | 531.875 | 325.433 | n.s | 553.2 | 43.533 | n.s | 918.9 | 5.033 | n.s | 124.325 | 3.167 | n.s |
| **BdTR13a** | 697.567 | 298.45 | n.s | 434.5 | 44.95 | n.s | 313.733 | 7.4 | n.s | **114.633** | **1.6** | * |
| **BdTr1i** | 582.925 | 351.925 | n.s | **640.25** | **46.1** | * | **727.325** | **5.7** | * | **251.15** | **2.425** | * |
| **BdTR2b** | 538.175 | 318.55 | n.s | 405.55 | 43.85 | n.s | 436.05 | 7.2 | n.s | 136.4 | 1.65 | n.s |
| **BdTR2g** | 750.55 | 451.6 | n.s | **816.325** | **42.775** | * | **1517.95** | **5.875** | * | **517.675** | **3.6** | * |
| **BdTR3c** | 659.525 | 317.5 | n.s | 701.275 | 67.833 | n.s | 1032.85 | 57.633 | n.s | 194.4 | 19.533 | n.s |
| **BdTR5i** | 364.233 | 277.933 | n.s | 352.133 | 36.733 | n.s | 440.233 | 6.367 | n.s | 77.333 | 2.367 | n.s |
| **BdTR9k** | 436.75 | 197.433 | n.s | 261.225 | 24.533 | n.s | 220.15 | 2.667 | n.s | 78.375 | 1.033 | n.s |
| **Bis1** | 450.233 | 364.067 | n.s | 391.6 | 37.833 | n.s | 541.133 | 6.067 | n.s | 155.1 | 1.133 | n.s |
| **Kah1** | 385.4 | 202.367 | n.s | 392.325 | 54.333 | n.s | 405.325 | 7.7 | n.s | 65.9 | 1.833 | n.s |
| **Kah5** | **432.775** | **303.925** | * | **610.7** | **55.275** | * | **481.95** | **11.675** | * | **86.375** | **3.325** | * |
| **Koz1** | 481.85 | 388.6 | n.s | 500.6 | 63.833 | n.s | 257.7 | 15.2 | n.s | 55.9 | 4.7 | n.s |
| **Koz3** | **898.525** | **344.1** | * | **1074.025** | **28.825** | * | **3441.075** | **6.975** | * | **812.2** | **1.775** | * |
| **RON2** | 656.167 | 288.3 | n.s | 789.467 | 65.933 | n.s | 558.7 | 31.833 | n.s | 140.433 | 12.033 | n.s |

The four dehydrins showed significantly increased expression levels in drought conditions in most accessions (Wilcoxon tests, Table 4; Tukey tests, Fig. 5, Supplementary Fig. S8). The DE levels were also significantly different among ecotypes, especially within the dry treatment, being highest in warm ecotypes Koz3 (*Bdhn*1a, *Bdhn*3, *Bdhn*7) and Adi10 (*Bdhn*2) and lowest in cold ecotypes ABR2 (*Bdhn*1) and ABR4 (*Bdhn*2, *Bdhn*3, *Bdhn*7) (Fig. 5, Supplementary Fig. S8). On average, across all ecotypes, drought increased dehydrin expression by about 5,74% in *Bdhn*1a, 39% in *Bdhn*2, 67,8% in *Bdhn*3 and 97,8% in *Bdhn*7 compared to watered plants (Supplementary Fig. S7b). Overexpression of dehydrins caused by drought stress was significantly correlated between all *Bdhn* gene pairs (Supplementary Table 8; Supplementary Fig. S9).

**Figure 5.**
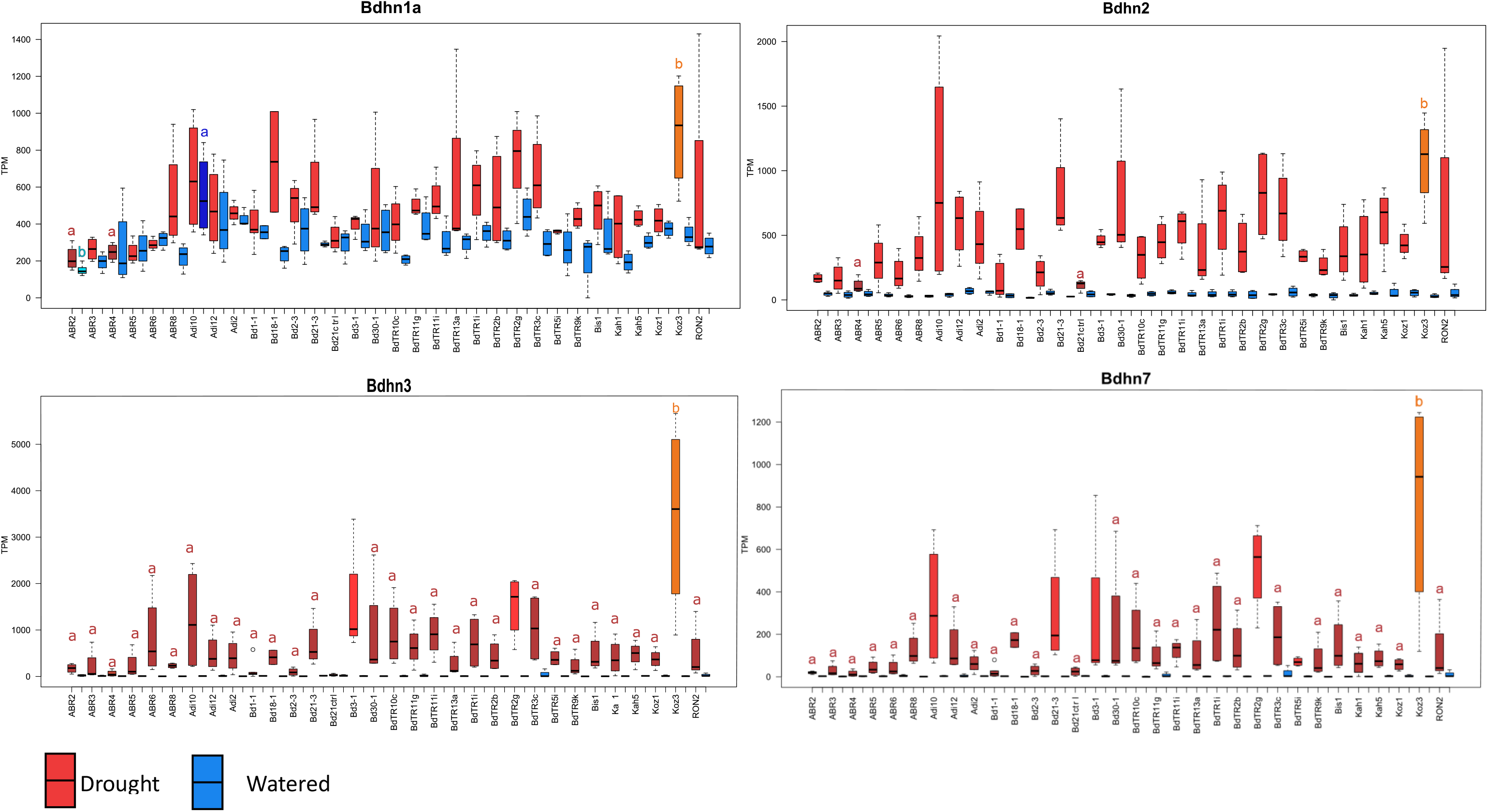
Differentially expressed *Bdhn*1a, *Bdhn*2, *Bdhn*3 and *Bdhn*7 dehydrin genes (transcript per million, TPM) in 32 ecotypes of *B. distachyon* under drought (D, red) vs watered (W, blue) conditions. Different letters in the boxplots indicate significant group differences (Tukey tests) (see also Supplementary Fig. S8).

Our *B. distachyon* - *T. aestivum* DE comparative analysis found that the drought-induced *Bdhn*1, *Bdhn*2, *Bdhn*3, and *Bdhn*7 genes belong to the same ortholog groups as 15 out of the 16 differentially expressed wheat dehydrin (DHN) genes (Supplementary Table 9) under natural field drought stress (Galvez et al 2019) or greenhouse imposed drought stress (Reddy et al. 2014, Chu et al. 2021). In wheat there is a physical clustering of several dehydrin genes in two gene clusters located in the 5L and 6L groups of wheat chromosomes (Supplementary Table 9; Supplementary Fig. S10). The 6L cluster contains 25 dehydrins and includes nine DHN3 genes and three DHN4 genes (all ortholog to *Bdhn*3), whereas the 5L cluster has 13 DHN38 genes of which six are orthologs to *Bdhn*7. In addition, the DHN11 genes, located in another portion of 6L chromosomes, are orthologs to the *Bdhn*1 and *Bdhn*2 genes. We observed that orthologs from *B. distachyon* and *T. aestivum* tended to show a similar pattern of expression response to soil drying. Specifically, the duplicated Bdhn1 and Bdhn2 genes and the DHN11(A1) gene were all up-regulated in drought treatments, as were the Bdhn7 gene and the duplicated DHN38 (B1, B2) genes and the Bdhn3 gene and the duplicated DHN3 (A1, A6, B6, D1, D4, D6, D8, D9) genes (Supplementary Table 9; Supplementary Fig. S10; Galvez et al. 2019).

### Effects of drought on dehydrin gene expression and drought-response phenotypic traits

The 12 drought-response phenotypic traits studied also showed significant different values in dry vs watered conditions across ecotypes (Supplementary Table 10). On average, drought significantly decreased the values of six traits (17.14% in abvgrd, 34.78% in ttlmass, 4% in leaf_rwc, 36.5% in leaf_wc, 12.5% in leafn, 2.8% in WUE) and significantly increased those of five traits (33% in pro, 5.5% in rmr, 2.96% in leafc, 5.71% in lma, 21.4% in cn) compared to watered conditions, but did not significantly affect the blwgr trait (Supplementary Fig. S11), as shown previously (Des Marais et al. 2017).

Drought-induced effects caused significant positive and negative correlations between the averaged expressed values of the four *Bdhn* genes and changes in most phenotypic trait values (Supplementary Table 11; Supplementary fig. S12). Regression models for independent correlations between the *Bdhn*1a, *Bdhn*2, *Bdhn*3 and *Bdhn*7 expressions and the changes in the 12 phenotypic traits showed significant positive correlations for most dehydrin *Bdhn* genes with pro, blwgrd, rmr, WUE, leafc and c:n, negative correlations with leaf_rwc, leaf_wc and leafn, and non-significant correlations with ttlmass and abvgrd (Supplementary fig. S13).

### Phylogenetic signal in *Brachypodium distachyon*

The potential phylogenetic signal of dehydrin expression, phenotypic trait changes, and climate variation was evaluated on both the *B. distachyon* nuclear species tree (Gordon et al., 2017) and the *B. distachyon* dehydrin *Bdhn* tree. Few traits showed significant phylogenetic signal when tested on the *B. distachyon* nuclear species tree, and most of their K and lambda values were low (Supplementary Table 12; Supplementary Fig. S13); by contrast, a larger number of traits showed significant phylogenetic signal and higher K and lambda values when tested on the *B. distachyon Bdhn* tree (Table 5; Fig. 6). None of the dehydrin gene expression values under W or D had significant K or lambda values on the *B. distachyon* species tree (Supplementary Table 12a; Supplementary Fig. S13a); however, *Bdhn*2W (K=0.34, p<0.05; lambda=0.75, p<0.01), *Bdhn*7W (K=0.43, p<0.05), and *Bdhn*3D (lambda=0.95, p<0.05; K=0.43, p=0.06) carried phylogenetic signal when tested on the *B. distachyon Bdhn* tree (Table 5a; Fig. 6a). Similarly, the phenotypic rmrW (K=0.67, p<0.01; lambda=1.00, p<0.001), leafnW (K=0.31, p<0.05; lambda=0.70, p<0.05), cnW (K=0.29, p<0.05), abwgrW (lambda=0.58, p<0.05), leafcD (K=0.40. p<0.01), leafnD (lambda=0.74, p<0.001), and cnD (lambda=0.74, p<0.001) traits’ values carried phylogenetic signal, and leaf_rwcD (K=0.29, p=0.06), and abvgdD (lambda=0.43, p=0.05) marginal phylogenetic signal on the *Bdhn* tree (Table 5b; Fig. 6b). The climate niche of ecotypes showed significant phylogenetic signal on both trees, but the signal was higher (K=0.46, P<0.01; lambda=0.75, p<0.01) on the *Bdhn* tree (Table 5c, Fig. 6c) than in the nuclear species tree (Supplementary Table 12c; Supplementary Fig. S13c).

**Figure 6.**
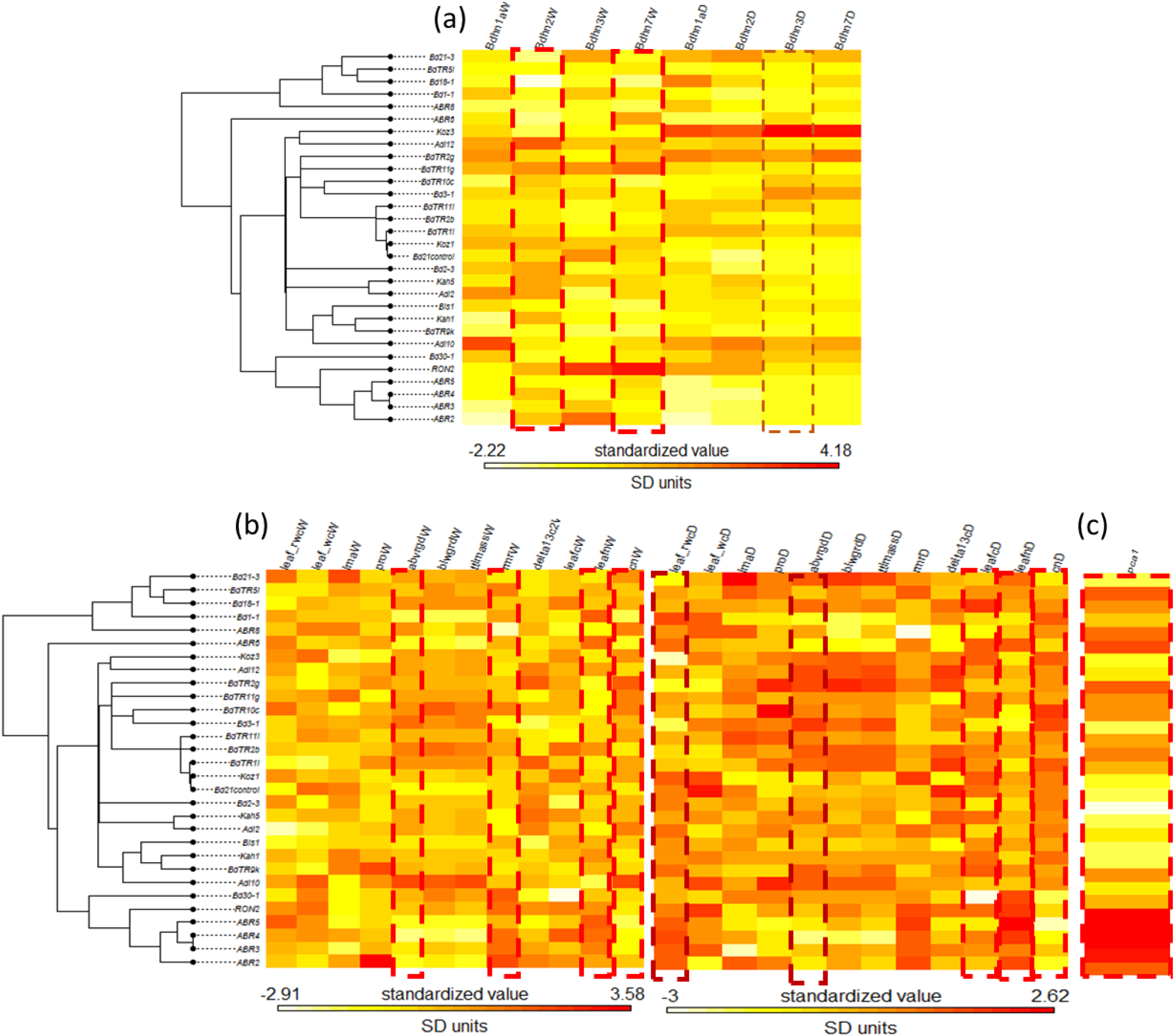
Maximum Likelihood *B. distachyon Bdhn* tree cladogram showing the relationships of 30 ecotypes. Phyloheatmaps of normalized values for different sets of variables: (a) dehydrin (*Bdhn*1, *Bdhn*2, *Bdhn*3, *Bdhn*7) gene expression values under watered (W) and drought (D) conditions; (b) drought-response phenotypic traits (leaf_rwc; leaf_wc; lma; pro; abvrgd; blwgrd; ttlmass; rmr; delta13c; leafc; leafn; cn) values under watered (W) and drought (D) conditions; (c) climate niche PC1 values. Traits showing significant phylogenetic signal are highlighted with dotted lines (see Table 5).

**Table 5.** Phylogenetic signal of **(a)** dehydrin gene expressions under watered (W) and dry (D) conditions, **(b)** drought-induced phenotypic traits changes and **(c)** climate niche variation assessed in the *B. distachyon Bdhn* tree using the *phylosig* option of the *phytools* R package. Blomberg’s K and Pagel’s lambda values close to one indicate phylogenetic signal and values close to zero phylogenetic independence. K, p-values based on 1000 randomizations; lambda, p-values based on the Likelihood Ratio test. Significant and marginal significant values are highlighted in bold.

| *Bdhn* gene | Treatment | K | p-value | lambda($\lambda$) | logL($\lambda$) | LR($\lambda=0$) | p-value |
| --- | --- | --- | --- | --- | --- | --- | --- |
| *Bdhn1aW* | W | 0.231879 | 0.142 | 0.782475 | -175.48 | 1.121 | 0.289703 |
| ***Bdhn2W*** |  | **0.34558** | **0.017** | **0.759422** | **-112.036** | **6.78197** | **0.00920834** |
| *Bdhn3W* |  | 0.10957 | 0.603 | 6.64306E-05 | -93.2539 | 0.00063134 | 1 |
| ***Bdhn7W*** |  | **0.433394** | **0.031** | **0.97584** | **-60.2457** | **3.42969** | **0.0640339** |
| *Bdhn1aD* | D | 0.308794 | 0.051 | 0.502054 | -194.065 | 0.710376 | 0.399319 |
| *Bdhn2D* |  | 0.211919 | 0.176 | 6.64306E-05 | -208.371 | 0.00063094 | 1 |
| ***Bdhn3D*** |  | **0.433425** | **0.064** | **0.951732** | **-234.916** | **4.48918** | **0.0341101** |
| *Bdhn7D* |  | 0.379302 | 0.083 | 0.917646 | -195.278 | 1.74672 | 0.18629 |

(b)
| Trait | K | p-value | lambda($\lambda$) | logL($\lambda$) | LR($\lambda=0$) | p-value |
| --- | --- | --- | --- | --- | --- | --- |
| leaf_rwcW | 0.00633355 | 0.137 | 6.61E-05 | -21.1366 | -21.1361 | 1 |
| leaf_wcW | 0.00601205 | 0.142 | 0.712778 | -124.3118 | -124.641 | 0.4171331 |
| lmaW | 0.0127346 | 0.067 | 0.5672695 | -53.1973 | -54.02652 | 0.1978138 |
| proW | 0.00751385 | 0.279 | 6.61E-05 | -65.7397 | -65.73915 | 1 |
| **abvrgdW** | **0.00982265** | **0.118** | **0.9056281** | **-143.4905** | **-147.2048** | **0.006419007** |
| blwgrdW | 0.00503958 | 0.192 | 0.372553 | -117.8161 | -117.602 | 1 |
| ttlmassW | 0.00760606 | 0.15 | 0.8313881 | -153.3088 | -155.4348 | 0.0392068 |
| **rmrW** | **0.0198041** | **0.031** | **0.8500612** | **-65.57032** | **-69.23347** | **0.006795293** |
| WUE_W | 0.00079528 | 0.921 | 6.61E-05 | -9.565582 | -9.565045 | 1 |
| leafcW | 0.00344133 | 0.413 | 6.61E-05 | -84.79111 | -84.79067 | 1 |
| **leafnW** | **0.00111356** | **0.809** | **0.6412875** | **-71.1111** | **-72.40537** | **0.1076397** |
| **cnW** | **0.00101427** | **0.842** | **6.15E-01** | **-35.11085** | **-36.01121** | **0.1796235** |
| **leaf_rwcD** | **0.00595303** | **0.202** | **0.8240433** | **-70.83287** | **-71.94764** | **0.1353942** |
| leaf_wcD | 0.0191719 | 0.029 | 0.5318181 | -129.0792 | -129.2025 | 0.6194317 |
| lmaD | 0.00655413 | 0.198 | 0.4014225 | -60.3207 | -60.04825 | 1 |
| proD | 0.00178069 | 0.645 | 6.61E-05 | -118.2053 | -118.2048 | 1 |
| **abvrgdD** | **0.0548587** | **0.005** | **0.9862024** | **-132.3802** | **-137.5324** | **0.001327115** |
| blwgrdD | 0.00799846 | 0.135 | 0.6778591 | -112.6454 | -113.232 | 0.2787568 |
| ttlmassD | 0.0254584 | 0.02 | 0.9586421 | -144.0437 | -147.1176 | 0.01315614 |
| rmrD | 0.0117638 | 0.1 | 0.8587116 | -71.60054 | -74.00027 | 0.02846858 |
| WUE_D | 0.0206935 | 0.023 | 0.5455282 | -9.438273 | -9.102205 | 1 |
| **leafcD** | **0.00564166** | **0.206** | **6.61E-05** | **-86.81046** | **-86.80988** | **1** |
| **leafnD** | **0.0127895** | **0.066** | **0.8379002** | **-52.22653** | **-57.04403** | **0.001909058** |
| **cnD** | **0.015233** | **0.051** | **0.8350859** | **-24.49639** | **-28.06983** | **0.007509447** |

(c)
| Trait | K | p-value | lambda($\Lambda$) | logL($\Lambda$) | LR($\Lambda=0$) | p-value |
| --- | --- | --- | --- | --- | --- | --- |
| PCA1 | 0.462712 | 0.005 | 0.753963 | -72.2802 | 9.38246 | 0.00219071 |

## Discussion

### The dehydrin family in *Brachypodium*: evolution and functionality

Our comparative genomic analysis of the dehydrin genes in the reference genomes of four *Brachypodium* species and in the genomes of 54 ecotypes of *B. distachyon* has allowed us to identify 47 *Bdhn* genes. These are almost twice the number of LEA2 genes previously found in *B. distachyon* Bd21 (Filiz et al., 2013) and a number of DHNs proportional to those found in diploid rice species (Verma et al., 2017) though less abundant than those detected in allohexaploid wheat (Wang et al., 2014). All the dehydrin types but *Bdhn*4 are present in all the studied species of *Brachypodium* (Table 1; Figs. 1, 3), suggesting that they were inherited from their common ancestor. Orthology and evolutionary analysis indicate that most of these proteins were probably present in the ancestor of the grasses (Supplementary Table 4; Fig. 3). The *Bdhn*1 and *Bdhn*2 (ERD14) genes, segmentally duplicated in *Brachypodium*, and *Bdhn*10 (HIRD11) encode the most divergent dehydrins (Fig. 3), whereas the remaining Y_n_SK_n_ genes (*Bdhn*9, *Bdhn*4-5, *Bdhn*6, *Bdhn*3, *Bdhn*7/*Bdhn*8) have apparently evolved more recently in the grass tree. The tandemly duplicated *Bdhn*7 and *Bdhn*8 genes, only found in *Brachypodium* species, were likely inherited from their common ancestor (Fig. 3). *Bdhn*7 and *Bdhn*8 proteins probably evolved, in turn, from a duplication of an ancestral *Bdhn*3 gene through insertion/deletions (*Bdhn*7-8) and the loss one K-segment (*Bdhn*8) (Figs. 1b, 3).

Segmental and tandem duplications of *Bdhn*1-*Bdhn*2 and *Bdhn7*-*Bdhn*8 genes have been detected in all studied *Brachypodium* species (Figs. 3, 4; Supplementary Fig. S3); however, *Bdhn*4 is inferred to have originated from a tandem duplication of *Bdhn*5 exclusively in the ancestor of the *B. distachyon* lineage and *Bdhn*1b from a tandem duplication of *Bdhn*1a in the *B. sylvaticum* lineage (Supplementary Table 5; Fig. 3; Supplementary Fig. S3). The allotetraploid *B. hybridum* exhibits homeologous copies inherited from its diploid progenitor species for the same sets of tandemly and segmentally duplicated genes (Table 1; Figs. 3, 4). Nonetheless, the loss of the *Bdhn*4 gene in its D subgenome probably occurred after the hybridization and whole genome duplication (WGD) event that originated this reference genome (Catalán et al., 2012; Gordon et al., 2020), as this gene is largely present in the *B. distachyon* ecotypes studied (Supplementary Table 5; Supplementary Fig. S3). Our data support the hypothesis of a highly dynamic evolution of duplications and losses of dehydrin paralogs in *Brachypodium*. This was also evidenced in other grasses such as *Hordeum vulgare* (Choi et al. 1999), *Triticum aestivum* (Wang et al., 2014), and some *Oryza* species (Verma et al., 2017). Extrapolations from inferred ages of *Brachypodium* lineages depict a scenario of rapid turnover rates of *Bdhn* duplications and gene losses within the last ten (*Brachypodium* crown node split) to half million (*B. distachyon* – *B. hybridum* split) years (Sancho et al., 2018; Gordon et al., 2020).

The consensus ML phylogenetic tree of *Brachypodium* species based on ten *Bdhn* genes (Fig. 3) showed a strong support for most clades and a congruent topology in seven gene clades resulting in more ancestral *B. stacei* sequences followed by the split of more recent *B. distachyon* and *B. sylvaticum* copies. This resolution was fully congruent with that of the *Brachypodium* species tree (Díaz-Pérez et al., 2018); the branch swaps observed in the three remaining *Bdhn*2, *Bdhn*6, and *Bdhn*10 clades (Fig. 3) likely resulted from incomplete lineage-sorting events. Therefore, the evolution of the dehydrin genes was in pace with the organismal evolution of the *Brachypodium* lineages, supporting their species-level evolutionary synchrony. By contrast, the *Bdhn* intraspecific phylogenies of *B. distachyon* ecotypes were more variable or unresolved (Supplementary Fig. S4), and the combined *B. distachyon Bdhn* tree (Supplementary Fig. S5a) showed a congruent topology with respect to those of the *B. distachyon* nuclear species tree and plastome tree but more similar to the later (Supplementary Table 7; Supplementary Figs. S5). These results indicate that the conserved dehydrin genes track the fast divergences of the recent *B. distachyon* ecotypes, recovering a relatively supported EDF+ clade and a lineage divergence order concordant with that of the plastome tree (Supplementary Fig. S5a, c).

Selection analysis of *Bdhn* genes have consistently detected ɷ<1 values for all ten genes in the studied species, including the duplicated *Bdhn* genes (Supplementary Tables 6a, b). Our results indicate that all *Brachypodium* dehydrins are probably functional and that the duplicated paralogs are under selective constraint, irrespective of their ancestral or recent origins (Fig. 3; Supplementary Fig. S4). Paralogs may experience pseudogenization or neofunctionalization events after their duplication (Riley et al., 2019); however, in contrast to the pseudogenization trend observed in other duplicated genes of *Brachypodium* (e. g., serpin genes; Rehman et al. 2020), no dehydrin pseudogenes have been detected in our *Brachypodium* genomic search, although dehydrin pseudogenes have been recorded in other angiosperms (e. g., *Populus*, Liu et al., 2017; *Arabidopsis*, Riley et al., 2019; Yu et al., 2015).

The amino acid composition, structure, and biochemical features inferred for the *Brachypodium* dehydrins (Table 2) support their potential roles as regulators of the water-deficit in the cells (Graether & Boddington, 2014; Riley et al., 2019; Verma et al., 2017). *Bdhn* dehydrins variation in size, molecular weight, pI and GRAVY values falls within those observed for rice dehydrins (Verma et al., 2017) The large differences in *Bdhn* pI values [≤6 (*Bdhn*1, 2, 9) – >9 (*Bdhn*3, 4, 5, 6, 7, 8) suggest that those proteins may be located in specific compartments of the cell, like the cytoplasm and the nucleus. *Bdhn* dehydrins with high pI values and with phosphorylated S-segments (Y_n_SK_n_; e. g., *Bdhn*3, *Bdhn* 4, *Bdhn*5, *Bdhn* 6, *Bdhn*7, *Bdhn*8) may bind negatively charged molecules such as DNA (Graether & Boddington, 2014). The Bdh10 dehydrins (HIRD11 family), which lack the K-segment, could bind different ions and reduce the formation of reactive oxygen species (ROS), like that observed for the AtHIRD11 ortholog (Hara et al., 2013). The three-domain complex architecture of *Bdhn*9 (DnaJ-DnaJX-DHN) has been also observed in other grass dehydrins, like DHN1 of rice (Verma et al., 2017) and *Setaria italica* (Jiménez-Bremont et al., 2013); the DnaJ domain may have a chaperone function (Verma et al., 2017).

### Dehydrin expression induction in Brachypodium distachyon

Dehydrin expression varies considerably both in plant tissue and developmental stage and under different abiotic stress conditions (Graether & Boddington, 2014; Verma et al., 2017; Wang et al., 2014). Most DHN genes have shown high expression profiles in seeds and immature seedling stages in wheat and rice (Verma et al., 2017; Wang et al., 2014); however, their expression decreases in mature or late-development tissues or is even absent in some organs, like mature leaves in rice (Verma et al., 2017). Our expression analyses demonstrated that only four (*Bdhn*1a, *Bdhn*2, *Bdhn*3, *Bdhn*7) out of the ten detected *Brachypodium Bdhn* dehydrin genes were constitutively and inductively expressed in mature leaves of *B. distachyon* plants (Table 4; Fig. 5; Supplementary Figs. S7, S8). Our selection analysis indicate that all *Bdhn* genes are apparently functional (Supplementary Table 6); however, the absence of transcripts from the other six genes indicate that they are not expressed in mature leaves of *B. distachyon* under the studied conditions. Abiotic stresses enhance the expression of particular DHN genes in plants (Graether & Boddington, 2014). *In-silico* and RT-PCR expression analysis showed that YSK2-type dehydrins were upregulated in drought-stressed shoots of wheat, whereas Kn-type dehydrins were preferentially expressed in cold-stressed shoots (Wang et al., 2014). Other drought-PEG treatments have shown considerable upregulation of eight DHN genes in rice shoots a few days after the stress (Verma et al., 2017). The expressions of the four *Brachypodium Bdhn* genes were upregulated by drought in all *B. distachyon* ecotypes (Fig. 5). The effect of drought significantly contributed to transcriptional upregulation of the four genes in dry vs watered plants, irrespective of the temperature treatment (Supplementary Figs. S7, S8). This analysis demonstrated a 2.5 to 21.5-fold increase in the expression (TPM) of *Bdhn*1a but a much higher and variable 2 to 490-fold increase in those of *Bdhn*2, *Bdhn*3 and *Bdhn*7 under dry than under water conditions (Table 4; Fig. 5; Supplementary Fig. S7). By contrast, the temperature effect did not affect the overall expression of the four genes, and only tests performed on watered plants showed significant DEs between cold vs hot conditions for *Bdhn*3 and *Bdhn*7 but not for *Bdhn*1a and *Bdhn*2 (Supplementary Fig. S7c). *Bdhn*1a and *Bdhn*2 are (FS)K_n_-type genes whereas *Bdhn*3 and *Bdhn*7 are Y_n_SK_n_-type genes (Table 1; Fig. 3); thus our results partially depart from those found in wheat by Wang et al., (2014) suggesting that both types of genes are preferentially upregulated by drought rather than by temperature in mature leaves and that cold and heat stress only affects Y_n_SK_n_-type genes in *B. distachyon*, even for the mild drought and temperature conditions of our experiments. By contrast, our comparative DE analysis between the upregulated *B. distachyon Bdhn* genes under drought conditions and those of wheat flag leaves under natural field drought stress (Galvez et al. 2019) or greenhouse-imposed drought stress (Reddy et al. 2014, Chu et al. 2021) inferred that 15 out of the 16 DE wheat dehydrins belong to the same *Brachypodium* DE ortholog groups (Supplementary Table 9; Supplementary Fig. S10). These shared functional responses to drought by orthologous dehydrin genes’ induction in both species reinforce the potential of *B. distachyon* as a model system for cereals, like bread wheat. Our results also highlight the importance of these four dehydrins in the protection of *B. distachyon* plants to water stress conditions in mature individuals, the developmental stage when they face the most severe drought conditions of their life cycle (Martínez et al., 2018)

The drought-induced upregulation of *Bdhn* genes was significantly different among the studied *B. distachyon* ecotypes for all four *Bdhn* genes (Table 4; Fig. 5; Supplementary Fig. S8). Recent analysis of dehydrin gene expression in *B. distachyon* and its close *B. stacei* and *B. hybridum* species have shown that *Bdhn*3 was strongly up-regulated by drought (>400-fold higher expression in dry-plants compared to control-plants), although the level of induction depended on genotype (Martínez et al., 2018). Our data support these results and additionally demonstrate that drought induction also upregulates the expressions of the *Bdhn*1a, *Bdhn*2 and *Bdhn*7 genes and that overexpression mostly affect the warm climate ecotypes (Koz3, Adi10) and less the cold climate ecotypes (ABR2, ABR4), whereas mesic climate ecotypes show intermediate and also significantly different expression levels (Table 4; Fig. 5; Supplementary Fig. S8). Fisher et al. (2016) classification of *B. distachyon* ecotypes into drought–tolerant, –intermediate, and –susceptible, based on phenotypic plant water content and wilting index values, roughly correspond to our warm, mesic and cold ecotype climate classes, thought their plants were subjected to uncontrolled severe drought treatments which may have confounded plant size with soil water content. The significant differences found between the drought-induced dehydrin expression levels in our climate class *B. distachyon* ecotypes (Table 4; Fig. 5; Supplementary Fig. S8) suggests that ecotypes adapted to warm climates may have developed higher *Bdhn*1a, *Bdhn*2, *Bdhn*3 and *Bdhn*7 expression inductions, a strategy to protect the mature plants against harsh water deprivation conditions and to ensure the survival and reproduction of the individuals in their habitats. By contrast, in mesic and, especially, in cold climate adapted ecotypes those inductions are much lower possibly due to the absence or mitigated presence of the natural stressor.

The constitutive and induced expression of the stress-responsive dehydrins is upregulated by the presence of specific cis-regulatory elements in the promoter region of their genes (Yamaguchi-Shinozaki & Shinozaki, 2005). Our *de novo* analysis of cis-regulatory elements consistently found three TF-binding sites, BES1/BZR1, MYB124 and ZAT, across the *Bdhn* promoters of the studied *Brachypodium* species, and more abundantly in the promoter regions of the *Bdhn*1a and *Bdhn*2 genes (Table 3; Fig. 2). BES1/BZR1 is a brassinosteroid signaling positive regulator (BZR1) family protein involved in the regulation pathway in response to drought (Cui et al., 2019). The MYB gene protein MYB124 is related to the abcisic acid (ABA) response, an hormone-regulated pathway implicated in multiple stress response such as drought or cold stress (Ambawat et al.,2013.; Khan et al., 2018). The ZAT C2H2 zinc finger is involved in response to salinity stress (Ciftci-Yilmaz et al., 2007). The presence of these cis-regulatory elements in the promoters of most *Bdhn* genes suggest that these dehydrins could be highly upregulated in *Brachypodium* plants under different water deficit stresses such as drought, cold and salinity. The scans of the three conserved motifs across the promoters of the ten *Bdhn* genes in the four *Brachypodium* species show that BES1 and ZAT were present in multiple *Bdhn* genes (BES1 in *Bdhn*2, *Bdhn*5, *Bdhn*7 and *Bdhn*8; ZAT in *Bdhn*4, *Bdhn*5 and *Bdhn*10), whereas MYB124 was constantly present in *Bdhn*2 within the *Brachypodium* genomes. The study of the four dehydrin genes upregulated by drought conditions in the *B. distachyon* ecotypes indicate that MYB124 and BES1 may play an important role in the induction of the *Bdhn*1a and *Bdhn*2 genes in the studied ecotypes, especially in those adapted to warm climates.

### Correlated dehydrin and phenotypic drought response and phylogenetic signal in Brachypodium distachyon

Water deficit stress affects the physiology, the phenotypes and the fitness of plants (Des Marais et al., 2017; Hossain et al., 2016). Des Marais et al. (2017) demonstrated that most of the 12 studied phenotypic traits across the *B. distachyon* ecotypes responded separately to soil drying, while several also displayed an interactive effect of temperature and soil water content. Only one trait – above ground biomass -- showed a significant main effect of temperature. Similarly, our dehydrin expression study has shown that dehydrins respond more strongly to dehydration than they do to elevated temperature, significantly affecting the upregulation of the four *Bdhn* genes, irrespective of the temperature condition (Supplementary Fig. S7). We did not directly test for temperature by water interaction in this analysis. As shown earlier by Des Marais et al (2017), drought effect also significantly influenced the changes of the phenotypic traits, reducing the water contents, total plant mass and leaf nitrogen content of dry plants but increasing their root biomass, leaf carbon content, proline and WUE (Supplementary Table 10; Supplementary Fig. S11). Our linear regression models indicated that the expressions of the four *Bdhn* genes were significantly correlated with the changes of most phenotypic traits in the drought treatment (Supplementary Table 11; Supplementary Fig. S12). The high correlations observed between the expression of *Bdhn*1a, *Bdhn*2, *Bdhn*3 and *Bdhn*7 genes and the decrease of leaf water and nitrogen contents and the increase of belowground biomass, root mass, and leaf carbon content and proline content (Supplementary Fig. S13) were strongly associated to drought stress. Our regression models also found significant correlation between *Bdhn*1a, *Bdhn*2, *Bdhn*3 and *Bdhn*7 dehydrin regulation and WUE increase (Supplementary Fig. S12). Martínez et al. (2018) reported a significant decrease in plant water content but no significant changes in proline content in *B. distachyon* ecotypes under drought stress. By contrast, Fisher et al. (2016) found significant differences in both traits across *B. distachyon* ecotypes under mild and severe drought treatments, with drought-tolerant ecotypes showing more prominent water and proline contents than the drought-intermediate and drought-susceptible ecotypes. Our data corroborate the last results and further illustrate that drought-induced proline overproduction is significantly higher in warm-to-mesic climate *B. distachyon* ecotypes (Adi10, Koz3) and lower in cold climate ecotypes (ABR5, ABR4, ABR3) (Supplementary Table 10) and that those differences overlap with the significant differences observed in their dehydrin overexpression profiles (Table 4; Fig. 5). Proline can serve as an osmoprotectant and a signaling molecule triggering adaptive responses to cell water stress (Verbruggen & Hermans, 2008). Our data indicate that drought-induced dehydrin expression is strongly correlated with proline synthesis (Supplementary Fig. S12). Our results also suggest that drought-induced dehydrin and proline upregulation are genotype-dependent, and mostly affect the *B. distachyon* warm climate ecotypes but not the cold climate ecotypes. In cool seasonal plants WUE is expected to increase with aridity (Cernusak et al., 2013). However, Manzaneda et al. (2015) found that drought-avoider *B. distachyton* ecotypes showed lower WUE than aridic drought-escape *B. hybridum* ecotypes, although the former had higher values of WUE plasticity related to climate than the second. Des Marais et al. (2017) also found an association of WUE with climate as *B. distachyon* ecotypes from cooler climates were more plastic in their WUE than those from warmer climates. Our data indicate that the overall dehydrin expression is significantly correlated with WUE (Supplementary Fig. S12). In addition, WUE shows a great plasticity across ecotypes of any climate class and under both drought and watered conditions (Supplemental Table 10).

The potential evolutionary signal of dehydrin expression values and phenotypic trait values gave different results when tested on the *B. distachyon* nuclear species tree or the *B. distachyon Bdhn* tree. The low-regulated expressions of *Bdhn*2 and *Bdhn*7 in watered plants and the upregulated expression of *Bdhn*3 in dry plants had moderate but significant or marginally significant phylogenetic signal on the *Bdhn* tree (Table 5a; Fig. 6a); however, none of the dehydrin expressions under W or D conditions showed significant phylogenetic signal on the nuclear species tree (Supplementary Table 12a; Supplementary Fig. S13a). In addition, seven out of the 12 drought-responsive phenotypic traits showed significant though low phylogenetic signal in the nuclear species tree (Supplementary Table 12b; Supplementary Fig. S13b), whereas five of those traits (root mass ratio, leaf nitrogen content and c:n ratio in watered conditions, and leaf relative water content and carbon content in dry conditions) showed high to moderate phylogenetic signal on the *Bdhn* tree (Table 5b; Fig. 6b). The phylogenetic signal of climate niche data (PCA1) was also significantly higher in the dehydrin *Bdhn* tree (Table 5c; Fig. 6c) than in the nuclear species tree (Supplementary Table 12c; Supplementary Fig. S13c). The absence of phylogenetic signal for the dehydrin expressions and the residual signal for some of their associated drought-response phenotypic traits in the nuclear species tree indicates that these drought-response mechanisms may have evolved independently and at different times along the life history of *B. distachyon*. However, several flowering time traits and their molecular regulators have shown a strongly correlated evolution with the nuclear species tree (Gordon et al., 2017), supporting the important role of flowering time in shaping the divergences of the *B. distachyon* lineages. Conversely, the significant phylogenetic signal of some dehydrin expressions, drought response phenotypic traits changes and climate niche data variation on the *B. distachyon Bdhn* tree suggests that the evolution of the *Bdhn* genes is determined by the adaptation of the *B. distachyon* ecotypes to more dry or more wet environmental conditions. It is surprising the high topological similarity found between the *Bdhn* tree and the plastome tree in contrast with its more dissimilar topology with respect to the nuclear species tree (Supplementary Fig. S5) for nuclear dehydrin genes that encode cytoplasmic and nuclear but not chloroplast proteins (Graether & Boddington, 2014). The relative congruence detected between the *Bdhn* and plastome trees could be a consequence of incomplete lineage sorting events of the recently evolved *B. distachyon* lineages (Sancho et al., 2018). However it could also imply a yet unknown organellar effect on the cellular response mechanism to adaptation to drought, like the role played by the chloroplast in inducing the expression of nuclear heat-response genes during heat stress in plants (Hu et al., 2020). Our data open new ways to investigate the potential implication of this organelle in the induction of drought-response nuclear genes, like those encoding for dehydrins, and in their evolutionary history.

## Conclusions

We have annotated and analyzed the ten *Brachypodium* dehydrin genes (*Bdhn*1-*Bdhn*10) present in the reference genomes of the three annual (*B. distachyon, B. stacei, B. hybridum*) and one perennial (*B. sylvaticum*) species of the genus. Most *Bdhn* genes have orthologs in other close grass species. Ancestral segmental and tandem duplications have been, respectively, detected in all species for the *Bdhn*1/*Bdhn*2 and *Bdhn*7/Bdh8 genes, and recent tandem duplications in *B. distachyon* for *Bdhn*4/*Bdhn*5 and in *B. sylvaticum* for *Bdhn*1a/*Bdhn*1b genes. Structural and biochemical properties of the *Brachypodium* dehydrins indicate that these disordered proteins may be present in the cytoplasmic and nuclear compartments of the cell. The three cis-regulatory elements identified in the promoter regions of the *Bdhn* genes suggests the predominant regulation of the *Bdhn* genes by ABA- and brassinosterioid-mediated response metabolic pathways. Only four dehydrin genes (*Bdhn*1a, *Bdhn*2, *Bdhn*3, *Bdhn*7) are expressed in mature laves of *B. distachyon*. Differential expression levels of these dehydrins are mainly induced by drought rather than temperature conditions and are genotype-dependent, being significantly higher in warm than in mesic or cold climate ecotypes. Drought-mediated dehydrin upregulation is significantly correlated with leaf water and nitrogen contents decreases and root biomass and leaf proline increases which are also genotype-dependent.

## Author contributions

PC, PH and SG designed the study. MD, SG, RS and DD generated the data. MD, SG, FA, RS, BC, DD and PC analyzed the data. MD, PH and PC wrote the draft manuscript. All authors contributed to the writing of the final version of the manuscript.

## Data availability Statement

The complete protocol, input and output data, and Supplementary information (Supplementary Data, Supplementary Tables, Supplementary Figures, Supplementary Materials) are available at Github (https://github.com/Bioflora/Brachypodium_dehydrins).

## Acknowledgements

This work was supported by the Spanish Ministries of Economy and Competitivity (Mineco) and of Science and Innovation CGL2016-79790-P and PID2019-108195GB-I00 grant projects. MD was funded by a Mineco FPI PhD fellowship. BCM was funded by Fundación ARAID. PC and MD were also funded by a European Social Fund/Aragón Government Bioflora research grant A01-17R. PH and SG were funded by project P18-RT-992 from Junta de Andalucía, Spain (co-funded by FEDER). The bioinformatic and evolutionary analyses were performed at the University of Malaga (Spain) and Escuela Politécnica Superior de Huesca (Universidad de Zaragoza, Spain) laboratories, respectively. Plant growth experiments and RNASequencing were supported USDA (NIFA-2011-67012-30663) to D.L.D.. The *B. sylvaticum* genome was used with permission under early release conditions of the DOE Joint Genome Institute.

## Supplementary Figures

**Figure S1.** Inferred tridimensional structure of some *Brachypodium* dehydrins: (A) *B. distachyon Bdhn*4; (B) *B. hybridum*D *Bdhn*3; (C) *B. stacei Bdhn*1; (D) *B.hybridum*S *Bdhn*10; (E): *B. sylvaticum Bdhn*1b. Complete sets of 3D structures for all *Brachypodium* species and proteins are available in the following links:

*B.distachyon* (https://galactus.uma.es/triticae/pipeline_web/production/compartida/raptorx/distachyon.php);

*B.hybridum*D (https://galactus.uma.es/triticae/pipeline_web/production/compartida/raptorx/hybridumD.php)

*B. stacei* (https://galactus.uma.es/triticae/pipeline_web/production/compartida/raptorx/stacei.php);

*B.hybridum*S (https://galactus.uma.es/triticae/pipeline_web/production/compartida/raptorx/hybridumS.php);

*B. sylvaticum* (https://galactus.uma.es/triticae/pipeline_web/production/compartida/raptorx/sylvaticum.php).

**Figure S2.** Analysis of *cis*-regulatory element discovery with Rsat::plants tools in the 5-upstream promoter region (−500-to-+200 bp) of 47 *Bdhn* genes from four *Brachypodium* species. **(a)** Maximum k-mer significance values from each analysis are shown for *Bdhn* sequences from each species; **(b)** Maximum number of sites from each background analysis are shown for *Bdhn* sequences from each species. Species: *B. distachyon* (blue); *B. stacei* (red); *B. hybridum* (purple); *B. sylvaticum* (green). In each case, another 47 random gene sequences from the respective reference genome were analyzed ten times as controls (grey bars; C1-C10).

**Figure S3.** Chromosomal location of the 10 *Bdhn* genes in 54 *B. distachyon* ecotypes. *Bdhn* genes (clusters) detected in each ecotype were compared against the dehydrins of the reference genome (Bd21 v3) and assigned to the cluster with highest score using global pairwise alignment (Needleman-Wunsch with BLOSUM62). *Bdhn* color codes and the accuracy of the annotations are indicated in the charts.

**Figure S4.** Maximum likelihood *B. distachyon* dehydrin trees obtained from the aligned exon and intron sequences of each independent *Bdhn* gene (*Bdhn*1 to *Bdhn*10). Trees were constructed with IQTREE using the *B. stacei* outgroup sequence to root the tree. Ultrafast bootstrap support is indicated on branches. Accession codes correspond to those indicated in Supplementary Table S1.

**Figure S5. (a)** Maximum likelihood *B. distachyon Bdhn* tree. Consensus tree constructed for 54 *B. distachyon* ecotypes from concatenated exon and intron aligned sequences of six dehydrin genes (*Bdhn*1, *Bdhn*2, *Bdhn*3, *Bdhn*6, *Bdhn*7, *Bdhn*8) that showed congruent topologies with IQTREE. The EDF+ (extremely delayed flowering time, blue) clade and T+ (Turkish and East Mediterranean, orange) and S+ (Spain and West Mediterranean, green) lineages correspond to those indicated in Gordon et al. (2017) and Sancho et al (2018). Ultrafast bootstrap support is indicated on branches. Accession codes correspond to those indicated in Supplementary Table S1. **(b)** *B. distachyon* nuclear species tree of Gordon et al. (2017) based on nuclear genome-wide 3.9 million SNPs. **(c)** *B. distachyon* plastome tree of Sancho et al. (2018) based on full plastome sequences.

**Figure S6.** Bidimensional PCA plot of 54 *B. distachyon* ecotypes obtained from 19 climate variables (see Supplementary Table S3). PC1 and PC2 axes accumulate 48.5% and 22.4% of the variance, respectively. Ecotype codes correspond to those indicated in Supplementary Table S1. Ellipses include ecotypes classified within cold (aquamarine), mesic (green) and warm (red) climate classes according to their PC1 values.

**Figure S7.** Boxplots and Wilcoxon pairwise significance tests of differential gene expression values (normalized transcript per million, TPM) of the four expressed dehydrin genes (*Bdhn*1a, *Bdhn*2, *Bdhn*3, *Bdhn*7) under joint and separately analyzed drought and temperature stress conditions in 32 *B. distachyon* ecotypes. Averaged expression values for C (Cold), H (Hot), W (Watered), and D (Drought) treatments and their combinations (see text). **(a)** Pairwise comparative tests of combined CD-CW-HD-HW treatments; all dehydrins were significantly differentially expressed in all CD vs CW and HD vs HW tests, by contrast they were not significantly different in all CD vs HD tests and in two CW vs HW tests (*Bdhn*1a, *Bdhn*2); **(b)** Pairwise comparative tests of D vs W treatments; all dehydrins were significantly differentially expressed; **(c)** Pairwise comparative tests of C vs H treatments; none of the dehydrins were significantly differentially expressed.

**Figure S8.** Differentially expressed *Bdhn*1a, *Bdhn*2, *Bdhn*3 and *Bdhn*7 dehydrin genes (normalized transcript per million, TPM) across 32 ecotypes of *B. distachyon* under drought (D, red) vs watered (W, blue) conditions. Different letters in the boxplots indicate significant group differences (Tukey tests).

**Figure S9.** Linear model regression plots of pairwise dehydrine *Bdhn* expression values (normalized transcript per million, TPM) under joint drought and watered conditions.

**Figure S10.** Physical locations of orthologous *Brachypodium distachyon* and *Triticum aestivum* water stress responsive dehydrin genes. Drough-induced wheat dehydrin genes are highlighted in colors. *Brachypodium* chomosomes are drawn at 10x scale with respect to wheat chromosomes.

**Figure S11.** Drought-response phenotypic changes as a function of drought treatment averaged across 32 *B. distachyon* ecotypes [ leaf_rwc (relative water content in leaf); leaf_wc (water content in leaf); lma (leaf mass per area); pro (leaf proline content); abvrgd (above ground biomass); blwgrd (below ground biomass); ttlmass (total mass); rmr (root mass ratio); delta13c (carbon isotope, a proxy for lifetime integrated WUE); leafc (leaf carbon content); leafn (leaf nitrogen content); cn (leaf carbon/nitrogen ratio)]. Asterisks above boxes indicate Wilcoxon pairwise significant difference among drought (D, red) and watered (W, blue) conditions (p-value<0.001, ***).

**Figure S12.** Linear model regression plots of dehydrine *Bdhn*1a, *Bdhn*2, *Bdhn*3 and *Bdhn*7 expression values (normalized transcript per million, TPM) and drought-response phenotypic traits changes under total dry (D) and watered (W) conditions.

**Figure S13.** Maximum Likelihood *B. distachyon* nuclear species tree cladogram showing the relationships of 30 ecotypes. Phyloheatmaps of normalized values for different sets of variables: (a) dehydrin (*Bdhn*1, *Bdhn*2, *Bdhn*3, *Bdhn*7) gene expression values under watered (W) and drought (D) conditions; (b) drought-response phenotypic traits (leaf_rwc; leaf_wc; lma; pro; abvrgd; blwgrd; ttlmass; rmr; delta13c; leafc; leafn; cn) values under watered (W) and drought (D) conditions; (c) climate niche PCA1 values. Traits showing significant phylogenetic signal are highlighted with dotted lines (see Supplementary Table 12).

