## Supplementary Tables for "Evolution and functional dynamics of dehydrins in model *Brachypodium* grasses"

**Supplementary Table 1**. Sampling origins of the four studied Brachypodium species and 54 ecotypes of B. distachyon. All samples were used in the genomic analysis of the dehydrin genes. Asterisks indicate B. distachyon accessions additionally used in the dehydrin expression and drought-response phenotypic traits changes analyses (32 ecotypes); diamonds indicate accessions additionally used in the phylogenetic signal analysis (30 ecotypes).

| **Species** | **accession** | **longitude** | **latitude** | **Locality** |
| --- | --- | --- | --- | --- |
| *B.stacei* | ABR114 | 38.682846 | 1.398957 | Spain: Balearic isles, Formentera, Torrent |
| *B.hybridum* | ABR113 | 38.782993 | -9.250488 | Portugal: Lisboa, Belas |
| *B.sylvaticum* | Ain1 | 36.768235 | 8.707878 | Tunisia: Ain-Draham |
| *B.distachyon* | ABR2*♦ | 3,3000 | 43,6500 | France: Herault, Octon |
| *B.distachyon* | ABR3*♦ | 0,0731 | 42,1805 | Spain: Huesca, Aisa |
| *B.distachyon* | ABR4*♦ | 0,7168 | 42,2627 | Spain: Huesca, Aren |
| *B.distachyon* | ABR5*♦ | -0,5800 | 42,5810 | Spain: Huesca, Jaca, Banaguas |
| *B.distachyon* | ABR6*♦ | -2,2030 | 42,5810 | Spain: Navarra, Los Arcos |
| *B.distachyon* | ABR8*♦ | 11,3197 | 43,3146 | Italy: Siena |
| *B.distachyon* | ABR9 | 14,4895 | 46,0609 | Croatia: Ljubjana |
| *B.distachyon* | Adi10*♦ | 38,3523 | 38,7707 | Turkey: Adiyaman |
| *B.distachyon* | Adi12*♦ | 38,3523 | 38,7707 | Turkey: Adiyaman |
| *B.distachyon* | Adi2*♦ | 38,3523 | 38,7707 | Turkey: Adiyaman |
| *B.distachyon* | Arn1 | 0,7299 | 42,2565 | Spain: Huesca, Arén |
| *B.distachyon* | Bd1_1*♦ | 28,2510 | 38,4170 | Turkey: Manisa |
| *B.distachyon* | Bd18_1*♦ | 33,7300 | 39,3678 | Turkey: Kaman |
| *B.distachyon* | Bd2_3*♦ | 44,4031 | 33,7609 | Irak: Al Mansuriya |
| *B.distachyon* | Bd21_3*♦ | 44,5350 | 36,7660 | Irak: near Salakudin |
| *B.distachyon* | Bd29_1 | 33,5639 | 44,5153 | Ukraine: Krimea |
| *B.distachyon* | Bd3_1*♦ | 44,5350 | 36,7660 | Irak: Al Mansuriya |
| *B.distachyon* | BdTR10C*♦ | 31,8849 | 37,7782 | Turkey: Konya Province |
| *B.distachyon* | BdTR11A | 31,8849 | 37,7782 | Turkey: Konya Province |
| *B.distachyon* | BdTR11G*♦ | 27,4770 | 41,4220 | Turkey: Kirklareli |
| *B.distachyon* | BdTR11I*♦ | 28,0402 | 39,7382 | Turkey: Balikesir, Karakaya |
| *B.distachyon* | BdTR12C* | 34,6503 | 39,7482 | Turkey: Saray, Yozgat province |
| *B.distachyon* | BdTR13a*♦ | 32,4324 | 39,7565 | Turkey: Ankara |
| *B.distachyon* | BdTR13C* | 32,9881 | 39,4129 | Turkey: Ankara |
| *B.distachyon* | BdTR1i*♦ | 28,5830 | 38,0930 | Turkey: Aydin |
| *B.distachyon* | BdTR2B*♦ | 31,3311 | 40,0821 | Turkey: Karahisarkozlu |
| *B.distachyon* | BdTR2G*♦ | 32,9850 | 40,3940 | Turkey: Ankara |
| *B.distachyon* | BdTR3C*♦ | 32,9630 | 36,7830 | Turkey: Balkusan |
| *B.distachyon* | BdTR5i*♦ | 32,9854 | 40,3936 | Turkey: Cubuk |
| *B.distachyon* | BdTR7A | 34,6500 | 39,7480 | Turkey: Yozgat |
| *B.distachyon* | BdTR8i | 34,0714 | 37,1885 | Turkey: Berendi |
| *B.distachyon* | BdTR9K*♦ | 30,7886 | 39,7530 | Turkey: Eskisehir |
| *B.distachyon* | Bis1*♦ | 41,0151 | 37,8735 | Turkey: Bismil |
| *B.distachyon* | Foz1 | -1,3050 | 42,6370 | Spain: Navarra, Foz de Lumbier |
| *B.distachyon* | Gaz8 | 37,3910 | 37,1280 | Turkey: Gaziantep |
| *B.distachyon* | Jer1 | 0,0120 | 42,0550 | Spain: Huesca, Adahuesca |
| *B.distachyon* | Kah1*♦ | 38,5330 | 37,7340 | Turkey: Kahta |
| *B.distachyon* | Kah5*♦ | 38,5330 | 37,7340 | Turkey: Kahta |
| *B.distachyon* | Koz1*♦ | 41,6100 | 38,1520 | Turkey: Kozluk |
| *B.distachyon* | Koz3*♦ | 41,6100 | 38,1520 | Turkey: Kozluk |
| *B.distachyon* | Luc1 | -0,8930 | 42,6100 | Spain: Huesca, Berdun |
| *B.distachyon* | Mig3 | -0,2050 | 42,1470 | Spain: Huesca, Ibieca, San Miguel de Foces |
| *B.distachyon* | Mon3 | -0,2090 | 41,6520 | Spain: Zaragoza, Castejón de Monegros |
| *B.distachyon* | Mur1 | 0,8770 | 42,0980 | Spain: Lleida, Castillo de Mur |
| *B.distachyon* | Per1 | -1,7500 | 42,7370 | Spain: Navarra, Puerto del Perdon |
| *B.distachyon* | S8iiC | 0,1440 | 41,6054 | Spain: Huesca, Zaidín |
| *B.distachyon* | Sig2 | -1,0150 | 42,6130 | Spain: Zaragoza, Sigüés |
| *B.distachyon* | Tek2 | 26,9310 | 41,0850 | Turkey: Tekirdag |
| *B.distachyon* | Tek4 | 27,5191 | 41,0112 | Turkey: Tekirdag |

**Supplementary Table 2**. Brachypodium distachyon dehydrin expression data. Filtered and normalized transcripts per million (TPM) values of annotated dehydrins. Only four dehydrin genes (Bdhn1a, Bdhn2, Bdhn3, Bdhn7) were expressed in leaves of 3-to-4-days grown plants. Plants were subjected to drought (W: watered, D: Drought) and temperature (C: Cold, H: Hot) stress conditions (see text). Code indicates the sampling code used in the RNAseq analysis.

| **Drougth (D)** | | | | | |  | **Watered (W)** | | | | | |
| --- | --- | --- | --- | --- | --- | --- | --- | --- | --- | --- | --- | --- |
| **Code** | **accession** | ***Bdhn*1a** | ***Bdhn*2** | ***Bdhn*3** | ***Bdhn*7** |  | **Code** | **accession** | ***Bdhn*1a** | ***Bdhn*2** | ***Bdhn*3** | ***Bdhn*7** |
| BA030_HD_ABR2 | ABR2 | 309,6 | 207,4 | 131,8 | 21 |  | BA053_HW_ABR2 | ABR2 | 199,6 | 27,2 | 5,3 | 1 |
| BA101_HD_ABR2 | ABR2 | 150,4 | 142,4 | 278,7 | 19 |  | BA085_HW_ABR2 | ABR2 | 143,6 | 67,5 | 39,1 | 3,6 |
| BA366_CD_ABR2 | ABR2 | 181,8 | 183,1 | 227,7 | 26,6 |  | BA145_HW_ABR2 | ABR2 | 132,7 | 36,5 | 0 | 1,7 |
| BA439_CD_ABR2 | ABR2 | 214,9 | 135,5 | 51,5 | 11 |  | BA403_CW_ABR2 | ABR2 | 163,6 | 53 | 30,2 | 2,9 |
| BA006_HD_ABR3 | ABR3 | 304,9 | 182,2 | 32,1 | 10,7 |  | BA447_CW_ABR2 | ABR2 | 120,5 | 52 | 26 | 4,7 |
| BA103_HD_ABR3 | ABR3 | 197,7 | 326 | 738,2 | 75,7 |  | BA096_HW_ABR3 | ABR3 | 248,9 | 29,4 | 8,5 | 3,8 |
| BA418_CD_ABR3 | ABR3 | 328,3 | 116,6 | 43,9 | 9,4 |  | BA146_HW_ABR3 | ABR3 | 132,4 | 13,2 | 3,3 | 0 |
| BA465_CD_ABR3 | ABR3 | 224,2 | 51,4 | 65,3 | 22,3 |  | BA413_CW_ABR3 | ABR3 | 193,5 | 46,6 | 26,9 | 1,8 |
| BA038_HD_ABR4 | ABR4 | 299,5 | 76,8 | 7,3 | 2,4 |  | BA419_CW_ABR3 | ABR3 | 205,1 | 69,4 | 10,1 | 1,3 |
| BA170_HD_ABR4 | ABR4 | 226,2 | 194,7 | 164,3 | 37,6 |  | BA040_HW_ABR4 | ABR4 | 109,5 | 27,1 | 1,2 | 0 |
| BA368_CD_ABR4 | ABR4 | 270,3 | 65,9 | 15,7 | 15,7 |  | BA043_HW_ABR4 | ABR4 | 230,1 | 35,3 | 7,3 | 3,7 |
| BA521_CD_ABR4 | ABR4 | 192,6 | 96,3 | 61,4 | 4,7 |  | BA477_CW_ABR4 | ABR4 | 594,4 | 80,7 | 5,8 | 2,7 |
| BA024_HD_ABR5 | ABR5 | 335,1 | 298,2 | 62,4 | 17 |  | BA508_CW_ABR4 | ABR4 | 143,5 | 50,5 | 5,8 | 3,2 |
| BA104_HD_ABR5 | ABR5 | 234,1 | 580,1 | 681,7 | 92,8 |  | BA161_HW_ABR5 | ABR5 | 143,9 | 33,7 | 7,6 | 2,4 |
| BA454_CD_ABR5 | ABR5 | 188,7 | 55,4 | 140,2 | 45 |  | BA479_CW_ABR5 | ABR5 | 418 | 55,8 | 3,9 | 0,4 |
| BA522_CD_ABR5 | ABR5 | 217,7 | 279,5 | 46,2 | 21,9 |  | BA502_CW_ABR5 | ABR5 | 254,8 | 23,3 | 9,4 | 6,1 |
| BA037_HD_ABR6 | ABR6 | 333,5 | 130,4 | 147,8 | 13,5 |  | BA123_HW_ABR6 | ABR6 | 357,3 | 39,7 | 3,6 | 10,8 |
| BA099_HD_ABR6 | ABR6 | 281,5 | 397,3 | 2175,5 | 102,5 |  | BA153_HW_ABR6 | ABR6 | 340,4 | 26,1 | 3,9 | 3,9 |
| BA416_CD_ABR6 | ABR6 | 289 | 199,9 | 778,6 | 33,6 |  | BA437_CW_ABR6 | ABR6 | 258,1 | 26,1 | 4,4 | 2,5 |
| BA523_CD_ABR6 | ABR6 | 256,7 | 90,5 | 300,3 | 16,6 |  | BA452_CW_ABR6 | ABR6 | 308,2 | 18,1 | 3,1 | 2,6 |
| BA100_HD_ABR8 | ABR8 | 298,2 | 144,7 | 199,2 | 63,3 |  | BA008_HW_ABR8 | ABR8 | 224,6 | 36,4 | 4,2 | 0 |
| BA143_HD_ABR8 | ABR8 | 379,6 | 336,2 | 262,1 | 112 |  | BA138_HW_ABR8 | ABR8 | 128,8 | 21,1 | 3,5 | 0 |
| BA415_CD_ABR8 | ABR8 | 502,2 | 312,7 | 177,6 | 84,1 |  | BA360_CW_ABR8 | ABR8 | 249 | 35,8 | 5,6 | 1,7 |
| BA506_CD_ABR8 | ABR8 | 940,2 | 644,1 | 285,3 | 251,7 |  | BA517_CW_ABR8 | ABR8 | 292,4 | 23,1 | 3,3 | 2,3 |
| BA049_HD_Adi10 | Adi10 | 440 | 197,7 | 215,9 | 112,4 |  | BA067_HW_Adi10 | Adi10 | 632,4 | 37,3 | 1,7 | 0,4 |
| BA052_HD_Adi10 | Adi10 | 357,1 | 249,5 | 258,2 | 64,7 |  | BA105_HW_Adi10 | Adi10 | 415,9 | 49,3 | 8,4 | 3,1 |
| BA478_CD_Adi10 | Adi10 | 820,4 | 1251,9 | 1961,7 | 462 |  | BA428_CW_Adi10 | Adi10 | 841,2 | 51 | 8,7 | 6,5 |
| BA513_CD_Adi10 | Adi10 | 1020,1 | 2043,2 | 2431,9 | 692,5 |  | BA520_CW_Adi10 | Adi10 | 341,2 | 20,1 | 8,1 | 2,9 |
| BA036_HD_Adi12 | Adi12 | 558 | 751,2 | 290,8 | 59,5 |  | BA041_HW_Adi12 | Adi12 | 339 | 81,9 | 4,9 | 2,4 |
| BA140_HD_Adi12 | Adi12 | 377,1 | 260,2 | 133 | 56,5 |  | BA044_HW_Adi12 | Adi12 | 396,1 | 43,2 | 0 | 0 |
| BA455_CD_Adi12 | Adi12 | 778,7 | 840 | 1104,1 | 330 |  | BA407_CW_Adi12 | Adi12 | 193,2 | 96 | 30,9 | 13,7 |
| BA525_CD_Adi12 | Adi12 | 240,8 | 515,5 | 466,2 | 113,3 |  | BA423_CW_Adi12 | Adi12 | 746,4 | 53,9 | 11,2 | 1,7 |
| BA050_HD_Adi2 | Adi2 | 458,9 | 161,7 | 37,1 | 11 |  | BA176_HW_Adi2 | Adi2 | 402,2 | 68 | 5 | 3,8 |
| BA094_HD_Adi2 | Adi2 | 396,1 | 457,9 | 318,7 | 55,9 |  | BA357_CW_Adi2 | Adi2 | 489,2 | 36,5 | 1,1 | 3,2 |
| BA500_CD_Adi2 | Adi2 | 528,3 | 913,4 | 957 | 123,8 |  | BA468_CW_Adi2 | Adi2 | 398,6 | 65,7 | 4 | 2 |
| BA509_CD_Adi2 | Adi2 | 456,6 | 405,1 | 463,1 | 64,5 |  | BA354_CW_Bd1-1 | Bd1-1 | 390,9 | 46,9 | 8,1 | 1,5 |
| BA025_HD_Bd1-1 | Bd1-1 | 475,8 | 282,7 | 88,6 | 27,7 |  | BA496_CW_Bd1-1 | Bd1-1 | 320 | 17,6 | 14,4 | 4 |
| BA051_HD_Bd1-1 | Bd1-1 | 352,8 | 22 | 3,3 | 2,2 |  | BA060_HW_Bd18-1 | Bd18-1 | 239,9 | 21,8 | 5 | 0 |
| BA122_HD_Bd1-1 | Bd1-1 | 235,1 | 42,5 | 53,7 | 4,5 |  | BA063_HW_Bd18-1 | Bd18-1 | 278,7 | 16,2 | 2,8 | 0,9 |
| BA442_CD_Bd1-1 | Bd1-1 | 369,3 | 70,6 | 42 | 14,3 |  | BA375_CW_Bd18-1 | Bd18-1 | 160,7 | 14,5 | 6,9 | 0,8 |
| BA475_CD_Bd1-1 | Bd1-1 | 582,4 | 352,4 | 578,5 | 79,5 |  | BA446_CW_Bd18-1 | Bd18-1 | 266,7 | 14,6 | 6,3 | 0 |
| BA093_HD_Bd18-1 | Bd18-1 | 464,3 | 392 | 257,2 | 138,3 |  | BA069_HW_Bd21 | Bd21 | 324,2 | 24,3 | 1,3 | 0,7 |
| BA453_CD_Bd18-1 | Bd18-1 | 1009,6 | 704,2 | 564 | 207,5 |  | BA112_HW_Bd21 | Bd21 | 363,1 | 70 | 24 | 5,7 |
| BA056_HD_Bd21 | Bd21 | 292,8 | 125 | 23,5 | 12,3 |  | BA386_CW_Bd21 | Bd21 | 183,1 | 24,9 | 2,4 | 1 |
| BA163_HD_Bd21 | Bd21 | 249,1 | 129,2 | 28,3 | 45,6 |  | BA456_CW_Bd21 | Bd21 | 330 | 59,6 | 37,8 | 6 |
| BA459_CD_Bd21 | Bd21 | 440,2 | 148,4 | 65,8 | 36,5 |  | BA054_HW_Bd21-3 | Bd21-3 | 307,6 | 25,8 | 9,4 | 1,8 |
| BA499_CD_Bd21 | Bd21 | 325,3 | 52,3 | 4,7 | 2,9 |  | BA171_HW_Bd21-3 | Bd21-3 | 276,1 | 26,2 | 18,8 | 0,7 |
| BA097_HD_Bd21-3 | Bd21-3 | 478,7 | 621,3 | 567 | 243,4 |  | BA458_CW_Bd21-3 | Bd21-3 | 288,6 | 25,4 | 14,7 | 2 |
| BA111_HD_Bd21-3 | Bd21-3 | 452,8 | 647 | 487,3 | 145,7 |  | BA086_HW_Bd2-3 | Bd2-3 | 181,2 | 37,4 | 5 | 2,1 |
| BA430_CD_Bd21-3 | Bd21-3 | 966,8 | 1402 | 1464,4 | 692,9 |  | BA114_HW_Bd2-3 | Bd2-3 | 422,7 | 54,3 | 0 | 0 |
| BA512_CD_Bd21-3 | Bd21-3 | 503,2 | 539,2 | 266,5 | 103,9 |  | BA487_CW_Bd2-3 | Bd2-3 | 327 | 82,5 | 6,9 | 3 |
| BA023_HD_Bd2-3 | Bd2-3 | 531,4 | 253 | 121,1 | 16,7 |  | BA492_CW_Bd2-3 | Bd2-3 | 542,5 | 48,5 | 3,8 | 4,9 |
| BA088_HD_Bd2-3 | Bd2-3 | 635,6 | 173,8 | 50,8 | 38,3 |  | BA005_HW_Bd30-1 | Bd30-1 | 266,2 | 37,1 | 5 | 3 |
| BA353_CD_Bd2-3 | Bd2-3 | 291,6 | 40,1 | 6,9 | 1,9 |  | BA018_HW_Bd30-1 | Bd30-1 | 244,7 | 20,3 | 2,5 | 0,4 |
| BA494_CD_Bd2-3 | Bd2-3 | 550,1 | 341,3 | 200,5 | 60,2 |  | BA417_CW_Bd30-1 | Bd30-1 | 504,7 | 32,9 | 8,1 | 2,9 |
| BA079_HD_Bd30-1 | Bd30-1 | 353,5 | 492,1 | 287 | 76,5 |  | BA474_CW_Bd30-1 | Bd30-1 | 444,6 | 44 | 7,3 | 2,9 |
| BA162_HD_Bd30-1 | Bd30-1 | 198,6 | 406,5 | 439,7 | 73,3 |  | BA007_HW_Bd3-1 | Bd3-1 | 321,4 | 34,7 | 4,6 | 1,5 |
| BA425_CD_Bd30-1 | Bd30-1 | 1005,9 | 1632,6 | 2616,2 | 685,5 |  | BA012_HW_Bd3-1 | Bd3-1 | 287,1 | 46,4 | 2,6 | 2,2 |
| BA481_CD_Bd30-1 | Bd30-1 | 396,7 | 515,6 | 283,3 | 55,5 |  | BA429_CW_Bd3-1 | Bd3-1 | 477,4 | 43 | 20,9 | 0 |
| BA166_HD_Bd3-1 | Bd3-1 | 316,4 | 405,3 | 1017,8 | 55,9 |  | BA434_CW_Bd3-1 | Bd3-1 | 256,1 | 46,6 | 10 | 7,2 |
| BA398_CD_Bd3-1 | Bd3-1 | 441,6 | 446,3 | 730,1 | 78 |  | BA119_HW_BdTR10c | BdTR10c | 196,7 | 36,9 | 0 | 0 |
| BA422_CD_Bd3-1 | Bd3-1 | 427,5 | 544,5 | 3386,7 | 854,4 |  | BA131_HW_BdTR10c | BdTR10c | 232 | 65 | 25,5 | 0 |
| BA121_HD_BdTR10c | BdTR10c | 242,1 | 214,3 | 471,6 | 66,8 |  | BA384_CW_BdTR10c | BdTR10c | 223 | 29,9 | 1,3 | 0 |
| BA173_HD_BdTR10c | BdTR10c | 380,8 | 122,7 | 284,2 | 82 |  | BA436_CW_BdTR10c | BdTR10c | 177,4 | 56,1 | 7,1 | 3,5 |
| BA421_CD_BdTR10c | BdTR10c | 603,2 | 483 | 1912,7 | 439,9 |  | BA021_HW_BdTR11g | BdTR11g | 320,5 | 63 | 6,2 | 1,4 |
| BA440_CD_BdTR10c | BdTR10c | 415,4 | 490,6 | 1028,6 | 187,7 |  | BA032_HW_BdTR11g | BdTR11g | 547 | 46,8 | 9,1 | 2,9 |
| BA059_HD_BdTR11g | BdTR11g | 452,4 | 370,4 | 615,1 | 63,5 |  | BA406_CW_BdTR11g | BdTR11g | 373,6 | 78,3 | 51,5 | 22,4 |
| BA090_HD_BdTR11g | BdTR11g | 590,1 | 645,2 | 1214,5 | 215,9 |  | BA493_CW_BdTR11g | BdTR11g | 314,6 | 49,3 | 3,6 | 2,1 |
| BA397_CD_BdTR11g | BdTR11g | 479,7 | 522,4 | 607 | 65,4 |  | BA042_HW_BdTR11i | BdTR11i | 274,4 | 30,1 | 2,8 | 2,1 |
| BA510_CD_BdTR11g | BdTR11g | 466,4 | 280,8 | 135,7 | 37,8 |  | BA148_HW_BdTR11i | BdTR11i | 230,3 | 39,3 | 4,2 | 1,4 |
| BA102_HD_BdTR11i | BdTR11i | 483,9 | 567,1 | 981,6 | 139,8 |  | BA377_CW_BdTR11i | BdTR11i | 258,1 | 27,6 | 4 | 2,2 |
| BA128_HD_BdTR11i | BdTR11i | 505,6 | 651 | 1554,8 | 175,1 |  | BA526_CW_BdTR11i | BdTR11i | 443,8 | 72,9 | 8,3 | 5,2 |
| BA408_CD_BdTR11i | BdTR11i | 708,1 | 680,3 | 833,4 | 135,7 |  | BA022_HW_BdTR13a | BdTR13a | 345,6 | 79,5 | 5,7 | 0 |
| BA450_CD_BdTR11i | BdTR11i | 429,9 | 314,4 | 305,8 | 46,7 |  | BA174_HW_BdTR13a | BdTR13a | 324 | 45,8 | 2,6 | 2,6 |
| BA361_CD_BdtR12c | BdtR12c | 297,8 | 37 | 9,7 | 5,8 |  | BA371_CW_BdTR13a | BdTR13a | 213,6 | 23,8 | 6,7 | 0 |
| BA061_HD_BdTR13a | BdTR13a | 382 | 213,3 | 110,5 | 41,8 |  | BA405_CW_BdTR13a | BdTR13a | 310,6 | 30,7 | 14,6 | 3,8 |
| BA144_HD_BdTR13a | BdTR13a | 369,6 | 249,2 | 130,1 | 69,2 |  | BA070_HW_BdTR1i | BdTR1i | 274,8 | 23,6 | 2,5 | 0 |
| BA394_CD_BdTR13a | BdTR13a | 363,5 | 160 | 95,3 | 31,8 |  | BA155_HW_BdTR1i | BdTR1i | 375,1 | 48,1 | 5,9 | 3,8 |
| BA460_CD_BdTR13a | BdTR13a | 1347,2 | 930,2 | 735,4 | 270,3 |  | BA469_CW_BdTR1i | BdTR1i | 409,4 | 34,5 | 9 | 3,3 |
| BA108_HD_BdTR1i | BdTR1i | 639,7 | 788 | 1133,3 | 488,1 |  | BA486_CW_BdTR1i | BdTR1i | 348,4 | 78,2 | 5,4 | 2,6 |
| BA134_HD_BdTR1i | BdTR1i | 315,7 | 191,5 | 199,9 | 73,1 |  | BA115_HW_BdTR2b | BdTR2b | 348,7 | 11,1 | 0 | 0 |
| BA457_CD_BdTR1i | BdTR1i | 579,7 | 591,8 | 246,9 | 79 |  | BA133_HW_BdTR2b | BdTR2b | 270,5 | 58,2 | 6,4 | 3,9 |
| BA480_CD_BdTR1i | BdTR1i | 796,6 | 989,7 | 1329,2 | 364,4 |  | BA378_CW_BdTR2b | BdTR2b | 259,5 | 14,5 | 3,5 | 1,2 |
| BA003_HD_BdTR2b | BdTR2b | 874,2 | 662,1 | 896,5 | 314,5 |  | BA503_CW_BdTR2b | BdTR2b | 377,6 | 73,2 | 10,9 | 2,1 |
| BA091_HD_BdTR2b | BdTR2b | 658,5 | 524,5 | 502,4 | 140 |  | BA107_HW_BdTR2g | BdTR2g | 475,5 | 44,3 | 3,2 | 4,4 |
| BA389_CD_BdTR2b | BdTR2b | 299,5 | 213,3 | 177,2 | 58,6 |  | BA167_HW_BdTR2g | BdTR2g | 401,5 | 36,3 | 7,7 | 3,3 |
| BA472_CD_BdTR2b | BdTR2b | 320,5 | 222,3 | 168,1 | 32,5 |  | BA445_CW_BdTR2g | BdTR2g | 334,5 | 41,6 | 5,2 | 4,2 |
| BA124_HD_BdTR2g | BdTR2g | 784,1 | 1119 | 2009,5 | 616,6 |  | BA527_CW_BdTR2g | BdTR2g | 594,9 | 48,9 | 7,4 | 2,5 |
| BA165_HD_BdTR2g | BdTR2g | 402,6 | 473,2 | 1420,3 | 511,6 |  | BA071_HW_BdTR3c | BdTR3c | 349,4 | 22 | 2,1 | 1,4 |
| BA364_CD_BdTR2g | BdTR2g | 1009,2 | 1135,3 | 2063,8 | 712,3 |  | BA129_HW_BdTR3c | BdTR3c | 227,6 | 38,8 | 2,6 | 2,6 |
| BA369_CD_BdTR2g | BdTR2g | 806,3 | 537,8 | 578,2 | 230,2 |  | BA362_CW_BdTR3c | BdTR3c | 368,9 | 105,8 | 162,7 | 53,3 |
| BA073_HD_BdTR3c | BdTR3c | 432,8 | 335,1 | 402,5 | 61,8 |  | BA427_CW_BdTR3c | BdTR3c | 234,2 | 75,7 | 8,1 | 3,9 |
| BA172_HD_BdTR3c | BdTR3c | 676,9 | 752,3 | 1712,3 | 351,6 |  | BA160_HW_BdTR5i | BdTR5i | 258,8 | 48,1 | 3,3 | 0,4 |
| BA370_CD_BdTR3c | BdTR3c | 542,5 | 585,9 | 354,6 | 53,6 |  | BA372_CW_BdTR5i | BdTR5i | 119,9 | 23,4 | 9,3 | 1,5 |
| BA424_CD_BdTR3c | BdTR3c | 985,9 | 1131,8 | 1662 | 310,6 |  | BA464_CW_BdTR5i | BdTR5i | 455,1 | 38,7 | 6,5 | 5,2 |
| BA065_HD_BdTR5i | BdTR5i | 358,7 | 296 | 254,4 | 57,5 |  | BA113_HW_BdTR9k | BdTR9k | 270,8 | 55,8 | 0 | 0 |
| BA082_HD_BdTR5i | BdTR5i | 366,7 | 390,6 | 459,3 | 80,5 |  | BA125_HW_BdTR9k | BdTR9k | 282,5 | 32,3 | 5,4 | 2,7 |
| BA470_CD_BdTR5i | BdTR5i | 367,3 | 369,8 | 607 | 94 |  | BA383_CW_BdTR9k | BdTR9k | 309,8 | 41,3 | 2,6 | 0,4 |
| BA473_CD_BdTR5i | BdTR5i | 345,6 | 298,7 | 254,6 | 50,9 |  | BA515_CW_BdTR9k | BdTR9k | 0 | 0 | 0 | 0 |
| BA002_HD_BdTR9k | BdTR9k | 377,8 | 193,6 | 54,1 | 21,5 |  | BA118_HW_Bis1 | Bis1 | 252,7 | 32,8 | 2,3 | 0 |
| BA033_HD_BdTR9k | BdTR9k | 400,2 | 201,3 | 104,5 | 29,4 |  | BA156_HW_Bis1 | Bis1 | 577,5 | 32,3 | 10,6 | 1,1 |
| BA363_CD_BdTR9k | BdTR9k | 514,8 | 390,2 | 585,7 | 209,9 |  | BA385_CW_Bis1 | Bis1 | 237,8 | 29,9 | 3,4 | 1,7 |
| BA410_CD_BdTR9k | BdTR9k | 454,2 | 259,8 | 136,3 | 52,7 |  | BA390_CW_Bis1 | Bis1 | 276,9 | 51,3 | 4,2 | 0,6 |
| BA110_HD_Bis1 | Bis1 | 606,4 | 739,8 | 1165,4 | 357,5 |  | BA147_HW_Kah1 | Kah1 | 167,3 | 41,4 | 3,1 | 0 |
| BA142_HD_Bis1 | Bis1 | 544,7 | 393,2 | 352,6 | 133,2 |  | BA169_HW_Kah1 | Kah1 | 218 | 68,9 | 8,6 | 1,5 |
| BA373_CD_Bis1 | Bis1 | 455,8 | 282,8 | 276,4 | 64,5 |  | BA356_CW_Kah1 | Kah1 | 135 | 47,9 | 3 | 0,8 |
| BA519_CD_Bis1 | Bis1 | 288,5 | 152,2 | 181,6 | 43,3 |  | BA420_CW_Kah1 | Kah1 | 254,1 | 46,2 | 11,5 | 3,2 |
| BA046_HD_Kah1 | Kah1 | 251,7 | 91,2 | 14,7 | 1,5 |  | BA047_HW_Kah5 | Kah5 | 280,1 | 28,2 | 5,6 | 0,9 |
| BA168_HD_Kah1 | Kah1 | 184,9 | 185,4 | 223,9 | 41 |  | BA158_HW_Kah5 | Kah5 | 268,7 | 128,5 | 16 | 3,1 |
| BA382_CD_Kah1 | Kah1 | 552,3 | 517,7 | 470,1 | 81,4 |  | BA399_CW_Kah5 | Kah5 | 351,8 | 34 | 19,5 | 7,3 |
| BA495_CD_Kah1 | Kah1 | 552,7 | 775 | 912,6 | 139,7 |  | BA482_CW_Kah5 | Kah5 | 315,1 | 30,4 | 5,6 | 2 |
| BA048_HD_Kah5 | Kah5 | 401,4 | 219,2 | 144,2 | 44,3 |  | BA141_HW_Koz1 | Koz1 | 323,6 | 22,5 | 1 | 1 |
| BA095_HD_Kah5 | Kah5 | 386,8 | 648,6 | 480,8 | 58,5 |  | BA157_HW_Koz1 | Koz1 | 353 | 68,7 | 9,5 | 0,6 |
| BA401_CD_Kah5 | Kah5 | 444,7 | 708 | 526,8 | 88,6 |  | BA501_CW_Koz1 | Koz1 | 415,6 | 43,4 | 4,3 | 2,9 |
| BA489_CD_Kah5 | Kah5 | 498,2 | 867 | 776 | 154,1 |  | BA511_CW_Koz1 | Koz1 | 397,2 | 79,4 | 31,8 | 10,6 |
| BA127_HD_Koz1 | Koz1 | 337,2 | 319,5 | 341,6 | 41,9 |  | BA057_HW_Koz3 | Koz3 | 325,3 | 18,2 | 1,3 | 0 |
| BA132_HD_Koz1 | Koz1 | 377,6 | 429,1 | 638,9 | 75,8 |  | BA074_HW_Koz3 | Koz3 | 435,3 | 20,5 | 2 | 1,1 |
| BA395_CD_Koz1 | Koz1 | 458,4 | 415,4 | 127,4 | 27 |  | BA411_CW_Koz3 | Koz3 | 283,6 | 28,8 | 15,3 | 3,7 |
| BA507_CD_Koz1 | Koz1 | 505,3 | 585,8 | 388 | 84,8 |  | BA484_CW_Koz3 | Koz3 | 332,2 | 47,8 | 9,3 | 2,3 |
| BA081_HD_Koz3 | Koz3 | 523,6 | 592,8 | 892,7 | 120 |  | BA026_HW_Ron2 | Ron2 | 218 | 122,8 | 15,3 | 1,7 |
| BA388_CD_Koz3 | Koz3 | 1094,8 | 1066,9 | 5661,6 | 1244,7 |  | BA151_HW_Ron2 | Ron2 | 350,5 | 38,4 | 7 | 1,5 |
| BA467_CD_Koz3 | Koz3 | 1201,8 | 1447,1 | 4545,9 | 1203 |  | BA409_CW_Ron2 | Ron2 | 296,4 | 36,6 | 73,2 | 32,9 |
| BA089_HD_Koz-3 | Koz3 | 773,9 | 1189,3 | 2664,1 | 681,1 |  | BA431_CW_Ron2 | Ron2 | 259,2 | 15,4 | 6,5 | 4 |
| BA035_HD_Ron2 | Ron2 | 1429,9 | 1947,5 | 1398,8 | 364,2 |  |  |  |  |  |  |  |
| BA379_CD_Ron | Ron2 | 264 | 166 | 74,8 | 15,9 |  |  |  |  |  |  |  |
| BA432_CD_Ron2 | Ron2 | 274,6 | 254,9 | 202,5 | 41,2 |  |  |  |  |  |  |  |

**Supplementary table 3**. *Brachypodium distachyon* climate data. Values of 19 current climate parameters retrieved from worldclim for the sampled localities of the studied *B. distachyon* ecotypes. PCA1 and PCA2, coordinate values of the first and second PCA axes obtained from the climate PC analysis. Climate, climatic class of the *B. distachyon* ecotypes classified according to their PCA1 values (cold:> 2.5; mesic: (-2.5) – (2.5); warm: < -2.5; see Supplementary Figure S6).

| # | ecotype | longitude | latitude | altitude | bio1 | bio2 | bio3 | bio4 | bio5 | bio6 | bio7 | bio8 | bio9 | bio10 | bio11 | bio12 | bio13 | bio14 | bio15 | bio16 | bio17 | bio18 | bio19 | PCA1 | Climate |
| --- | --- | --- | --- | --- | --- | --- | --- | --- | --- | --- | --- | --- | --- | --- | --- | --- | --- | --- | --- | --- | --- | --- | --- | --- | --- |
| 1 | ABR2 | 3.3 | 43.65 | 265 | 13.3 | 10 | 3.7 | 577.4 | 27.9 | 1.3 | 26.6 | 9.9 | 20.8 | 20.8 | 6 | 707 | 85 | 29 | 23 | 218 | 126 | 126 | 188 | 2.5793 | Cold |
| 2 | ABR3 | 0.07311 | 42.1805 | 798 | 10.1 | 10 | 3.8 | 566.4 | 24.6 | -1.5 | 26.1 | 12.1 | 3 | 17.5 | 3 | 688 | 79 | 40 | 18 | 206 | 143 | 162 | 143 | 4.8611 | Cold |
| 3 | ABR4 | 0.7168 | 42.26265 | 932 | 9.9 | 9.5 | 3.6 | 582.3 | 24.2 | -1.6 | 25.8 | 11.8 | 2.6 | 17.4 | 2.6 | 873 | 96 | 53 | 18 | 262 | 176 | 227 | 176 | 5.6901 | Cold |
| 4 | ABR5 | -0.58 | 42.581 | 986 | 8.7 | 9.8 | 3.9 | 540.9 | 22.6 | -2.4 | 25 | 10.5 | 15.8 | 15.8 | 2 | 842 | 90 | 48 | 15 | 237 | 177 | 184 | 208 | 5.3953 | Cold |
| 5 | ABR6 | -2.203 | 42.581 | 557 | 12.1 | 10.1 | 3.9 | 557.9 | 26.6 | 0.9 | 25.7 | 8.8 | 19.3 | 19.3 | 5.1 | 668 | 71 | 36 | 18 | 191 | 131 | 131 | 174 | 3.4004 | Cold |
| 6 | ABR8 | 11.319695 | 43.314569 | 300 | 13.7 | 9 | 3.4 | 600.3 | 28.7 | 2.5 | 26.2 | 10.5 | 21.6 | 21.6 | 6.4 | 757 | 101 | 29 | 29 | 262 | 117 | 117 | 203 | 2.1759 | Mesic |
| 7 | Adi10 | 38.352277 | 38.770694 | 839 | 13.6 | 9.8 | 2.7 | 936.7 | 33.4 | -2.8 | 36.2 | 12.2 | 25.3 | 25.3 | 1.3 | 459 | 63 | 2 | 57 | 172 | 13 | 25 | 161 | -1.7567 | Mesic |
| 8 | Adi12 | 38.352277 | 38.770694 | 839 | 13.6 | 9.8 | 2.7 | 936.7 | 33.4 | -2.8 | 36.2 | 12.2 | 25.3 | 25.3 | 1.3 | 459 | 63 | 2 | 57 | 172 | 13 | 25 | 161 | -1.7567 | Mesic |
| 9 | Adi2 | 38.352277 | 38.770694 | 839 | 13.6 | 9.8 | 2.7 | 936.7 | 33.4 | -2.8 | 36.2 | 12.2 | 25.3 | 25.3 | 1.3 | 459 | 63 | 2 | 57 | 172 | 13 | 25 | 161 | -1.7567 | Mesic |
| 10 | Bd1-1 | 28.251 | 38.417 | 644 | 13.8 | 11.5 | 3.8 | 660.1 | 30.2 | 0.7 | 29.5 | 5.6 | 22 | 22.4 | 5.6 | 698 | 146 | 8 | 72 | 360 | 38 | 39 | 360 | -0.5973 | Mesic |
| 11 | Bd18-1 | 33.730025 | 39.36784 | 1057 | 10.4 | 10.9 | 3.4 | 747.4 | 27.4 | -4.3 | 31.7 | 0.4 | 19.4 | 19.6 | 0.4 | 445 | 63 | 6 | 48 | 161 | 29 | 49 | 161 | 0.7309 | Mesic |
| 12 | Bd2-3 | 44.403075 | 33.760883 | 40 | 22.7 | 15.2 | 3.8 | 877.5 | 43.7 | 4.5 | 39.2 | 12.7 | 33.7 | 33.7 | 11.3 | 171 | 31 | 0 | 86 | 88 | 0 | 0 | 87 | -5.7177 | Warm |
| 13 | Bd21ctrl | 44.535 | 36.766 | 1089 | 15 | 12.1 | 3.1 | 961.1 | 36 | -2.8 | 38.8 | 3.6 | 27 | 27 | 2.6 | 728 | 136 | 0 | 86 | 385 | 3 | 3 | 349 | -3.5166 | Warm |
| 14 | Bd21-3 | 44.535 | 36.766 | 1089 | 15 | 12.1 | 3.1 | 961.1 | 36 | -2.8 | 38.8 | 3.6 | 27 | 27 | 2.6 | 728 | 136 | 0 | 86 | 385 | 3 | 3 | 349 | -3.5166 | Warm |
| 15 | Bd3-1 | 44.535 | 36.766 | 1089 | 15 | 12.1 | 3.1 | 961.1 | 36 | -2.8 | 38.8 | 3.6 | 27 | 27 | 2.6 | 728 | 136 | 0 | 86 | 385 | 3 | 3 | 349 | -3.5166 | Warm |
| 16 | Bd30-1 | -3.558733 | 36.990489 | 810 | 14.7 | 11.4 | 3.8 | 603.2 | 31.7 | 2.2 | 29.5 | 8.4 | 22.9 | 22.9 | 7.5 | 467 | 63 | 5 | 54 | 181 | 24 | 24 | 175 | -0.4576 | Mesic |
| 17 | BdTR10c | 31.884911 | 37.778233 | 1448 | 9.4 | 11.5 | 3.5 | 750 | 27.2 | -5.6 | 32.8 | -0.3 | 18.6 | 18.9 | -0.3 | 522 | 77 | 10 | 49 | 209 | 36 | 52 | 209 | 0.8878 | Mesic |
| 18 | BdTR11g | 27.477 | 41.422 | 89 | 13 | 11.3 | 3.7 | 666.2 | 29.5 | -0.3 | 29.8 | 6.2 | 20.9 | 21.5 | 4.5 | 598 | 78 | 18 | 37 | 227 | 74 | 86 | 202 | 1.0333 | Mesic |
| 19 | BdTR11i | 28.040197 | 39.738164 | 229 | 13.7 | 10.9 | 3.7 | 683.7 | 29.8 | 0.5 | 29.3 | 7 | 21.9 | 22.4 | 5 | 652 | 108 | 11 | 56 | 284 | 48 | 49 | 282 | 0.0435 | Mesic |
| 20 | BdTR1i | 28.583 | 38.093 | 956 | 12.6 | 11.5 | 3.7 | 697.5 | 29.7 | -0.9 | 30.6 | 3.9 | 21.3 | 21.6 | 3.9 | 748 | 146 | 10 | 68 | 376 | 43 | 49 | 376 | -0.2852 | Mesic |
| 21 | BdTR2b | 31.331114 | 40.082097 | 894 | 10.9 | 10.5 | 3.4 | 714.2 | 27.4 | -2.9 | 30.3 | 1.5 | 19.6 | 19.8 | 1.5 | 488 | 61 | 17 | 34 | 159 | 60 | 85 | 159 | 1.5604 | Mesic |
| 22 | BdTR2g | 32.985 | 40.394 | 1531 | 7.3 | 10.5 | 3.3 | 722 | 23.9 | -7.1 | 31 | -2.4 | 16 | 16.2 | -2.4 | 623 | 84 | 20 | 42 | 221 | 67 | 97 | 221 | 2.6038 | Cold |
| 23 | BdTR5i | 32.985367 | 40.393647 | 1531 | 7.3 | 10.5 | 3.3 | 722 | 23.9 | -7.1 | 31 | -2.4 | 16 | 16.2 | -2.4 | 623 | 84 | 20 | 42 | 221 | 67 | 97 | 221 | 2.6038 | Cold |
| 24 | BdTR9k | 30.788631 | 39.75295 | 900 | 10.7 | 11 | 3.5 | 729.1 | 27.8 | -3.5 | 31.3 | 9.8 | 19.5 | 19.8 | 1.1 | 419 | 52 | 11 | 37 | 136 | 45 | 67 | 134 | 1.1200 | Mesic |
| 25 | Bis1 | 41.015083 | 37.876556 | 608 | 16.5 | 13.1 | 3.3 | 928.6 | 38.9 | -0.6 | 39.5 | 9.7 | 28.3 | 28.3 | 4.5 | 548 | 83 | 1 | 70 | 237 | 6 | 10 | 219 | -3.6626 | Warm |
| 26 | Kah1 | 38.533 | 37.734 | 657 | 16.9 | 11 | 2.9 | 914.8 | 37.3 | 0.6 | 36.7 | 5.3 | 28.6 | 28.6 | 5.3 | 586 | 108 | 1 | 77 | 291 | 7 | 10 | 291 | -3.5423 | Warm |
| 27 | Kah5 | 38.533 | 37.734 | 657 | 16.9 | 11 | 2.9 | 914.8 | 37.3 | 0.6 | 36.7 | 5.3 | 28.6 | 28.6 | 5.3 | 586 | 108 | 1 | 77 | 291 | 7 | 10 | 291 | -3.5423 | Warm |
| 28 | Koz1 | 41.61 | 38.152 | 819 | 15.3 | 12.1 | 3 | 951.9 | 37.2 | -2.1 | 39.3 | 8.3 | 27.4 | 27.4 | 3.1 | 703 | 104 | 1 | 69 | 303 | 10 | 10 | 283 | -3.2693 | Warm |
| 29 | Koz3 | 41.61 | 38.152 | 819 | 15.3 | 12.1 | 3 | 951.9 | 37.2 | -2.1 | 39.3 | 8.3 | 27.4 | 27.4 | 3.1 | 703 | 104 | 1 | 69 | 303 | 10 | 10 | 283 | -3.2693 | Warm |
| 30 | RON2 | -0.963 | 42.781 | 956 | 8.9 | 9.9 | 3.9 | 539.5 | 22.9 | -2.2 | 25.1 | 3.1 | 16 | 16 | 2.2 | 952 | 101 | 52 | 16 | 274 | 190 | 190 | 257 | 5.4782 | Cold |

| Variable contribution | PCA1 | PCA2 |
| --- | --- | --- |
| bio1 | 7.04157791 | 2.8336316 |
| bio2 | 5.1431788 | 0.23831544 |
| bio3 | 3.59658313 | 1.20932109 |
| bio4 | 7.92629573 | 0.71530785 |
| bio5 | 9.29173338 | 0.57639096 |
| bio6 | 0.48821208 | 6.97369089 |
| bio7 | 8.7044993 | 0.30756729 |
| bio8 | 0.03627953 | 10.374377 |
| bio9 | 8.12122146 | 0.14333639 |
| bio10 | 9.05581757 | 0.73901616 |
| bio11 | 1.08408395 | 6.54349079 |
| bio12 | 2.28358537 | 11.4483415 |
| bio13 | 0.23276403 | 18.6365656 |
| bio14 | 8.64758564 | 0.05494295 |
| bio15 | 9.03019188 | 1.48194295 |
| bio16 | 0.49621767 | 18.6220566 |
| bio17 | 8.61189114 | 0.08756497 |
| bio18 | 9.14009127 | 0.07915297 |
| bio19 | 1.06819015 | 18.9349869 |

**Supplementary Table 4**. Sampled dehydrin sequences from grass species related to *Brachypodium*. The accession code and the protein name correspond to those indicated in Phytozome and Genbank.

| *Aegilops tauschii* | | *Hordeum vulgare* | | *Oryza sativa* | | *Sorghum bicolor* | | *Zea mays* | |
| --- | --- | --- | --- | --- | --- | --- | --- | --- | --- |
| Accession | **Name** | **Accession** | **Name** | **Accession** | **Name** | **Accession** | **Name** | **Accession** | **Name** |
| AET6Gv20653900 | DHNAtau 1 | HORVU5Hr1G092120 | DHNHvul1a | LOC_Os02g44870 | DHNOsat 1 | Sobic.001G149500 | DHNSbic 10 | GRMZM2G147014 | DHNZmays 1 |
| AET5Gv20866800 | DHNAtau 2 | HORVU5Hr1G092160 | DHNHvul 1b | LOC_Os11g26570 | DHNOsat 2 | Sobic.004G286600 | DHNSbic 2 | GRMZM2G373522 | DHNZmays 2 |
| AET4Gv20132600 | DHNAtau 3 | HORVU5Hr1G092100 | DHNHvul 2a | LOC_Os11g26780 | DHNOsat 3a | Sobic.009G116700 | DHNSbic 3 | Zm00001d010094 | DHNZmays 3 |
| AET6Gv20864900 | DHNAtau 4 | HORVU5Hr1G092150 | DHNHvul 2b | LOC_Os11g26790 | DHNOsat 3b | Sobic.010G041900 | DHNSbic 4 | GRMZM2G052364 | DHNZmays 4 |
| AET5Gv20866700 | DHNAtau 5 | HORVU6Hr1G084070 | DHNHvul 3 | LOC_Os01g50700 | DHNOsat 4 | Sobic.003G270200 | DHNSbic 5 | GRMZM2G098750 | DHNZmays 5 |
| AET6Gv20865700 | DHNAtau 5 | HORVU6Hr1G084010 | DHNHvul 7 | LOC_Os03g45280 | DHNOsat 13 |  |  | GRMZM2G169372 | DHNZmays 10a |
| AET6Gv20866400 | DHNAtau 6 | HORVU6Hr1G064620 | DHNHvul 8 |  |  |  |  | GRMZM2G448511 | DHNZmays 10b |
| AET6Gv20866000 | DHNAtau 7 | AY681974 | DHNHvul 13 |  |  |  |  |  |  |
| AET3Gv20620600 | DHNAtau 8 |  |  |  |  |  |  |  |  |

**Supplementary Table 5**. Chromosomal location of Bdhn genes across the four studied Brachypodium species and genomes. Chr, chromosome number (B. distachyon: Bd1-Bd5; B. hybridum subgenome D: BhD1-BhD5; B. hybridum subgenome S: BhS1-BhS10; B. stacei: Bs1-Bs10; B. sylvaticum: Bsy1-Bsy9). Lenghts and positions correspond to the respective reference genomes.

|  |  | ***Bdhn*1a** | ***Bdhn*1b** | ***Bdhn*2** | ***Bdhn*3** | ***Bdhn*4** | ***Bdhn*5** | ***Bdhn*6** | ***Bdhn*7** | ***Bdhn*8** | ***Bdhn*9** | ***Bdhn*10** |
| --- | --- | --- | --- | --- | --- | --- | --- | --- | --- | --- | --- | --- |
| ***B.distachyon*** | **Chr** | Bd5 |  | Bd3 | Bd1 | Bd 4 | Bd 4 | Bd 4 | Bd 3 | Bd 3 | Bd 2 | Bd 1 |
|  | **Lenght** | 28630136 |  | 59640145 | 75071545 | 48594894 | 48594894 | 48594894 | 59640145 | 59640145 | 59130575 | 75071545 |
|  | **from** | 14358126 |  | 52061272 | 33400971 | 26400080 | 26401854 | 22188216 | 45300502 | 45290875 | 47751383 | 10098645 |
|  | **to** | 14359531 |  | 52062769 | 33402000 | 26400807 | 26402677 | 22189942 | 45301669 | 45291925 | 47757368 | 10100931 |
|  | **direction** | forward |  | forward | reverse | reverse | reverse | forward | reverse | forward | forward | reverse |
| ***B.hybridum*D** | **Chr** | BhD5 |  | BhD3 | BhD1 |  | BhD 4 | BhD 4 | BhD3 | BhD 3 | BhD 2 | BhD 1 |
|  | **Lenght** | 28673805 |  | 59422649 | 73190849 |  | 48381198 | 48381198 | 59422649 | 59422649 | 59597844 | 73190849 |
|  | **from** | 14493652 |  | 51909664 | 32658384 |  | 25616401 | 21335117 | 45017036 | 45007503 | 47969138 | 9743153 |
|  | **to** | 14494867 |  | 51910828 | 32659379 |  | 25617226 | 21336708 | 45018083 | 45008367 | 47975373 | 9743841 |
|  | **direction** | forward |  | forward | reverse |  | reverse | forward | reverse | forward | forward | reverse |
| ***B. hybridum*S** | **Chr** | BhS9 |  | BhS4 | BhS7 |  | BhS5 | BhS5 | BhS4 | BhS4 | BhS1 | BhS2 |
|  | **Lenght** | 21007308 |  | 25447193 | 21638549 |  | 23591727 | 23591727 | 25447193 | 25447193 | 30608744 | 28346489 |
|  | **from** | 7214663 |  | 7269373 | 14706419 |  | 3849103 | 7640493 | 13819476 | 13842937 | 10890765 | 19034663 |
|  | **to** | 7216004 |  | 7270541 | 14707427 |  | 3849990 | 7642071 | 13820451 | 13843854 | 10897704 | 19035490 |
|  | **direction** | forward |  | reverse | forward |  | forward | forward | forward | reverse | reverse | forward |
| ***B. stacei*** | **Chr** | Bs9 |  | Bs4 | Bs7 |  | Bs5 | Bs5 | Bs4 | Bs4 | Bs1 | Bs2 |
|  | **Lenght** | 20576529 |  | 24645555 | 20893312 |  | 23048618 | 23048618 | 24645555 | 24645555 | 30086066 | 27792411 |
|  | **from** | 9210852 |  | 6997868 | 14026099 |  | 3806913 | 6372578 | 13156720 | 13537514 | 10860306 | 18652003 |
|  | **to** | 9212190 |  | 6999029 | 14026708 |  | 3849990 | 6374159 | 13157693 | 13538422 | 10866472 | 18652813 |
|  | **direction** | forward |  | reverse | forward |  | forward | reverse | forward | forward | reverse | forward |
| ***B. sylvaticum*** | **Chr** | Bsy9 | Bsy9 | Bsy4 | Bsy7 |  | Bsy5 | Bsy4 | Bsy4 | Bsy4 | Bsy1 | Bsy2 |
|  | **Lenght** | 31923712 | 31923712 | 38747968 | 22318590 |  | 48035605 | 38747968 | 38747968 | 38747968 | 52666873 | 42817455 |
|  | **from** | 13763612 | 14190187 | 9798501 | 14062257 |  | 26543391 | 21623366 | 19478535 | 19507464 | 17853540 | 25255158 |
|  | **to** | 13765006 | 14191050 | 9799643 | 14062972 |  | 26543956 | 21625489 | 19479154 | 19508169 | 17853540 | 25256158 |
|  | **direction** | forward | reverse | reverse | forward |  | reverse | forward | forward | reverse | reverse | forward |

**Supplementary Table 6**. **(a)**Non-synonymous (dN) and synonymous (dS) substitution rate values and dN/dS (ω) ratio of Bdhn genes in the studied Brachypodium species and genomes. Abbreviatures of species and reference genomes: BD, B. distachyon; BHD, B. hybridum D-subgenome; BS, B. stacei; BHS, B. hybridum S-subgenome; BSY, B. sylvaticum. DHN, orthologous dehydrin genes from close outgroup grass species (Sbic, Sorghum bicolor; Zmays, Zea mays) used to estimate the dN and dS rates of the Brachypodium Bdhn genes.

| ***Bdhn* genes** | **DHN** | **dN** | **dS** | **ω** |
| --- | --- | --- | --- | --- |
| BD_*Bdhn*1a | DHNSbic2 | 0.092742 | 1.472219 | 0.062995 |
| BD_*Bdhn*2 | DHNSbic2 | 0.081048 | 2.395785 | 0.033829 |
| BHD_*Bdhn*1a | DHNSbic2 | 0.09344 | 1.447833 | 0.064538 |
| BHD_*Bdhn*2 | DHNSbic2 | 0.07907 | 1.564455 | 0.050541 |
| BHS_*Bdhn*1a | DHNSbic2 | 0.085076 | 1.790013 | 0.047528 |
| BHS_*Bdhn*2 | DHNSbic2 | 0.08452 | 1.372253 | 0.061592 |
| BS_*Bdhn*1a | DHNSbic2 | 0.085135 | 2.311014 | 0.036839 |
| BS_*Bdhn*2 | DHNSbic2 | 0.0845 | 1.378765 | 0.061286 |
| BSY_*Bdhn*1a | DHNSbic2 | 0.08266 | 1.550495 | 0.053312 |
| BSY_*Bdhn*1b | DHNSbic2 | 0.089337 | 2.89104 | 0.030901 |
| BSY_*Bdhn*2 | DHNSbic2 | 0.084348 | 1.375851 | 0.061306 |
| BD_*Bdhn*4 | DHNSbic3 | 0.145795 | 2.252359 | 0.06473 |
| BD_*Bdhn*5 | DHNSbic3 | 0.086338 | 1.28195 | 0.067349 |
| BD_*Bdhn*7 | DHNSbic3 | 0.034315 | 2.297179 | 0.014938 |
| BD_*Bdhn*8 | DHNSbic3 | 0.06164 | 2.253262 | 0.027356 |
| BHD_*Bdhn*5 | DHNSbic3 | 0.086338 | 1.28195 | 0.067349 |
| BHD_*Bdhn*7 | DHNSbic3 | 0.034315 | 2.297179 | 0.014938 |
| BHD_*Bdhn*8 | DHNSbic3 | 0.061509 | 2.239681 | 0.027463 |
| BHS_*Bdhn*5 | DHNSbic3 | 0.078414 | 3.069189 | 0.025549 |
| BHS_*Bdhn*7 | DHNSbic3 | 0.034086 | 2.250424 | 0.015147 |
| BHS_*Bdhn*8 | DHNSbic3 | 0.045006 | 2.259038 | 0.019923 |
| BS_*Bdhn*5 | DHNSbic3 | 0.078442 | 2.931374 | 0.026759 |
| BS_*Bdhn*7 | DHNSbic3 | 0.034086 | 2.250424 | 0.015147 |
| BS_*Bdhn*8 | DHNSbic3 | 0.039525 | 2.259377 | 0.017494 |
| BSY_*Bdhn*5 | DHNSbic3 | 0.099529 | 2.07334 | 0.048004 |
| BSY_*Bdhn*7 | DHNSbic3 | 0.034345 | 2.301785 | 0.014921 |
| BSY_*Bdhn*8 | DHNSbic3 | 0.064693 | 1.987505 | 0.03255 |
| BD_*Bdhn*6 | DHNSbic4 | 0.26622 | 1.868913 | 0.142447 |
| BHD_*Bdhn*6 | DHNSbic4 | 0.265666 | 2.07041 | 0.128315 |
| BHS_*Bdhn*6 | DHNSbic4 | 0.264243 | 2.127335 | 0.124213 |
| BS_*Bdhn*6 | DHNSbic4 | 0.264645 | 1.93607 | 0.136692 |
| BSY_*Bdhn*6 | DHNSbic4 | 0.266181 | 2.006112 | 0.132685 |
| BD_*Bdhn*9 | DHNSbic5 | 0.146505 | 1.884926 | 0.077724 |
| BHD_*Bdhn*9 | DHNSbic5 | 0.145158 | 1.939981 | 0.074824 |
| BHS_*Bdhn*9 | DHNSbic5 | 0.152573 | 2.459047 | 0.062046 |
| BS_*Bdhn*9 | DHNSbic5 | 0.152573 | 2.459047 | 0.062046 |
| BSY_*Bdhn*9 | DHNSbic5 | 0.145048 | 2.488954 | 0.058277 |
| BD_*Bdhn*3 | DHNZmays3 | 0.203305 | 2.088461 | 0.097347 |
| BHD_*Bdhn*3 | DHNZmays3 | 0.202558 | 2.292995 | 0.088338 |
| BHS_*Bdhn*3 | DHNZmays3 | 0.213816 | 2.501106 | 0.085489 |
| BS_*Bdhn*3 | DHNZmays3 | 0.201682 | 2.489685 | 0.081007 |
| BSY_*Bdhn*3 | DHNZmays3 | 0.223666 | 2.483027 | 0.090078 |
| BD_*Bdhn*10 | DHNSbic10 | 0.059922 | 2.734014 | 0.021917 |
| BHD_*Bdhn*10 | DHNSbic10 | 0.059922 | 2.734014 | 0.021917 |
| BHS_*Bdhn*10 | DHNSbic10 | 0.059736 | 1.308958 | 0.045636 |
| BS_*Bdhn*10 | DHNSbic10 | 0.059736 | 1.308958 | 0.045636 |
| BSY_*Bdhn*10 | DHNSbic10 | 0.059853 | 1.136323 | 0.052673 |

**(b).** Values of non-synonymous (dN) and synonymous (dS) substitutions, their ratio dS/dN (ω), for duplicated Bdhn genes across the studied species and genomes of Brachypodium.

| ***Bdhn* gene pair** | **dN** | **dS** | **ω** | **Mode of duplication** |
| --- | --- | --- | --- | --- |
| BD *Bdhn* 1/ *Bdhn* 2 | 0.017505 | 0.257996 | 0.067852 | segmental |
| BHD *Bdhn* 1/*Bdhn* 2 | 0.01557 | 0.314666 | 0.049481 | segmental |
| BS *Bdhn* 1/*Bdhn* 2 | 0.018561 | 0.167476 | 0.110829 | segmental |
| BHS *Bdhn*1/*Bdhn*2 | 0.018554 | 0.15123 | 0.122685 | segmental |
| BSY *Bdhn* 1a/*Bdhn* 1b | 0.02137 | 0.229128 | 0.093265 | segmental |
| BSY *Bdhn* 1a/ *Bdhn* 2 | 0.013553 | 0.381983 | 0.035481 | segmental |
| BSY *Bdhn* 1b/ *Bdhn* 2 | 0.020532 | 0.262587 | 0.078193 | segmental |
| BD Bhdn 4/Bhdn 5 | 0.058539 | 1.282730 | 0.045637 | tandem |
| BD *Bdhn* 7/*Bdhn* 8 | 0.025959 | 0.315468 | 0.082288 | tandem |
| BHD *Bdhn*7/*Bdhn*8 | 0.025909 | 0.474996 | 0.054546 | tandem |
| BS *Bdhn* 7/*Bdhn* 8 | 0.005075 | 0.258587 | 0.019626 | tandem |
| BHS *Bdhn*7/*Bdhn*8 | 0.010198 | 0.258743 | 0.039412 | tandem |
| BSY *Bdhn*7/*Bdhn*8 | 0.025979 | 1.769220 | 0.014684 | tandem |

**Supplementary Table 7.** Topological congruence tests between **(a)** the *B. distachyon* nuclear species tree (Gordon et al. 2017) and **(b)** the *B. distachyon* plastome tree (Sancho et al. 2018) *versus* the *B. distachyon* dehydrin *Bdhn* tree. Test(s) were performed for significance of likelihood-score differences. KH: Kishino-Hasegawa test using normal approximation, two-tailed test**.** SH: Shimodaira-Hasegawa test using RELL bootstrap (one-tailed test). AU: Shimodaira Approximately Unbiased test**.** Values for KH/SH/AU tests are P values for the null hypothesis of no difference between trees**.** *the null hypothesis is accepted. Number of bootstrap replicates = 1,000,000.

|  | **KH test** | | | | | **SH** | | |
| --- | --- | --- | --- | --- | --- | --- | --- | --- |
| a) | **-lnL** | **Diff'-lnL** | **s.d.** | **T** | **P** | **SH-test** | **wtd-SH** | **AU** |
| **Tree1= Brachy nuclear tree (best)** | 46255.5731 | (best) |  |  |  |  |  |  |
| **Tree2= Brachy *Bdhn* tree** | 54391.9775 | 8136.40442 | 202.241 | 40.231 | <0.0001* | 0.0000* | 0.0000* | ~0* |
| **Tree1=Brachy *Bdhn* tree (best)** | 5898.89581 | (best) |  |  |  |  |  |  |
| **Tree2=Brachy nuclear tree** | 6170.77777 | 271.88196 | 59.738 | 4.551 | <0.0001* | 0.0000* | 0.0000* | ~0* |
|  | * P < 0.05 |  |  |  |  |  |  |  |
| (b) |  |  |  |  |  |  |  |  |
| **Tree1= Brachy plastome tree (best)** | 1442.4868 | (best) |  |  |  |  |  |  |
| **Tree2= Brachy *Bdhn* tree** | 2334.75366 | 892.26686 | 52.856 | 16.881 | <0.0001* | 0.0000* | 0.0000* | ~0* |
| **Tree1= Brachy *Bdhn* tree (best)** | 5898.89581 | (best) |  |  |  |  |  |  |
| **Tree2= Brachy plastome tree** | 6268.97012 | 370.07431 | 91.549 | 4.042 | 0.0001* | 0.0003* | 0.0003* | ~0* |
|  | * P < 0.05 |  |  |  |  |  |  |  |

**Supplementary Table 8**. Linear model (lm) regression analysis for comparative differential gene expressions of dehydrin *Bdhn* genes in the studied *B. distachyon* ecotypes. W (watered) and D (dry) conditions. Significant p-values (p≤ 0.05*; 0.01**; 0.001***).

| W+D | ***Bdhn*1a ~ *Bdhn*2** | ***Bdhn*1a ~ *Bdhn*3** | ***Bdhn*1a ~ *Bdhn*7** | ***Bdhn*2 ~ *Bdhn*3** | ***Bdhn*2 ~ *Bdhn*7** | ***Bdhn*3 ~ *Bdhn*7** |
| --- | --- | --- | --- | --- | --- | --- |
| Median | -11.12 | -14.77 | -14.05 | -68.64 | -72.05 | -37.35 |
| Residual standard error: | 134.6 | 169.5 | 156.3 | 220.8 | 210.7 | 237 |
| F-statistic: | 368 | 148.3 | 214.6 | 344.9 | 401 | 1840 |
| p-value: | < 2.20E-16*** | < 2.20E-16*** | < 2.20E-16*** | < 2.20E-16*** | < 2.20E-16*** | < 2.20E-16*** |

**Supplementary Table 9**. Comparative analysis of dehydrin genes showing upregulated expression under drought compared to watered conditions in *Brachypodium distachyon* and *Triticum aestivum*. Orthology between the differentially expressed genes in the two species was retrieved through Ensembl Plants BioMart, Ensembl Plants BLAST (*) and Blast searches using data from Galvez et al. (2019) (**).

| ***Brachypodium distachyon*** | | | ***Triticum aestivum*** | | |
| --- | --- | --- | --- | --- | --- |
| **Gene name** | **RefSeq.v3.0 Name** | **Ref.Seqv3.1 Name** | **Gene name** | **RefSeq v2.1 Name** | **Differentially expressed (Galvez et al. 2019)** |
| *Bdhn*1 | BRADI_5g10860v3 | Bradi5g10860 | DHN11-A1 | TraesCS6A02G253300 | Mild stress |
| *Bdhn*2 | BRADI_3g51200v3 | Bradi3g51200 | DHN11-B1 | TraesCS6B02G273400 |  |
|  |  |  | DHN11-D1 | TraesCS6D02G234700 |  |
| *Bdhn*3 | BRADI_1g37410v3 | Bradi1g37410 | DHN4-B1 | [TraesCS6B02G383500](https://plants.ensembl.org/triticum_aestivum/Gene/Summary?g=TraesCS6B02G383500) | Severe Stress |
|  |  |  | DHN4-D1 | [TraesCS6D02G332900](https://plants.ensembl.org/triticum_aestivum/Gene/Summary?g=TraesCS6D02G332900) | Severe Stress |
|  |  |  | DHN3-A1 | [TraesCS6A02G350600](https://plants.ensembl.org/triticum_aestivum/Gene/Summary?g=TraesCS6A02G350600) | Mild stress |
|  |  |  | DHN3-A5 | [TraesCS6A02G350800](https://plants.ensembl.org/triticum_aestivum/Gene/Summary?g=TraesCS6A02G350800) |  |
|  |  |  | DHN3-A6 | [TraesCS6A02G350700](https://plants.ensembl.org/triticum_aestivum/Gene/Summary?g=TraesCS6A02G350700) | Mild stress |
|  |  |  | DHN3-B6 | [TraesCS6B02G383600](https://plants.ensembl.org/triticum_aestivum/Gene/Summary?g=TraesCS6B02G383600) | Mild stress |
|  |  |  | DHN3-D6 | [TraesCS6D02G333200](https://plants.ensembl.org/triticum_aestivum/Gene/Summary?g=TraesCS6D02G333200) | Severe Stress |
|  |  |  | DHN3-D4 | [TraesCS6D02G333100](https://plants.ensembl.org/triticum_aestivum/Gene/Summary?g=TraesCS6D02G333100) | Severe Stress |
|  |  |  | DHN3-D8 | [TraesCS6D02G333300](https://plants.ensembl.org/triticum_aestivum/Gene/Summary?g=TraesCS6D02G333300) |  |
|  |  |  | DHN3-D9 | [TraesCS6D02G333600](https://plants.ensembl.org/triticum_aestivum/Gene/Summary?g=TraesCS6D02G333600) | Severe Stress |
| *Bdhn*7 | BRADI_3g43870v3 | Bradi3g43870 | DHN38-A1* | TraesCS5A02G424700 |  |
|  |  |  | DHN38-B1* | TraesCS5B02G426700 | Severe Stress |
|  |  |  | DHN38-D1** | TraesCS5D01G433200 |  |
|  |  |  | DHN38-A2* | TraesCS5A02G424800 |  |
|  |  |  | DHN38-B2** | TraesCS5B01G426800 | Severe Stress |
|  |  |  | DHN38-D2* | TraesCS5D02G433300 |  |

**Supplementary Table 10**. Summary statistics of 12 drought-response phenotypic traits [leaf_rwc (relative water content in leaf); leaf_wc (water content in leaf); lma (leaf mass per área); pro (leaf proline content); abvrgd (above ground biomass); blwgrd (below ground biomass); ttlmass (total mass); rmr (root mass ratio); delta13c (carbon isotope, a proxy for lifetime integrated WUE); leafc (leaf carbon content); leafn (leaf nitrogen content); cn (leaf carbon/nitrogen ratio)] in drought (D) vs watered (W) *Brachypodium distachyon* plants. (a) Kruskall-Wallis rank tests (W vs D) for each phenotypic trait. (b) comparative pairwise Wilcoxon tests in the studied *B. distachyon* ecotypes; p-values were adjusted with the Benjamini–Hochberg (BH) procedure, controlling the false discovery rate, to correct for multiple comparisons; n, number of replicates. n. s., non significant, *p≤ 0.05*; significant values are highlighted in bold.

**(a)**

| **Var** | **Leaf_rwc** | **Leaf_wc** | **Lma** | **Pro** | **abvgrd** | **blwgrd** | **ttlmas** | **rmr** | **WUE** | **leafC** | **LeafN** | **C:N** |
| --- | --- | --- | --- | --- | --- | --- | --- | --- | --- | --- | --- | --- |
| **t-test** | 197.24 | 172.97 | 166.68 | 189.78 | 190.61 | 161.24 | 181.31 | 192.97 | 192.67 | 169.61 | 188.34 | 188.8 |
| **df** | 63 | 63 | 63 | 63 | 63 | 63 | 63 | 63 | 63 | 63 | 63 | 63 |
| **p-value** | 9.46E-16*** | 3.42E-12*** | 2.62E-11*** | 1.24E-14*** | 9.32E-15*** | 1.47E-10*** | 2.17E-13*** | 4.15E-15*** | 4.60E-15*** | 1.02E-11*** | 2.02E-14*** | 7.32E-15*** |

**(b)**

| **Ecotype** | **n** | **leaf_rwc** | | | **leafwc** | | | **lma** | | | **pro** | | | **abvgrd** | | | **blwgrd** | | |
| --- | --- | --- | --- | --- | --- | --- | --- | --- | --- | --- | --- | --- | --- | --- | --- | --- | --- | --- | --- |
|  |  | **D** | **W** | **W-test** | **D** | **W** | **W-test** | **D** | **W** | **W-test** | **D** | **W** | **W-test** | **D** | **W** | **W-test** | **D** | **W** | **W-test** |
| ABR2 | 4 | **95.92** | **99.09** | ***** | **275.41** | **327.41** | ***** | **31.09** | **27.07** | ***** | **39.54** | **17.51** | ***** | 82.39 | 82.83 | n.s | 54.70 | 39.94 | n.s |
| ABR3 | 4 | **95.91** | **99.23** | ***** | 325.50 | 341.29 | n.s | 24.83 | 23.35 | n.s | **18.48** | **9.26** | ***** | 94.28 | 113.30 | n.s | 57.44 | 52.51 | n.s |
| ABR4 | 4 | **96.62** | **99.44** | ***** | **312.84** | **357.86** | ***** | 28.94 | 27.19 | n.s | **15.44** | **8.76** | ***** | **53.93** | **63.83** | ***** | **32.19** | **29.34** | ***** |
| ABR5 | 3 | 95.28 | 100.14 | n.s | 309.74 | 355.00 | n.s | 27.51 | 23.67 | n.s | 12.79 | 7.68 | n.s | 83.10 | 84.11 | n.s | 55.12 | 33.31 | n.s |
| ABR6 | 4 | **95.07** | **99.73** | ***** | **300.31** | **334.41** | ***** | **27.93** | **25.84** | ***** | **24.37** | **10.16** | ***** | 72.96 | 80.03 | n.s | **40.48** | **30.95** | ***** |
| ABR8 | 4 | **94.01** | **99.09** | ***** | 327.95 | 359.65 | n.s | **30.93** | **27.92** | ***** | **25.45** | **8.00** | ***** | **112.43** | **172.49** | ***** | **33.23** | **40.93** | ***** |
| Adi10 | 4 | **92.50** | **99.49** | ***** | **333.21** | **372.90** | ***** | **28.86** | **24.35** | ***** | **54.79** | **10.94** | ***** | **133.87** | **225.64** | ***** | 67.65 | 80.73 | n.s |
| Adi12 | 4 | **92.10** | **99.41** | ***** | **291.01** | **322.88** | ***** | **29.34** | **26.69** | ***** | **30.06** | **10.47** | ***** | **141.38** | **168.85** | ***** | **70.54** | **57.79** | ***** |
| Adi2 | 3 | 94.99 | 97.91 | n.s | 282.54 | 307.59 | n.s | 28.99 | 25.70 | n.s | 40.57 | 8.61 | n.s | 102.75 | 161.41 | n.s | 55.39 | 49.99 | n.s |
| Bd1-1 | 2 | 96.65 | 99.49 | n.s | 318.06 | 340.96 | n.s | 27.89 | 26.05 | n.s | 22.48 | 8.68 | n.s | 69.95 | 70.85 | n.s | 32.68 | 28.03 | n.s |
| Bd18-1 | 2 | 92.46 | 98.01 | n.s | 305.47 | 330.20 | n.s | 29.18 | 26.35 | n.s | 26.56 | 9.00 | n.s | 127.49 | 166.53 | n.s | 60.50 | 52.55 | n.s |
| Bd21ctrl | 4 | **94.72** | **98.53** | ***** | 348.66 | 370.79 | n.s | 29.95 | 28.19 | n.s | **16.88** | **7.80** | ***** | 104.63 | 104.94 | n.s | **48.95** | **34.90** | ***** |
| Bd21-3 | 3 | 88.24 | 99.96 | n.s | 300.78 | 333.15 | n.s | 34.40 | 29.66 | n.s | 46.00 | 7.99 | n.s | 132.92 | 166.58 | n.s | 76.00 | 57.59 | n.s |
| Bd2-3 | 4 | **94.68** | **99.54** | ***** | 301.70 | 323.22 | n.s | **29.30** | **27.49** | ***** | **24.56** | **7.18** | ***** | **109.68** | **142.81** | ***** | 52.55 | 48.69 | n.s |
| Bd30-1 | 4 | **91.94** | **98.74** | ***** | **290.94** | **360.88** | ***** | **29.98** | **24.18** | ***** | **39.52** | **8.27** | ***** | **108.47** | **150.05** | ***** | **55.83** | **70.51** | ***** |
| Bd3-1 | 3 | **87.95** | **98.93** | ***** | **314.60** | **346.29** | ***** | **30.34** | **26.74** | ***** | **39.47** | **11.38** | ***** | **147.08** | **202.11** | ***** | 64.45 | 66.88 | n.s |
| BdTR10c | 4 | 92.18 | 99.85 | n.s | 292.36 | 346.45 | n.s | 28.54 | 24.40 | n.s | 62.13 | 9.26 | n.s | 122.84 | 179.83 | n.s | 54.84 | 78.85 | n.s |
| BdTR11g | 4 | **92.67** | **99.17** | ***** | **303.40** | **348.85** | ***** | 30.29 | 28.51 | n.s | **26.36** | **6.46** | ***** | **118.32** | **162.40** | ***** | 58.96 | 59.50 | n.s |
| BdTR11i | 4 | **92.16** | **98.87** | ***** | **301.41** | **356.48** | ***** | **31.49** | **26.81** | ***** | **44.23** | **7.77** | ***** | **123.81** | **151.69** | ***** | 54.83 | 57.03 | n.s |
| BdTR13a | 3 | **94.01** | **99.13** | ***** | **298.81** | **335.52** | ***** | **30.04** | **28.16** | ***** | **20.39** | **7.56** | ***** | **101.63** | **130.20** | ***** | 50.92 | 49.18 | n.s |
| BdTR1i | 4 | **91.62** | **98.50** | ***** | **291.95** | **310.40** | ***** | **27.84** | **25.64** | ***** | **42.46** | **11.32** | ***** | **132.79** | **172.05** | ***** | 67.58 | 65.85 | n.s |
| BdTR2b | 4 | 90.58 | 99.16 | n.s | 286.28 | 334.79 | n.s | 30.04 | 25.59 | n.s | 32.55 | 9.08 | n.s | 135.43 | 184.43 | n.s | 69.31 | 74.33 | n.s |
| BdTR2g | 4 | **88.23** | **99.03** | ***** | **275.58** | **321.83** | ***** | **30.19** | **25.78** | ***** | **54.82** | **10.80** | ***** | 141.09 | 152.06 | n.s | 78.53 | 55.90 | n.s |
| BdTR3c | 4 | **91.01** | **99.87** | ***** | **287.62** | **335.43** | ***** | **29.31** | **24.88** | ***** | **29.50** | **8.46** | ***** | 106.26 | 174.36 | n.s | 58.46 | 55.66 | n.s |
| BdTR5i | 3 | 92.84 | 98.68 | n.s | 292.29 | 324.81 | n.s | 31.54 | 26.77 | n.s | 42.44 | 9.39 | n.s | 97.23 | 141.26 | n.s | 62.53 | 64.18 | n.s |
| BdTR9k | 4 | **95.21** | **98.97** | ***** | 303.20 | 315.82 | n.s | 27.63 | 28.14 | n.s | **19.61** | **14.46** | ***** | 133.27 | 151.78 | n.s | 58.40 | 51.30 | n.s |
| Bis1 | 3 | 93.77 | 98.41 | n.s | 300.90 | 329.00 | n.s | 29.34 | 25.63 | n.s | 21.24 | 7.71 | n.s | 97.16 | 113.62 | n.s | 45.96 | 41.93 | n.s |
| Kah1 | 4 | **94.02** | **99.23** | ***** | 315.47 | 334.60 | n.s | 29.18 | 28.18 | n.s | **31.07** | **9.33** | ***** | **109.04** | **136.56** | ***** | **55.51** | **48.92** | ***** |
| Kah5 | 4 | **93.11** | **98.86** | ***** | 311.66 | 343.46 | n.s | 27.96 | 26.56 | n.s | **33.53** | **7.36** | ***** | **122.20** | **161.41** | ***** | 48.54 | 50.30 | n.s |
| Koz1 | 2 | 95.95 | 99.58 | n.s | 332.48 | 335.81 | n.s | 26.64 | 26.38 | n.s | 16.15 | 7.35 | n.s | 82.20 | 135.72 | n.s | 56.78 | 46.36 | n.s |
| Koz3 | 4 | **85.40** | **99.62** | ***** | **310.63** | **366.61** | ***** | **30.12** | **23.45** | ***** | **44.27** | **5.49** | ***** | 122.40 | 167.18 | n.s | 62.03 | 56.69 | n.s |
| RON2 | 3 | 94.61 | 99.50 | n.s | 335.71 | 366.35 | n.s | 27.23 | 24.52 | n.s | 30.60 | 7.88 | n.s | 93.14 | 133.44 | n.s | 63.23 | 62.31 | n.s |

| **Ecotype** | **n** | **ttlmass** | | | **rmr** | | | **WUE** | | | **leafc** | | | **leafn** | | | **cn** | | |
| --- | --- | --- | --- | --- | --- | --- | --- | --- | --- | --- | --- | --- | --- | --- | --- | --- | --- | --- | --- |
|  |  | **D** | **W** | **W-test** | **D** | **W** | **W-test** | **D** | **W** | **W-test** | **D** | **W** | **W-test** | **D** | **W** | **W-test** | **D** | **W** | **W-test** |
| ABR2 | 4 | 137.09 | 122.76 | n.s | **38.89** | **33.48** | ***** | **11.32** | **10.27** | ***** | -31.00 | -31.29 | n.s | **397.18** | **394.22** | ***** | **35.17** | **39.21** | ***** |
| ABR3 | 4 | 151.72 | 165.80 | n.s | **37.51** | **31.32** | ***** | 12.01 | 9.45 | n.s | **-30.66** | **-31.50** | ***** | **401.27** | **389.46** | ***** | **33.47** | **41.64** | ***** |
| ABR4 | 4 | 86.11 | 93.11 | n.s | **37.90** | **31.43** | ***** | **11.37** | **9.41** | ***** | -30.98 | -31.69 | n.s | **402.26** | **397.76** | ***** | **35.51** | **42.42** | ***** |
| ABR5 | 3 | 138.14 | 117.42 | n.s | 39.48 | 27.78 | n.s | 10.80 | 9.06 | n.s | -31.21 | -32.05 | n.s | 397.90 | 389.95 | n.s | 37.03 | 43.31 | n.s |
| ABR6 | 4 | 113.44 | 110.98 | n.s | **35.52** | **27.32** | ***** | **11.77** | **9.26** | ***** | **-31.30** | **-31.93** | ***** | **404.47** | **389.55** | ***** | **34.51** | **42.18** | ***** |
| ABR8 | 4 | **145.65** | **213.41** | ***** | **23.44** | **19.36** | ***** | 12.49 | 11.39 | n.s | **-31.33** | **-31.58** | ***** | 393.03 | 383.02 | n.s | 31.77 | 34.08 | n.s |
| Adi10 | 4 | **201.51** | **305.91** | ***** | **33.44** | **26.40** | ***** | **12.53** | **12.19** | ***** | **-30.84** | **-31.58** | ***** | 397.39 | 388.03 | n.s | 31.74 | 32.45 | n.s |
| Adi12 | 4 | 211.91 | 226.64 | n.s | **33.45** | **24.94** | ***** | **12.47** | **10.13** | ***** | **-30.34** | **-31.28** | ***** | **400.60** | **390.93** | ***** | **32.22** | **38.80** | ***** |
| Adi2 | 3 | 158.14 | 211.40 | n.s | 34.89 | 23.65 | n.s | 12.80 | 11.38 | n.s | -30.63 | -31.27 | n.s | 396.23 | 389.55 | n.s | 31.17 | 34.69 | n.s |
| Bd1-1 | 2 | 102.62 | 98.88 | n.s | 32.62 | 28.27 | n.s | 12.27 | 8.92 | n.s | -30.71 | -31.69 | n.s | 396.41 | 386.51 | n.s | 32.63 | 43.38 | n.s |
| Bd18-1 | 2 | 188.09 | 219.08 | n.s | 31.97 | 23.64 | n.s | 12.44 | 11.16 | n.s | -30.57 | -31.84 | n.s | 407.79 | 397.53 | n.s | 32.83 | 36.54 | n.s |
| Bd21ctrl | 4 | **153.58** | **139.84** | ***** | **31.91** | **25.87** | ***** | **12.58** | **9.82** | ***** | **-30.22** | **-31.43** | ***** | **403.73** | **388.69** | ***** | **32.31** | **39.63** | ***** |
| Bd21-3 | 3 | 208.92 | 224.17 | n.s | 35.09 | 25.19 | n.s | 11.57 | 11.10 | n.s | -31.03 | -31.93 | n.s | 396.29 | 389.81 | n.s | 34.28 | 35.17 | n.s |
| Bd2-3 | 4 | **162.12** | **191.49** | ***** | **32.24** | **25.42** | ***** | 12.63 | 10.64 | n.s | **-30.73** | **-31.27** | ***** | **394.31** | **381.89** | ***** | **31.43** | **36.48** | ***** |
| Bd30-1 | 4 | **164.29** | **220.57** | ***** | 34.51 | 32.28 | n.s | **11.14** | **10.05** | ***** | **-31.16** | **-32.11** | ***** | **386.45** | **376.32** | ***** | **34.98** | **37.58** | ***** |
| Bd3-1 | 3 | **211.52** | **268.99** | ***** | **31.01** | **24.91** | ***** | **13.24** | **10.75** | ***** | **-30.66** | **-32.08** | ***** | **394.78** | **387.37** | ***** | **29.94** | **36.25** | ***** |
| BdTR10c | 4 | 177.68 | 258.67 | n.s | 31.05 | 30.12 | n.s | 13.38 | 11.49 | n.s | -30.71 | -31.65 | n.s | 402.76 | 390.60 | n.s | 30.43 | 34.09 | n.s |
| BdTR11g | 4 | **177.28** | **221.90** | ***** | **33.12** | **26.75** | ***** | **12.31** | **11.47** | ***** | **-31.34** | **-32.07** | ***** | 401.69 | 391.18 | n.s | 32.85 | 34.58 | n.s |
| BdTR11i | 4 | **178.64** | **208.72** | ***** | **31.09** | **27.21** | ***** | **12.52** | **10.94** | ***** | **-31.11** | **-31.91** | ***** | 399.66 | 384.59 | n.s | 32.06 | 36.11 | n.s |
| BdTR13a | 3 | **152.54** | **179.38** | ***** | **33.35** | **27.44** | ***** | **11.89** | **10.37** | ***** | -31.26 | -32.08 | n.s | **399.54** | **393.72** | ***** | **33.74** | **38.07** | ***** |
| BdTR1i | 4 | 200.37 | 237.90 | n.s | **33.23** | **26.66** | ***** | **12.99** | **11.00** | ***** | **-30.33** | **-31.11** | ***** | **401.43** | **389.86** | ***** | **31.01** | **35.67** | ***** |
| BdTR2b | 4 | 204.81 | 258.75 | n.s | 33.70 | 28.43 | n.s | 12.36 | 9.81 | n.s | -30.46 | -31.97 | n.s | 402.35 | 396.22 | n.s | 32.69 | 40.70 | n.s |
| BdTR2g | 4 | 219.62 | 207.97 | n.s | **35.40** | **27.05** | ***** | **12.52** | **12.08** | ***** | -30.43 | -31.05 | n.s | 397.63 | 392.77 | n.s | 31.94 | 33.34 | n.s |
| BdTR3c | 4 | **164.73** | **230.02** | ***** | **35.42** | **24.14** | ***** | **13.88** | **11.71** | ***** | **-30.84** | **-31.73** | ***** | **404.11** | **392.73** | ***** | **29.21** | **33.63** | ***** |
| BdTR5i | 3 | 159.77 | 205.44 | n.s | 38.12 | 30.36 | n.s | 12.04 | 10.29 | n.s | -31.08 | -31.94 | n.s | 400.25 | 389.96 | n.s | 33.50 | 38.15 | n.s |
| BdTR9k | 4 | 191.67 | 203.08 | n.s | **31.27** | **25.03** | ***** | **12.35** | **10.09** | ***** | **-30.75** | **-31.74** | ***** | **405.45** | **385.74** | ***** | **33.06** | **38.39** | ***** |
| Bis1 | 3 | 143.08 | 155.54 | n.s | 33.24 | 27.64 | n.s | 12.22 | 10.28 | n.s | -31.34 | -32.29 | n.s | 400.67 | 391.98 | n.s | 32.82 | 38.27 | n.s |
| Kah1 | 4 | 164.55 | 185.48 | n.s | **33.74** | **26.03** | ***** | **12.89** | **9.85** | ***** | **-30.89** | **-31.79** | ***** | **402.98** | **390.96** | ***** | **31.44** | **39.79** | ***** |
| Kah5 | 4 | **170.74** | **211.70** | ***** | **28.57** | **23.54** | ***** | **12.54** | **10.60** | ***** | **-30.50** | **-31.22** | ***** | **403.40** | **393.09** | ***** | **32.24** | **37.22** | ***** |
| Koz1 | 2 | 138.98 | 182.08 | n.s | 40.08 | 25.48 | n.s | 12.58 | 10.89 | n.s | -31.28 | -31.70 | n.s | 404.89 | 397.19 | n.s | 32.45 | 36.64 | n.s |
| Koz3 | 4 | 184.43 | 223.87 | n.s | **33.05** | **25.26** | ***** | **13.20** | **11.25** | ***** | **-30.74** | **-31.80** | ***** | **405.07** | **393.25** | ***** | **30.73** | **35.26** | ***** |
| RON2 | 3 | 157.49 | 195.76 | n.s | 38.91 | 31.37 | n.s | 11.70 | 10.03 | n.s | -30.61 | -31.32 | n.s | 404.26 | 396.07 | n.s | 35.48 | 39.74 | n.s |

**Supplementary Table 11**. Linear model (lm) regression analysis for comparative *Brachypodium distachyon* *Bdhn* gene expressions and drought-induced phenotypic trait changes under total watered (W) and dry (D) conditions. Significant p-values (p≤ 0.05*; 0.01**; 0.001***).

|  | *Bdhn*1a | | | | *Bdhn*2 | | | | *Bdhn*3 | | | | *Bdhn*7 | | | |
| --- | --- | --- | --- | --- | --- | --- | --- | --- | --- | --- | --- | --- | --- | --- | --- | --- |
| traits | median | Std error | F-statistic | p-value | median | Std error | F-statistic | p-value | median | Std error | F-statistic | p-value | median | Std error | F-statistic | p-value |
| leaf_rwc | -34.83 | 196.9 | 51.03 | 1.227E-11 *** | -41.95 | 265.3 | 169.3 | <2E-16*** | -8.7 | 564.8 | 136.8 | <2E-16*** | -4.23 | 145.1 | 129.7 | <2E-16*** |
| leaf_wc | -49.32 | 213.2 | 10.19 | 0.001609** | -70.69 | 321.3 | 43.09 | 3.51E-10*** | -166.9 | 682.3 | 22.33 | 4.02E-06*** | -43.52 | 174.8 | 16.68 | 2.32E-05*** |
| lma | -54.08 | 213.8 | 9.029 | 0.0025955 ** | -96.58 | 321.2 | 43.25 | 3.28E-10*** | -188 | 689.2 | 17.37 | 4.38E05*** | -46.48 | 175.3 | 17.44 | 4.22E-05*** |
| pro | -46.12 | 210.7 | 15.82 | 9.36E-05*** | -93.89 | 308.5 | 66.03 | 2.85E-14*** | -127.6 | 649.7 | 47.99 | 4.37E-11*** | -30.73 | 166.5 | 43.91 | 2.47E-10*** |
| abvgrd | -53.51 | 217.9 | 0.1635 | 0.6864 | -136.1 | 346.7 | 4.911 | 0.0277* | -270.9 | 713.4 | 1.055 | 0.30554 | -71.82 | 181.8 | 0.08986 | 0.765 |
| blwgrd | -40.97 | 208.7 | 20.63 | 9.07E-06*** | -72.33 | 333.1 | 24.22 | 1.65E-06*** | -165.2 | 687 | 18.89 | 2.09E-05*** | -38.5 | 173 | 23.87 | 1.95E-06*** |
| ttlmass | -52.88 | 216.7 | 2.607 | 0.1078 | -175 | 350.4 | 0.1187 | 0.73078 | -293.4 | 714.8 | 0.1826 | 0.6695 | -58.21 | 181.3 | 1.305 | 0.255 |
| rmr | -35.02 | 213.1 | 10.56 | 0.001331** | -77.24 | 313.3 | 56.99 | 1.06E-12*** | -152.7 | 674.7 | 27.97 | 2.90E-07*** | -41.8 | 174 | 20.85 | 8.16E-06*** |
| WUE | -129.55 | 206.2 | 26.68 | 5.26E-07*** | -52.93 | 308 | 66.88 | 2.04E-14*** | -122.5 | 664.9 | 35.53 | 9.50E-09*** | -31.64 | 167.8 | 39.49 | 1.67E-09*** |
| leafC | -39.98 | 210.7 | 15.95 | 8.80E-05*** | -90.2 | 323.7 | 39.07 | 2E-09*** | -183.4 | 682.8 | 21.94 | 4.85E-06*** | -50.17 | 175.8 | 15.87 | 9.14E-05*** |
| leafN | -42.83 | 199.8 | 43.07 | 3.54E-10*** | -62.2 | 311.9 | 59.54 | 3.77E-13*** | -160.7 | 658.5 | 40.66 | 1.E-09*** | -44.7 | 169 | 36.01 | 7.69E-09*** |
| CN | -40.24 | 198.4 | 46.9 | 6.93E-11*** | -58.73 | 307.7 | 67.44 | 1.64E-14*** | -131.6 | 652.1 | 45.95 | 1.04E-10*** | -33.83 | 168 | 39.09 | 1.98E-09*** |

**Supplementary Table 12**. Phylogenetic signal of **(a)** dehydrin gene expressions, **(b)** drought-induced phenotypic traits changes and **(c)** climate niche variation assessed in the *B. distachyon* nuclear species tree (Gordon et al. 2017). Blomberg’s K and Pagel’s lambda values close to one indicate phylogenetic signal and values close to zero phylogenetic independence. W, watered conditions; D, drought conditions. K, p-values based on 1000 randomizations; lambda, p-values based on the Likelihood Ratio test. Significant test values are indicated in bold.

**(a)**

| ***Bdhn* gene** | **K** | **P-value** | **lambda(ʎ)** | **logL(ʎ)** | **LR(ʎ=0)** | **P-value** |
| --- | --- | --- | --- | --- | --- | --- |
| *Bdhn*1aW | 0.00207548 | 0.579 | 6.6113E-05 | -170094 | -9.01E-04 | 1 |
| *Bdhn*2W | 0.00467069 | 0.235 | 6.6113E-05 | -111372 | -9.77E-04 | 1 |
| *Bdhn*3W | 0.00236035 | 0.553 | 6.6113E-05 | -90.2705 | -1.13E-03 | 1 |
| *Bdhn*7W | 0.00162038 | 0.695 | 6.6113E-05 | -59.9152 | -1.39E-03 | 1 |
| *Bdhn*1aD | 0.0038235 | 0.317 | 6.6113E-05 | -194.936 | -1.03E-03 | 1 |
| *Bdhn*2D | 0.00311909 | 0.364 | 6.6113E-05 | -209.636 | -1.03E-03 | 1 |
| *Bdhn*3D | 0.00257041 | 0.71 | 6.6113E-05 | -237.428 | -1.06E-03 | 1 |
| *Bdhn*7D | 0.00137242 | 0.753 | 6.6113E-05 | -196.53 | -1.04E-03 | 1 |

**(b)**

|  | Drought | | | | | | Watered | | | | | |
| --- | --- | --- | --- | --- | --- | --- | --- | --- | --- | --- | --- | --- |
| **Phenotypic trait** | **K** | **P-value (based on 1000 randomizations)** | **lambda(ʎ)** | **logL(ʎ)** | **LR(ʎ=0)** | **P-value** | **K** | **P-value** | **lambda(ʎ)** | **logL(ʎ)** | **LR(ʎ=0)** | **P-value** |
| leaf_rwc | 0.00595303 | 0.202 | 0.8240433 | -70.83287 | -71.94764 | 0.1353942 | 0.00633355 | 0.137 | 6.6113E-05 | -21.1366 | -21.1361 | 1 |
| leaf_wc | 0.0191719 | 0.029 | 0.5318181 | -129.0792 | -129.2025 | 0.6194317 | 0.00601205 | 0.142 | 0.712778 | -124.3118 | -124.641 | 0.4171331 |
| lma | 0.00655413 | 0.198 | 0.4014225 | -60.3207 | -60.04825 | 1 | **0.0127346** | **0.067** | 0.5672695 | -53.1973 | -54.02652 | 0.1978138 |
| pro | 0.00178069 | 0.645 | 6.61128E-05 | -118.2053 | -118.2048 | 1 | 0.00751385 | 0.279 | 6.6113E-05 | -65.7397 | -65.73915 | 1 |
| **abvrgd** | **0.0548587** | **0.005** | **0.9862024** | **-132.3802** | **-137.5324** | **0.00132712** | 0.00982265 | 0.118 | **0.9056281** | **-143.4905** | **-147.2048** | **0.00641901** |
| blwgrd | 0.00799846 | 0.135 | 0.6778591 | -112.6454 | -113.232 | 0.2787568 | 0.00503958 | 0.192 | 0.372553 | -117.8161 | -117.602 | 1 |
| **ttlmass** | **0.0254584** | **0.02** | **0.9586421** | **-144.0437** | **-147.1176** | **0.01315614** | 0.00760606 | 0.15 | **0.8313881** | **-153.3088** | **-155.4348** | **0.0392068** |
| rmr | 0.0117638 | 0.1 | **0.8587116** | **-71.60054** | **-74.00027** | **0.02846858** | **0.0198041** | **0.031** | **0.8500612** | **-65.57032** | **-69.23347** | **0.00679529** |
| **delta13c** | **0.0206935** | **0.023** | 0.5455282 | -9.438273 | -9.102205 | 1 | 0.00079528 | 0.921 | 6.6113E-05 | -9.565582 | -9.565045 | 1 |
| leafc | 0.00564166 | 0.206 | 6.61128E-05 | -86.81046 | -86.80988 | 1 | 0.00344133 | 0.413 | 6.6113E-05 | -84.79111 | -84.79067 | 1 |
| **leafn** | **0.0127895** | **0.066** | **0.8379002** | **-52.22653** | **-57.04403** | **0.00190906** | 0.00111356 | 0.809 | 0.6412875 | -71.1111 | -72.40537 | 0.1076397 |
| **cn** | **0.015233** | **0.051** | **0.8350859** | **-24.49639** | **-28.06983** | **0.00750945** | 0.00101427 | 0.842 | 0.6145648 | -35.11085 | -36.01121 | 0.1796235 |

**(c)**

| **Trait** | **K** | **p-value** | **lambda(ʎ)** | **logL(ʎ)** | **LR(ʎ=0)** | **p-value** |
| --- | --- | --- | --- | --- | --- | --- |
| **PCA1** | **0.0201727** | **0.024** | **0.905784** | **-70.641** | **12.567** | **0.000392627** |
