## Supplementary figures for "Evolution and functional dynamics of dehydrins in model *Brachypodium* grasses"

### Slide 1
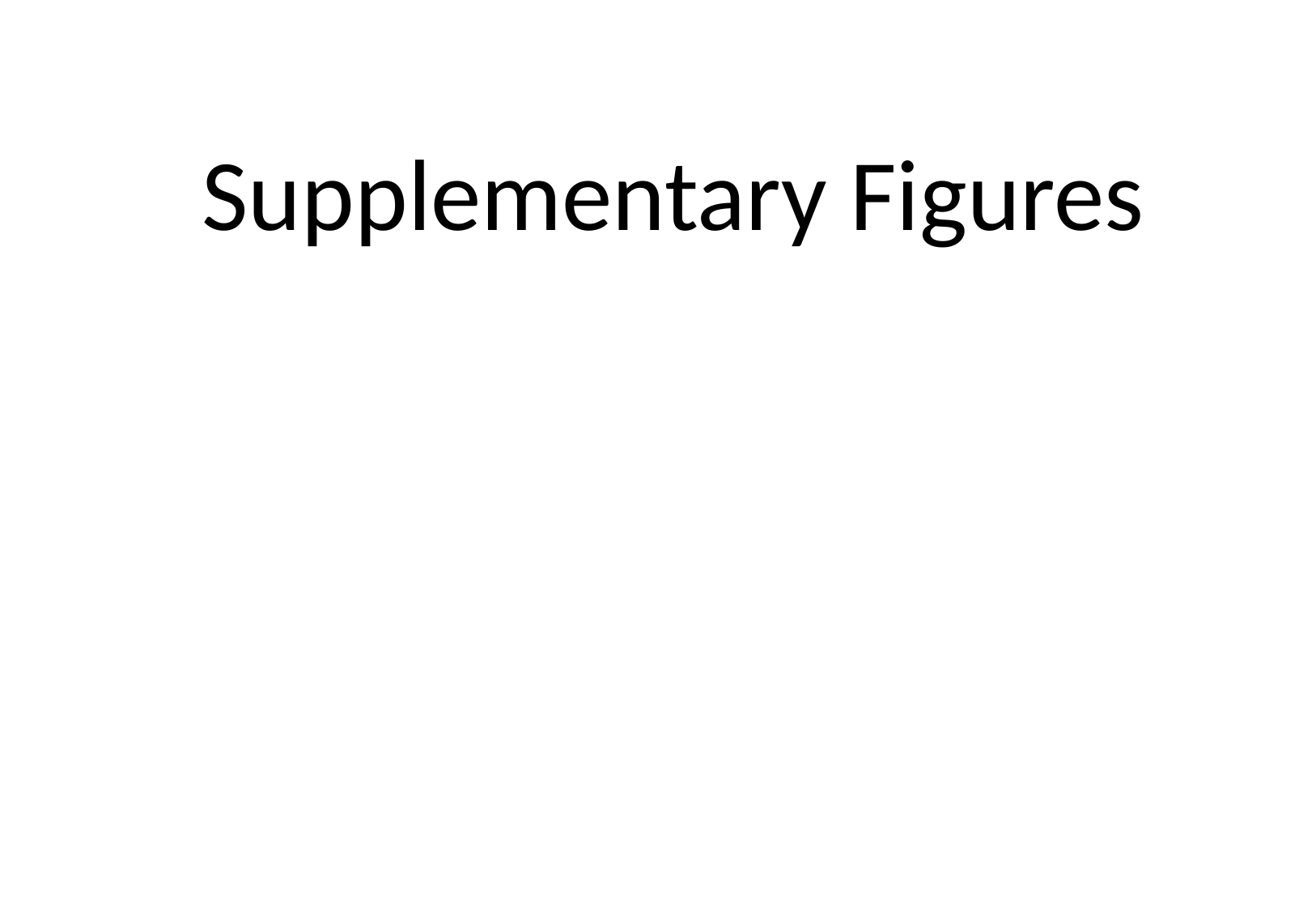

Supplementary Figures

### Slide 2
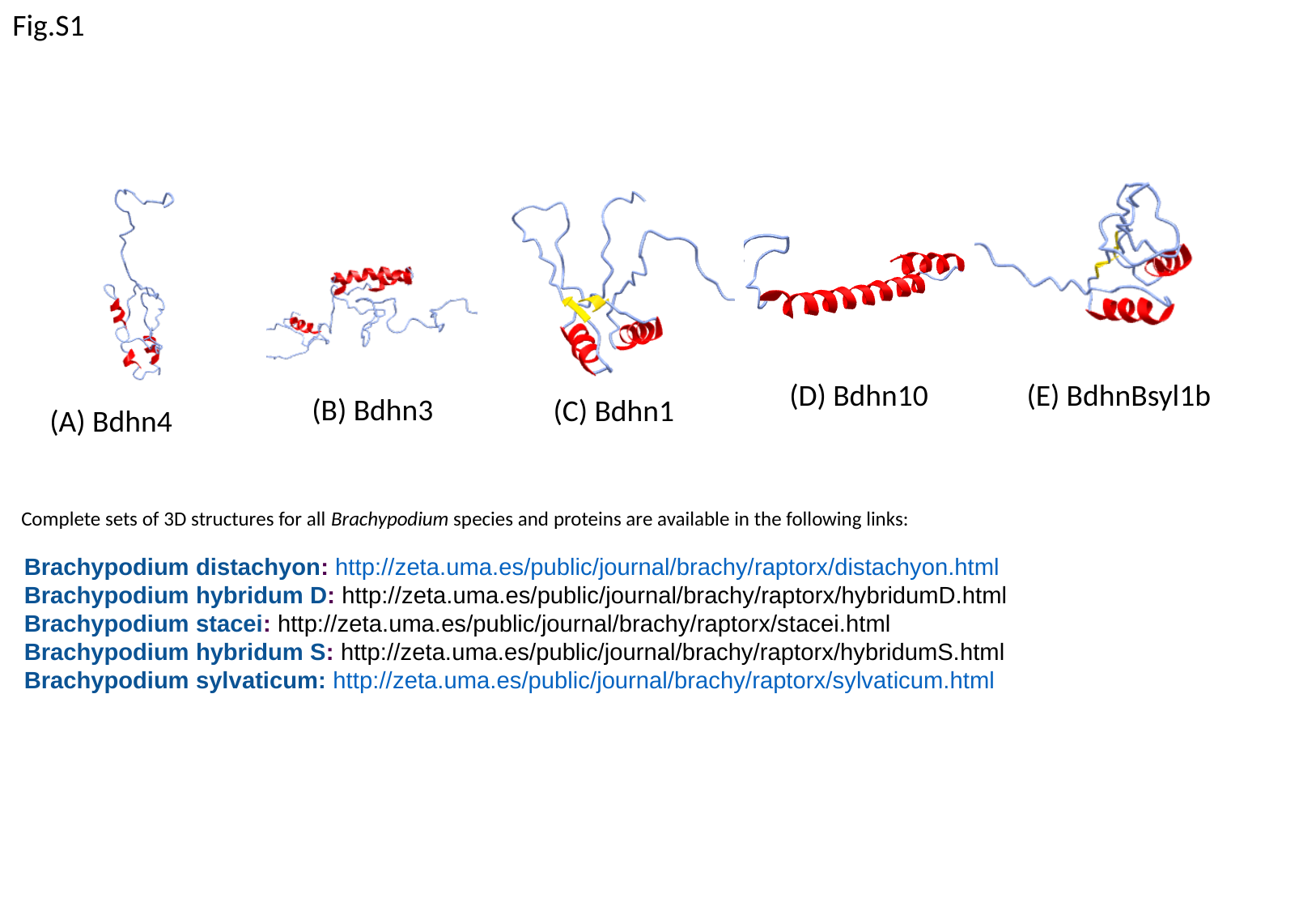

Fig.S1
(E) BdhnBsyl1b
(B) Bdhn3
(C) Bdhn1
(D) Bdhn10
(A) Bdhn4
Complete sets of 3D structures for all Brachypodium species and proteins are available in the following links:
Brachypodium distachyon: http://zeta.uma.es/public/journal/brachy/raptorx/distachyon.html
Brachypodium hybridum D: http://zeta.uma.es/public/journal/brachy/raptorx/hybridumD.html
Brachypodium stacei: http://zeta.uma.es/public/journal/brachy/raptorx/stacei.html
Brachypodium hybridum S: http://zeta.uma.es/public/journal/brachy/raptorx/hybridumS.html
Brachypodium sylvaticum: http://zeta.uma.es/public/journal/brachy/raptorx/sylvaticum.html

### Slide 3
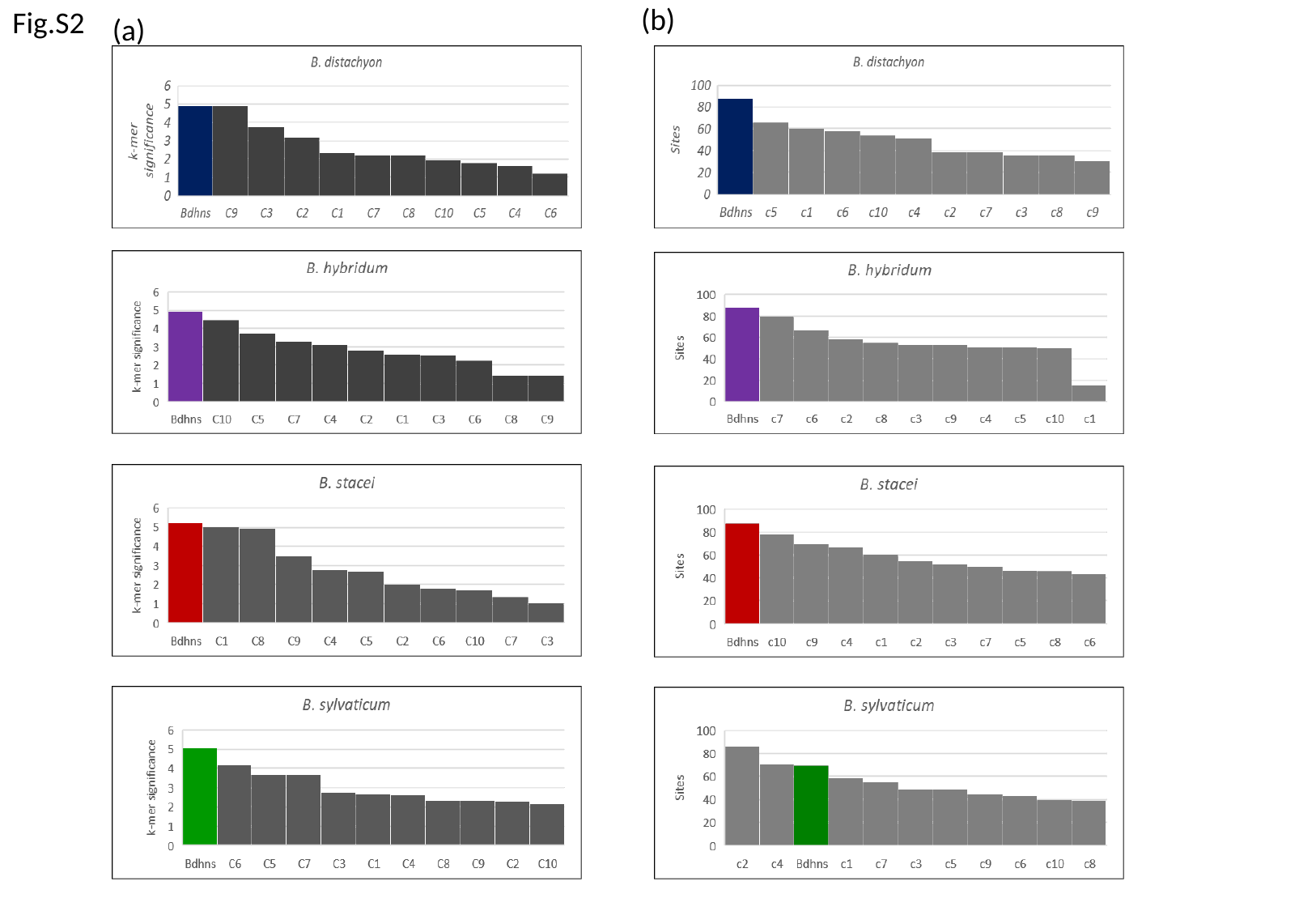

(b)
Fig.S2
(a)

### Slide 4
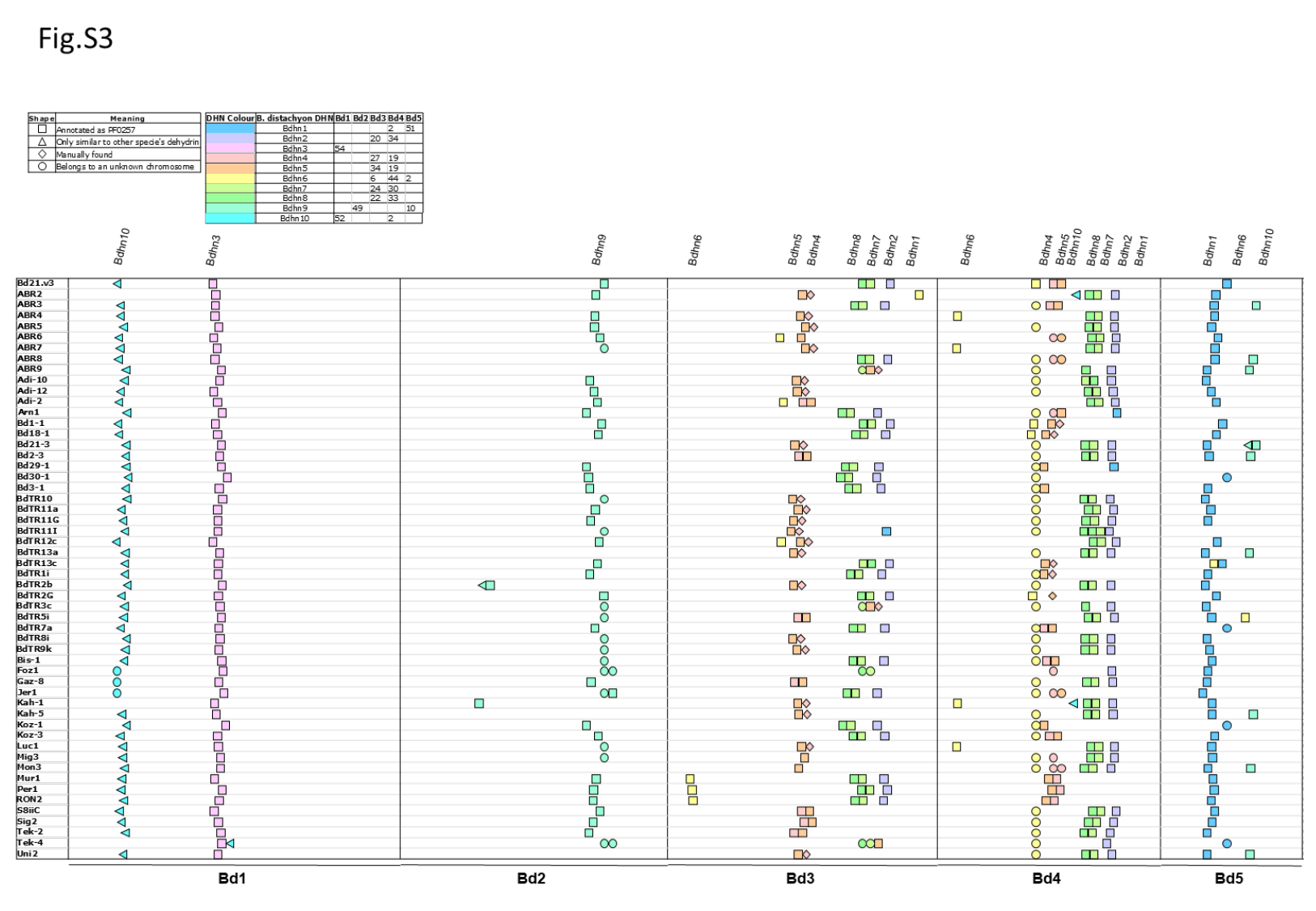

### Slide 5
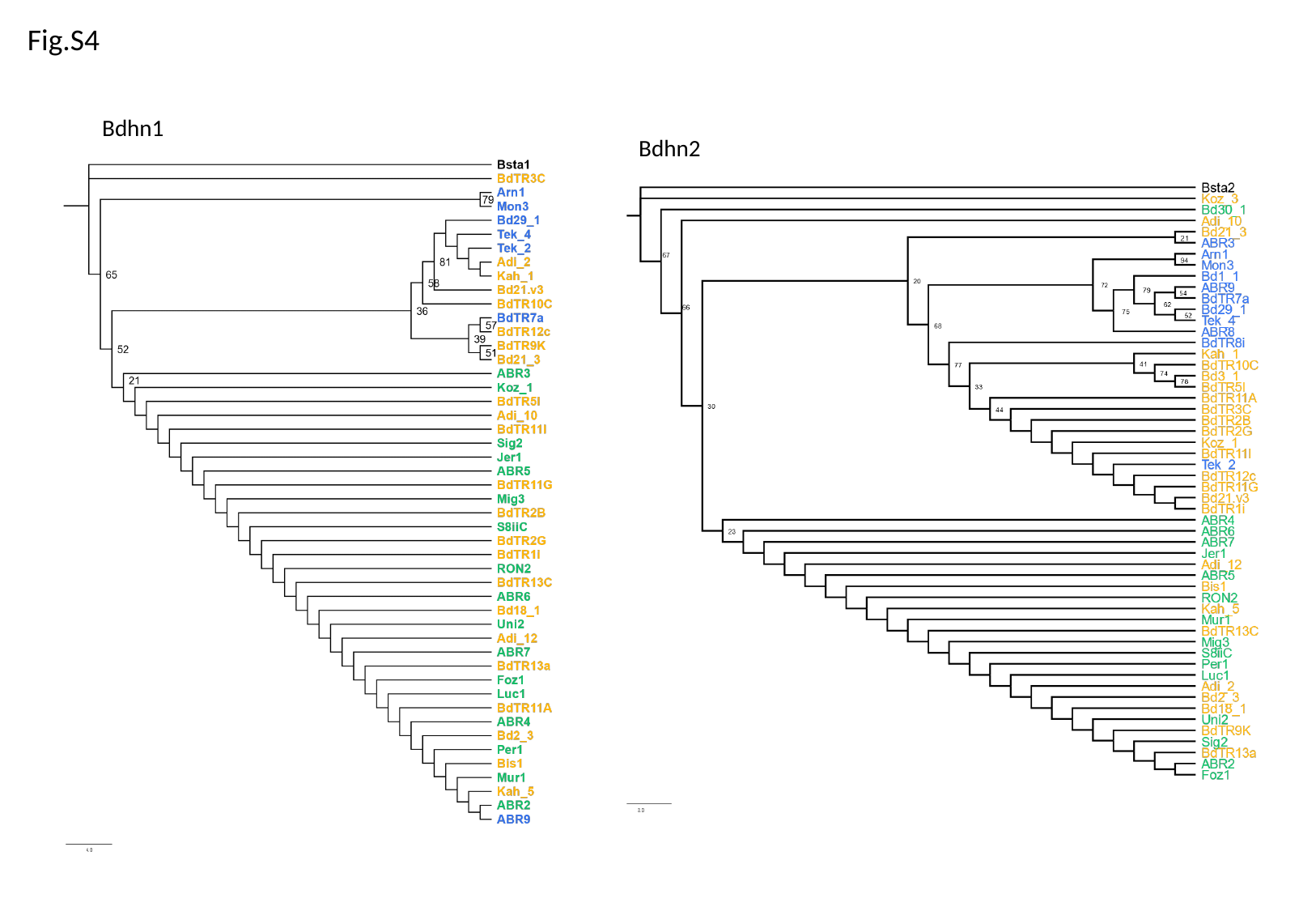

Fig.S4
Bdhn1
Bdhn2

### Slide 6
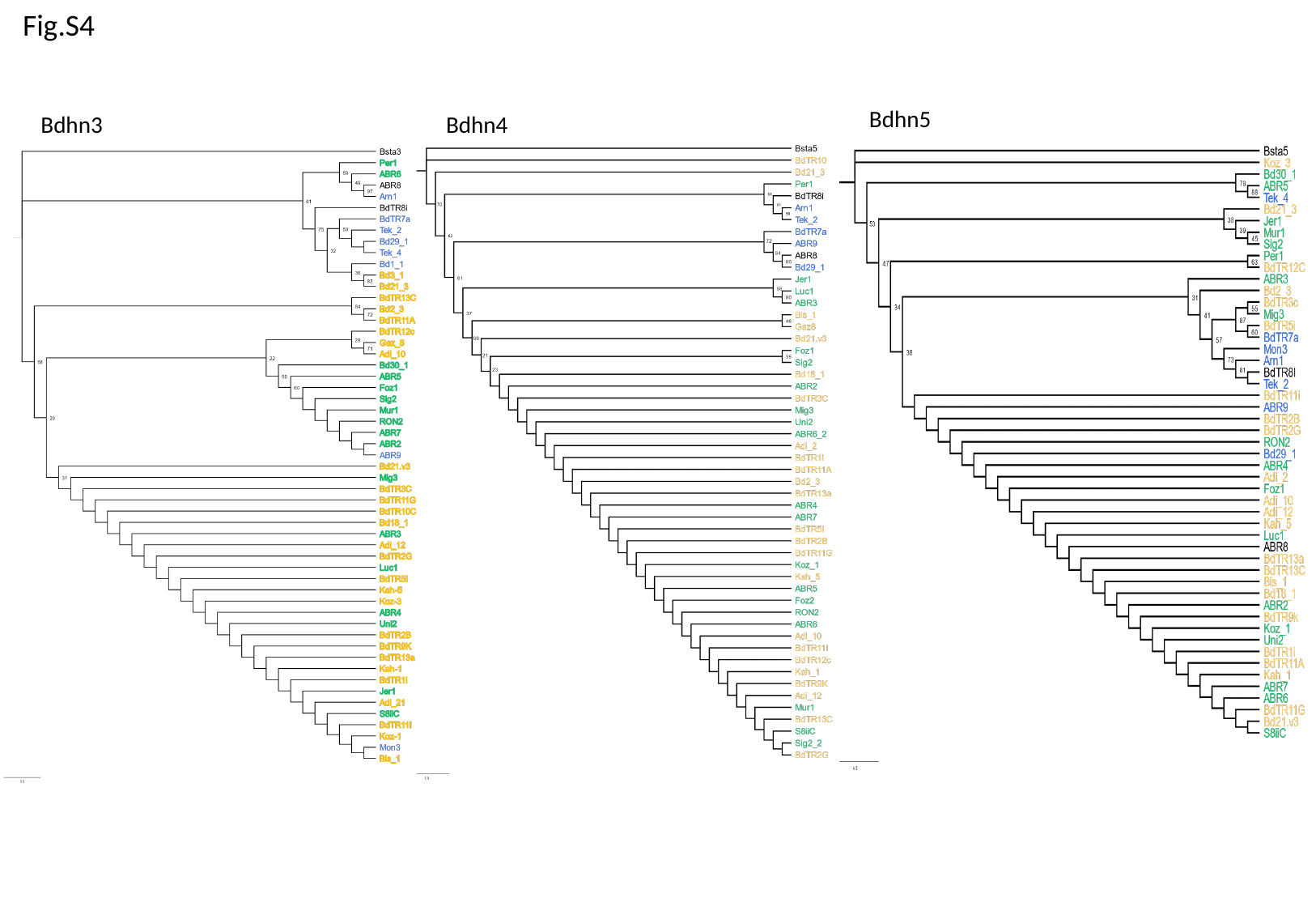

Fig.S4
Bdhn5
Bdhn4
Bdhn3

### Slide 7
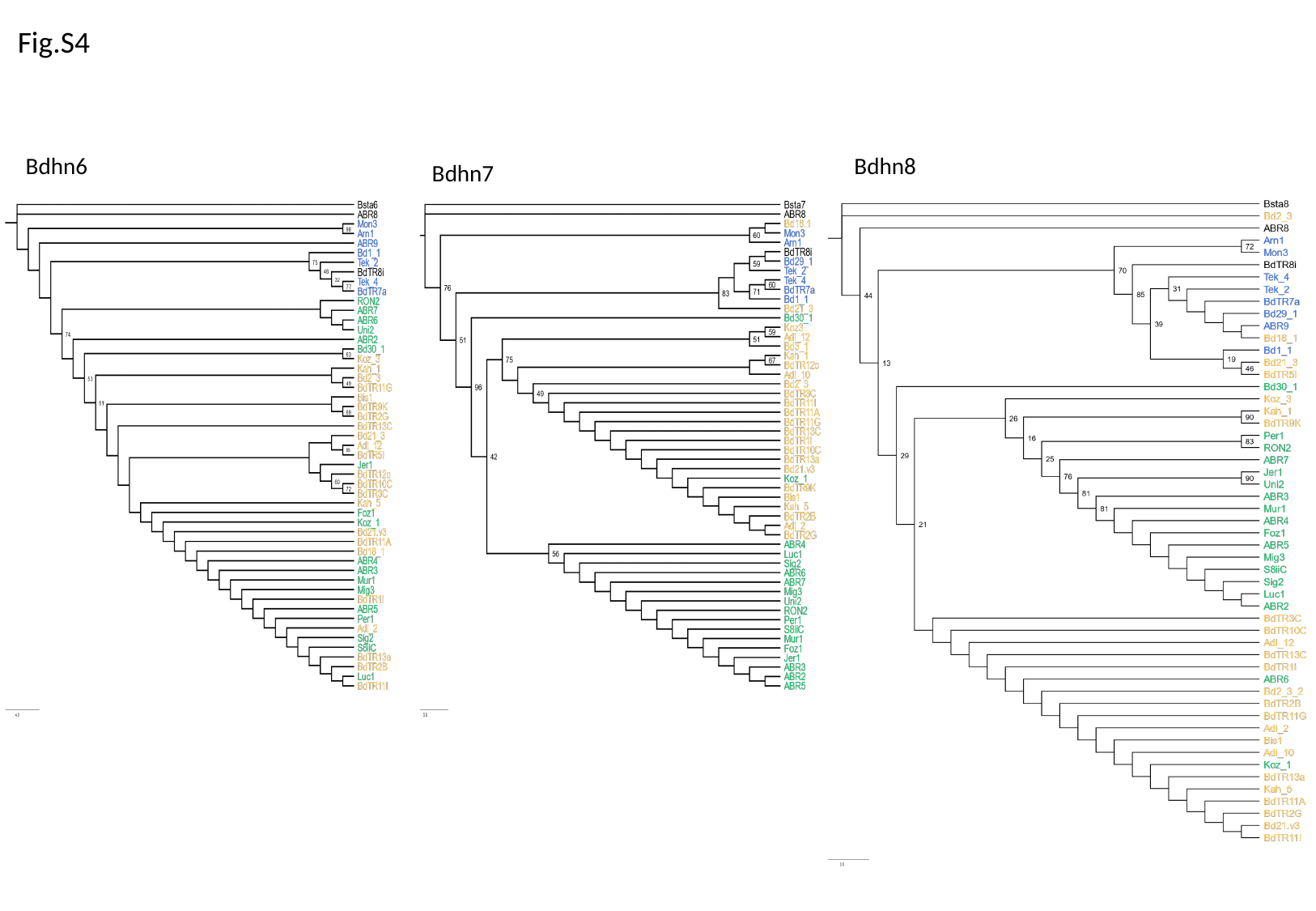

Fig.S4
Bdhn6
Bdhn8
Bdhn7

### Slide 8
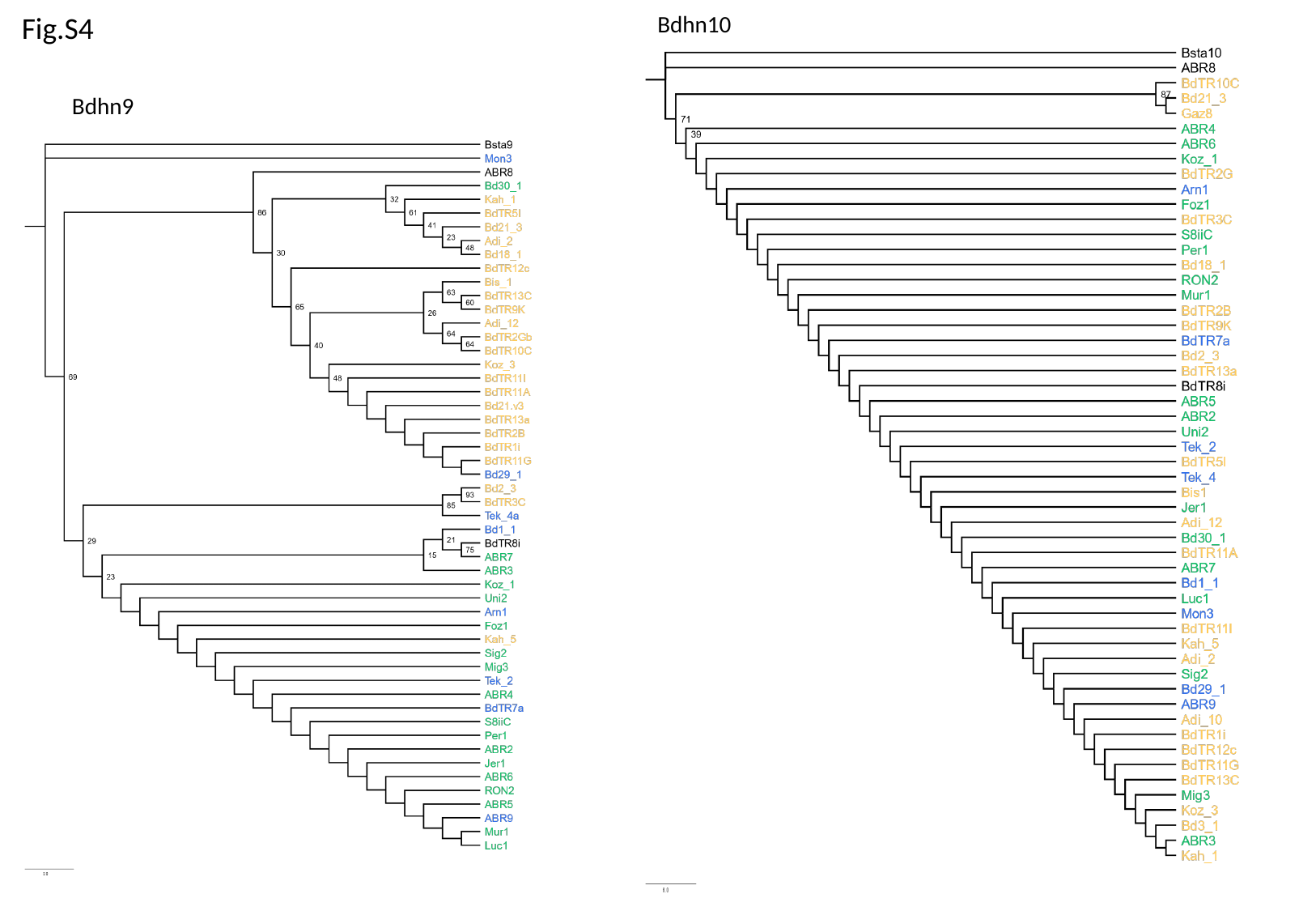

Fig.S4
Bdhn10
Bdhn9

### Slide 9
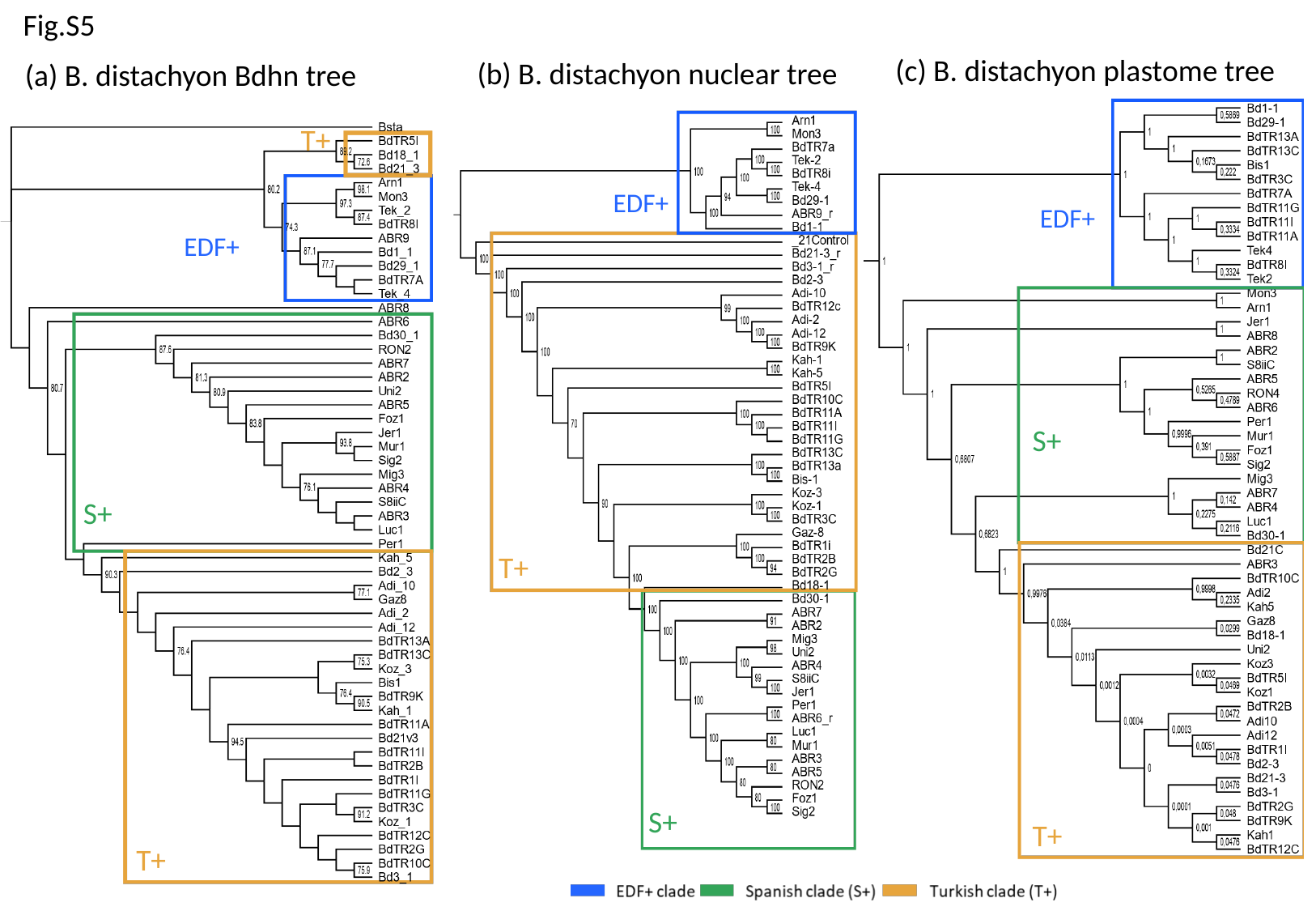

Fig.S5
(c) B. distachyon plastome tree
EDF+
S+
T+
(b) B. distachyon nuclear tree
EDF+
(a) B. distachyon Bdhn tree
T+
EDF+
S+
T+
T+
S+

### Slide 10
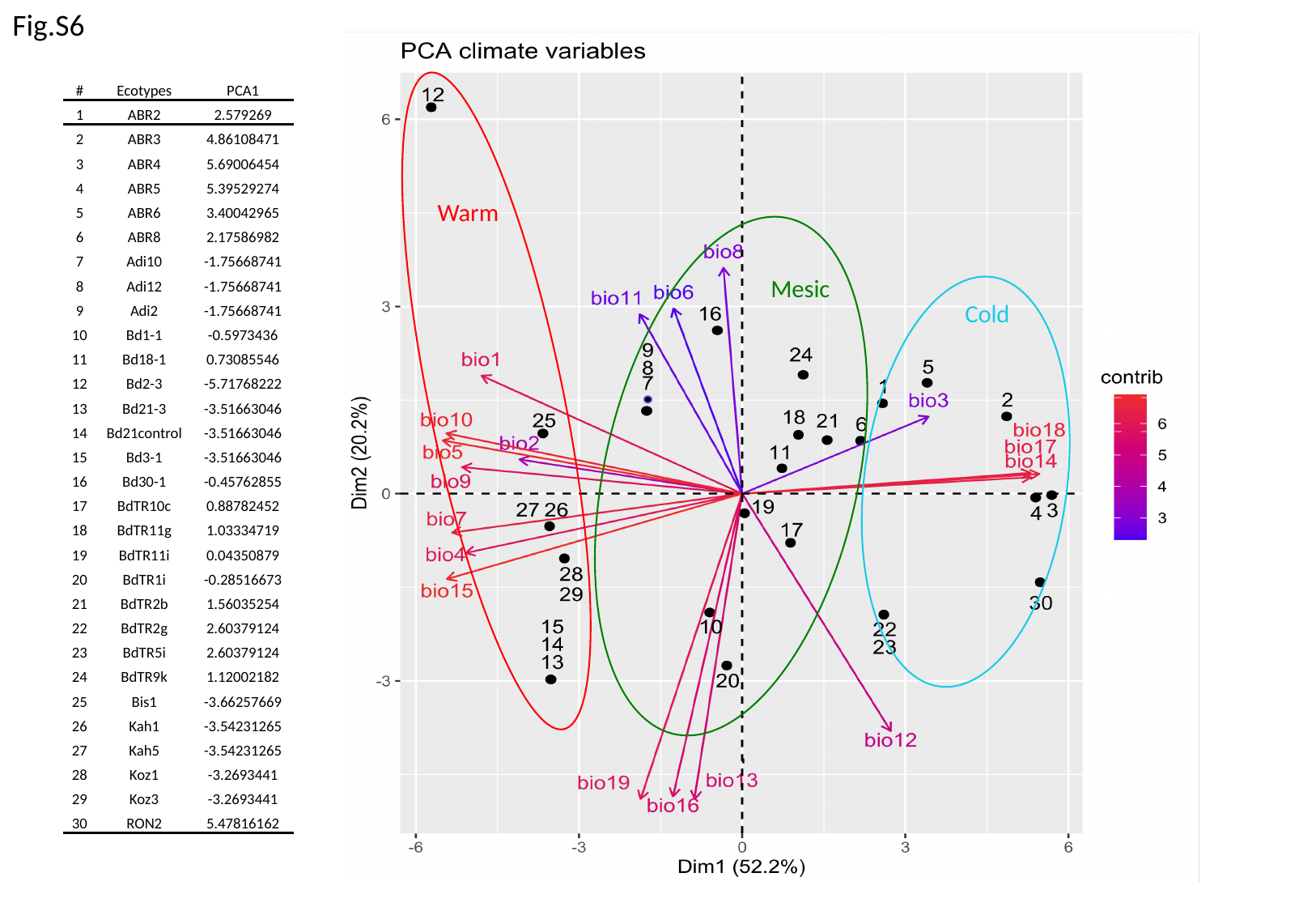

Fig.S6
Warm
Mesic
Cold
| # | Ecotypes | PCA1 |
| --- | --- | --- |
| 1 | ABR2 | 2.579269 |
| 2 | ABR3 | 4.86108471 |
| 3 | ABR4 | 5.69006454 |
| 4 | ABR5 | 5.39529274 |
| 5 | ABR6 | 3.40042965 |
| 6 | ABR8 | 2.17586982 |
| 7 | Adi10 | -1.75668741 |
| 8 | Adi12 | -1.75668741 |
| 9 | Adi2 | -1.75668741 |
| 10 | Bd1-1 | -0.5973436 |
| 11 | Bd18-1 | 0.73085546 |
| 12 | Bd2-3 | -5.71768222 |
| 13 | Bd21-3 | -3.51663046 |
| 14 | Bd21control | -3.51663046 |
| 15 | Bd3-1 | -3.51663046 |
| 16 | Bd30-1 | -0.45762855 |
| 17 | BdTR10c | 0.88782452 |
| 18 | BdTR11g | 1.03334719 |
| 19 | BdTR11i | 0.04350879 |
| 20 | BdTR1i | -0.28516673 |
| 21 | BdTR2b | 1.56035254 |
| 22 | BdTR2g | 2.60379124 |
| 23 | BdTR5i | 2.60379124 |
| 24 | BdTR9k | 1.12002182 |
| 25 | Bis1 | -3.66257669 |
| 26 | Kah1 | -3.54231265 |
| 27 | Kah5 | -3.54231265 |
| 28 | Koz1 | -3.2693441 |
| 29 | Koz3 | -3.2693441 |
| 30 | RON2 | 5.47816162 |

### Slide 11
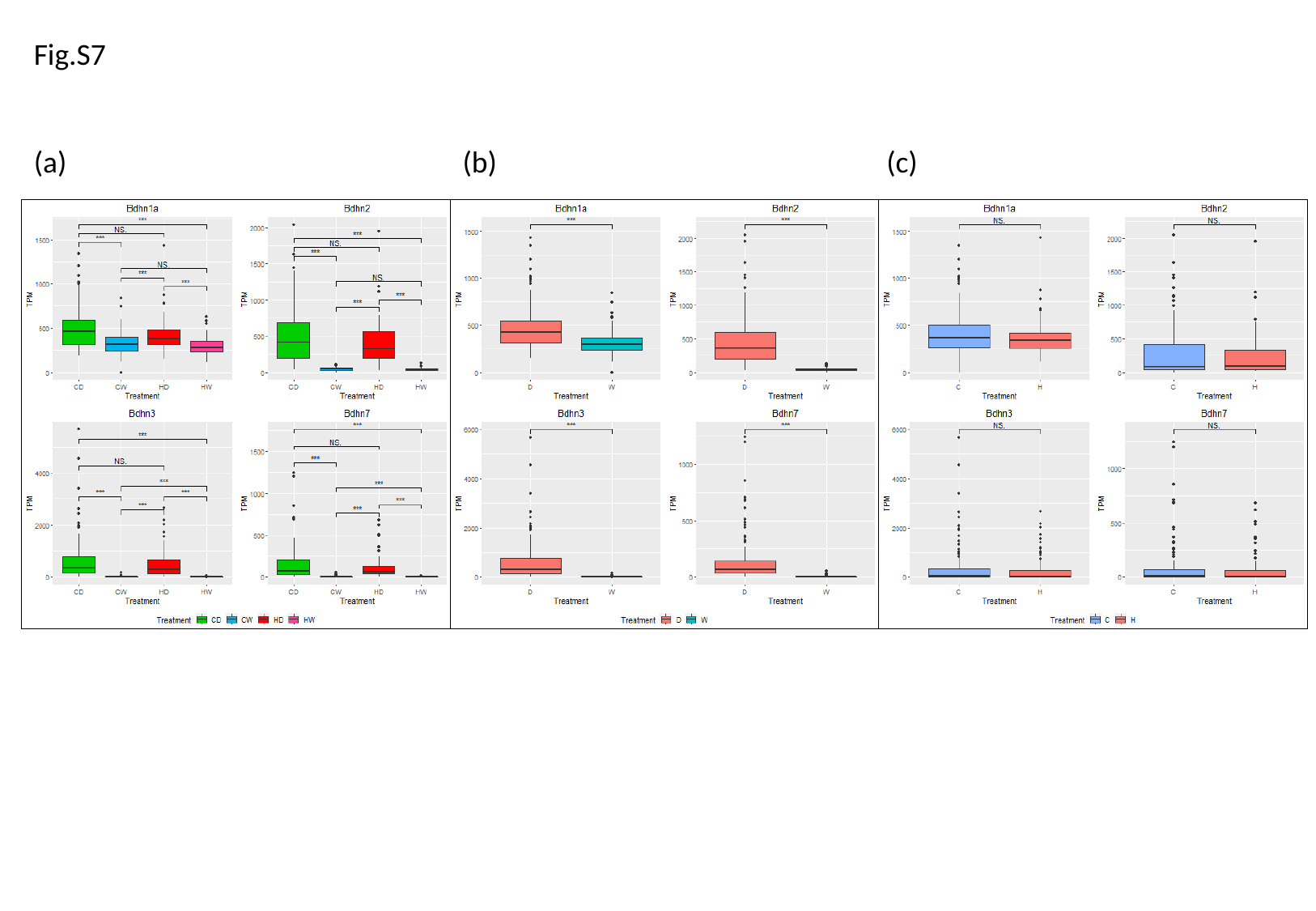

Fig.S7
(a)
(b)
(c)

### Slide 12
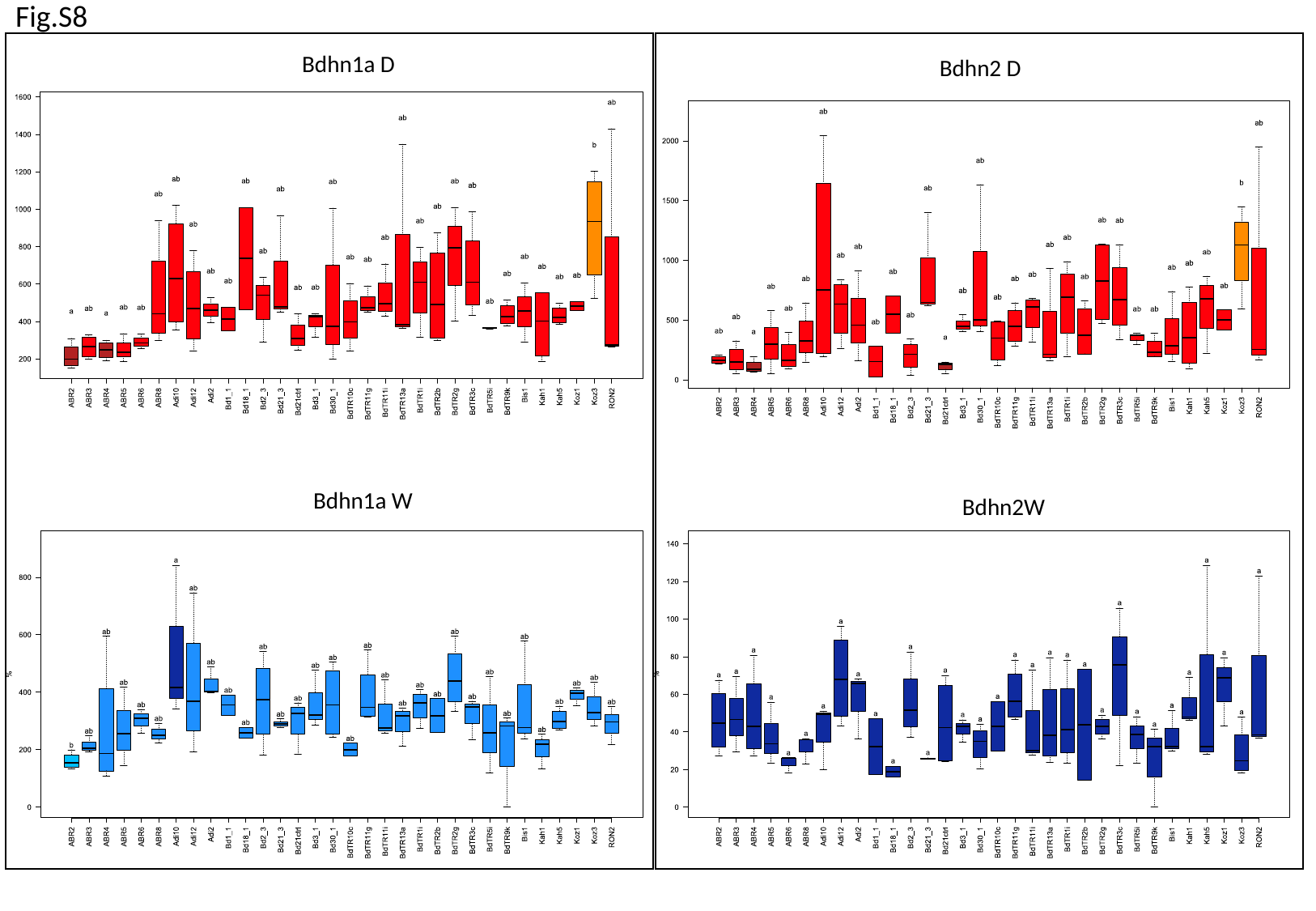

Fig.S8
Bdhn1a D
Bdhn2 D
Bdhn1a W
Bdhn1a W
Bdhn2W

### Slide 13
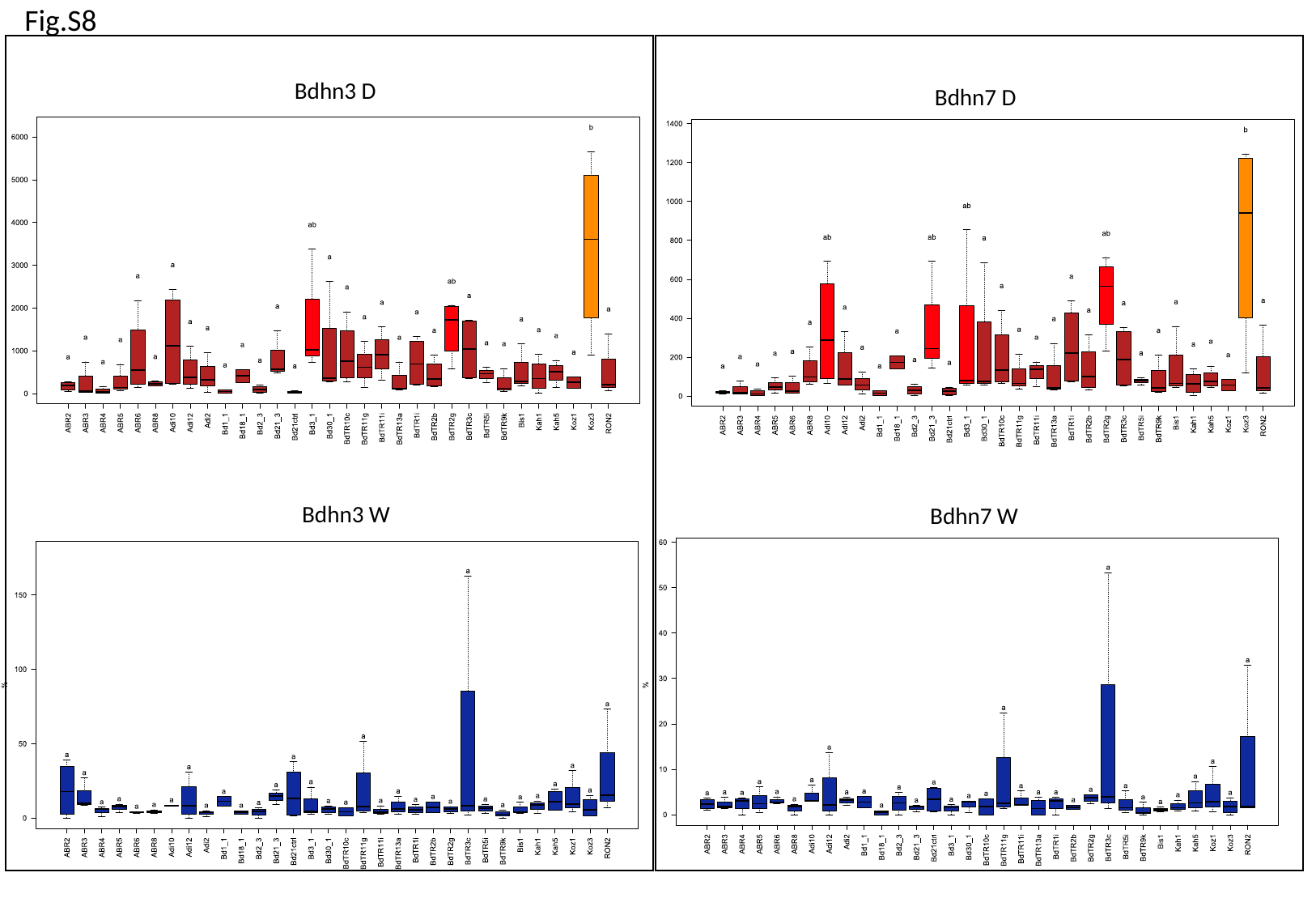

Fig.S8
Bdhn3 D
Bdhn7 D
Bdhn3 W
Bdhn7 W

### Slide 14
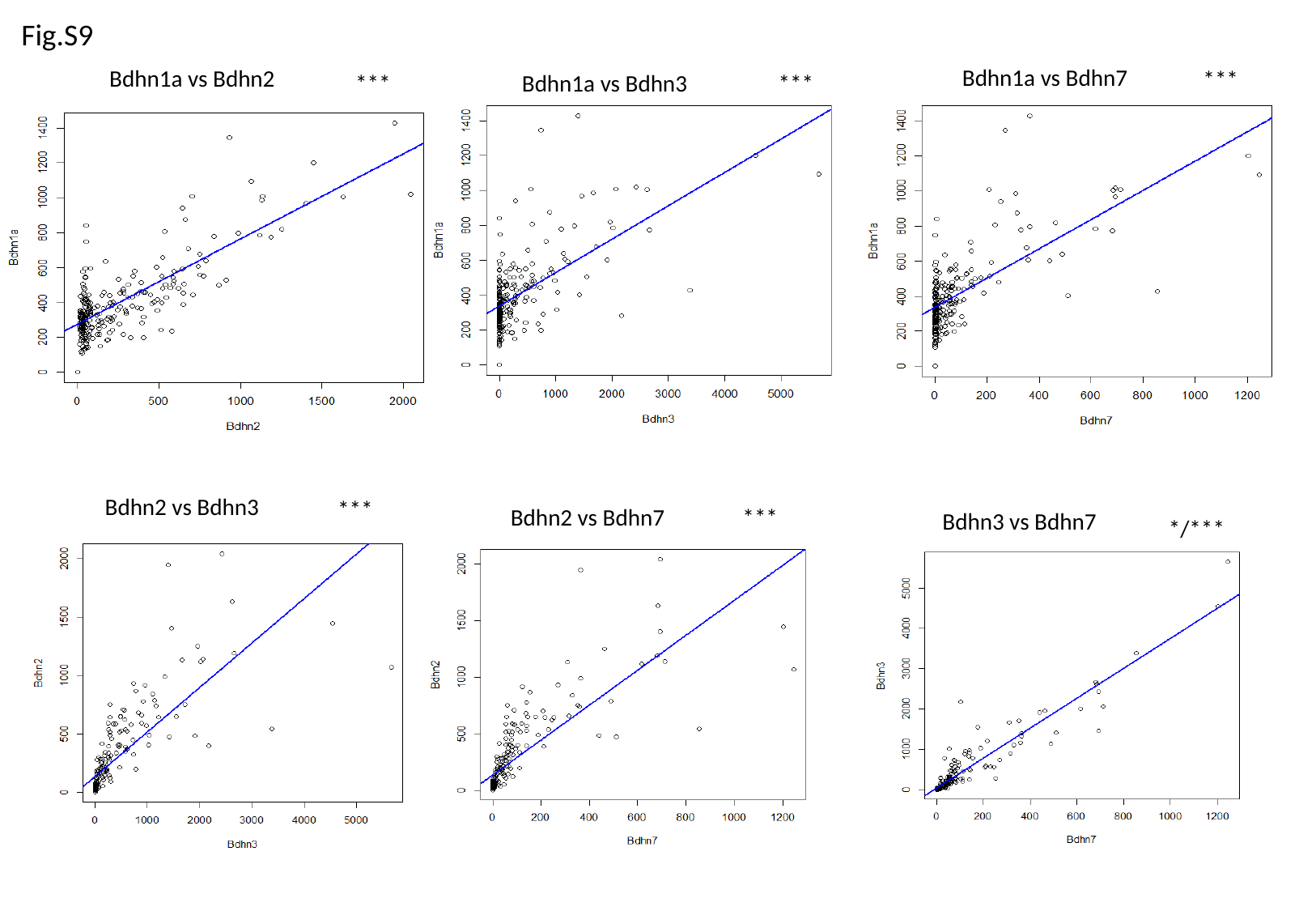

Fig.S9
Bdhn1a vs Bdhn3
***
Bdhn1a vs Bdhn7
***
Bdhn1a vs Bdhn2
***
Bdhn2 vs Bdhn3
***
***
Bdhn2 vs Bdhn7
Bdhn3 vs Bdhn7
*/***

### Slide 15
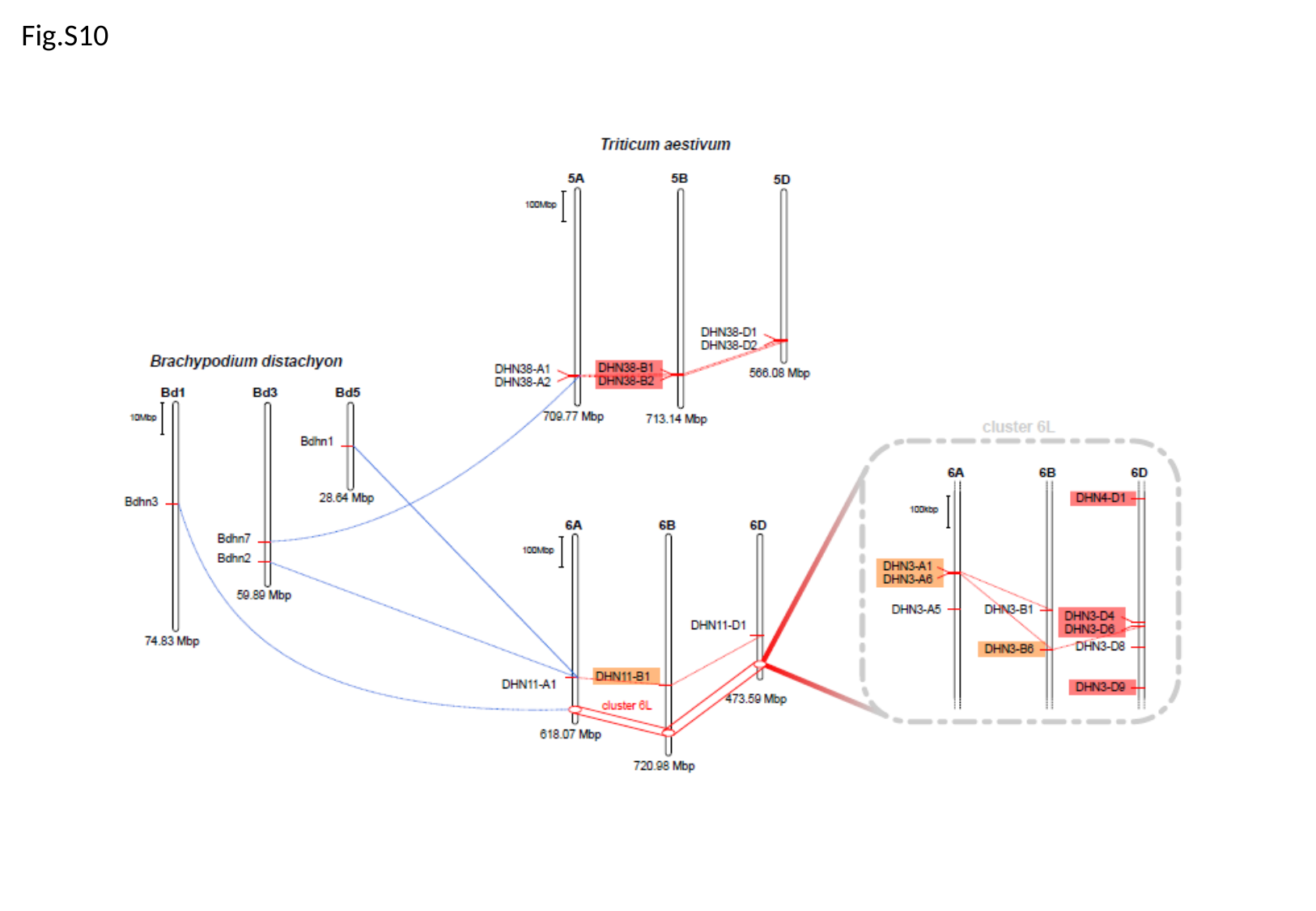

Fig.S10

### Slide 16
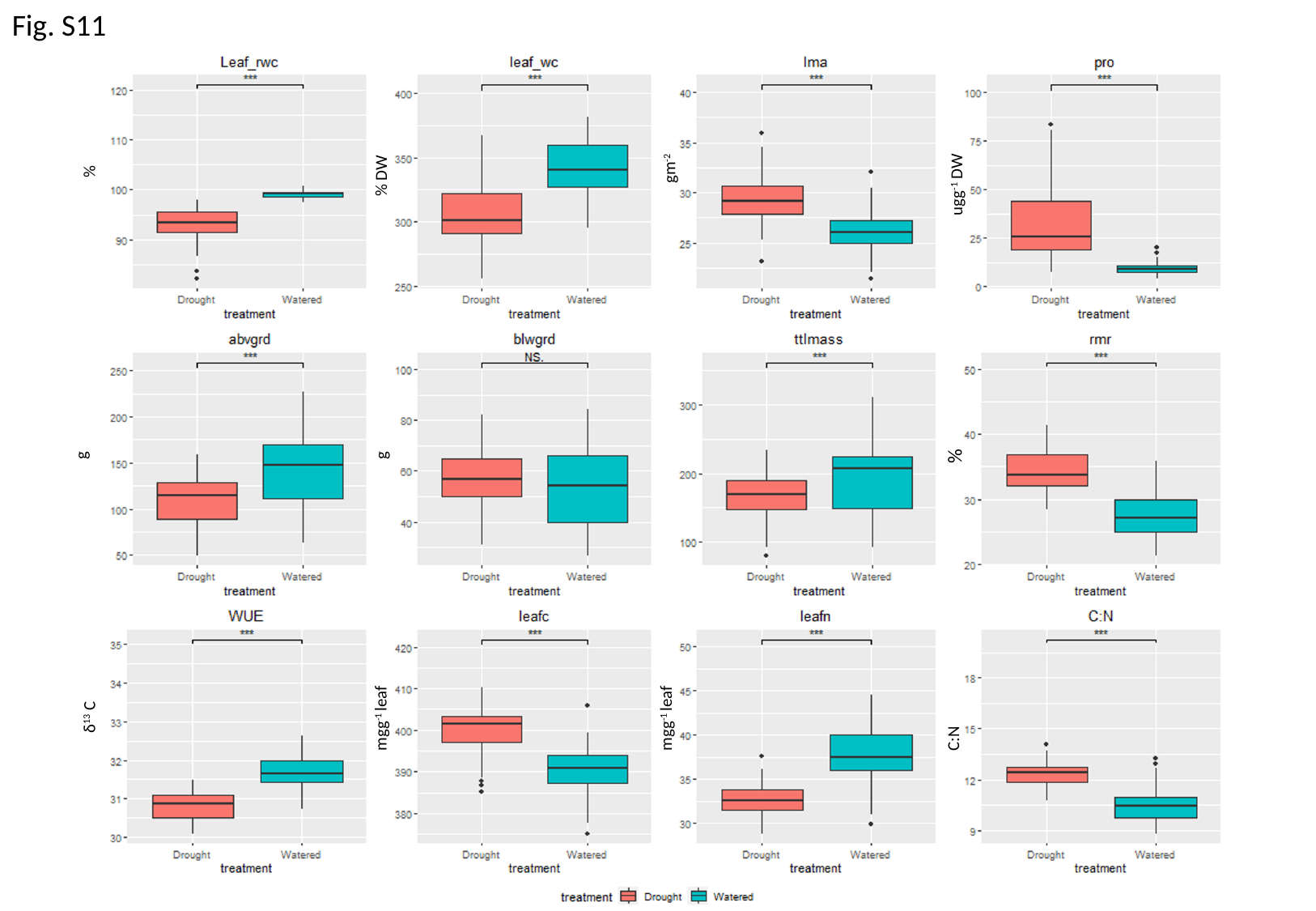

Fig. S11
gm^-2^
%
% DW
ugg^-1^ DW
%
g
g
δ^13^ C
mgg^-1^ leaf
mgg^-1^ leaf
C:N

### Slide 17
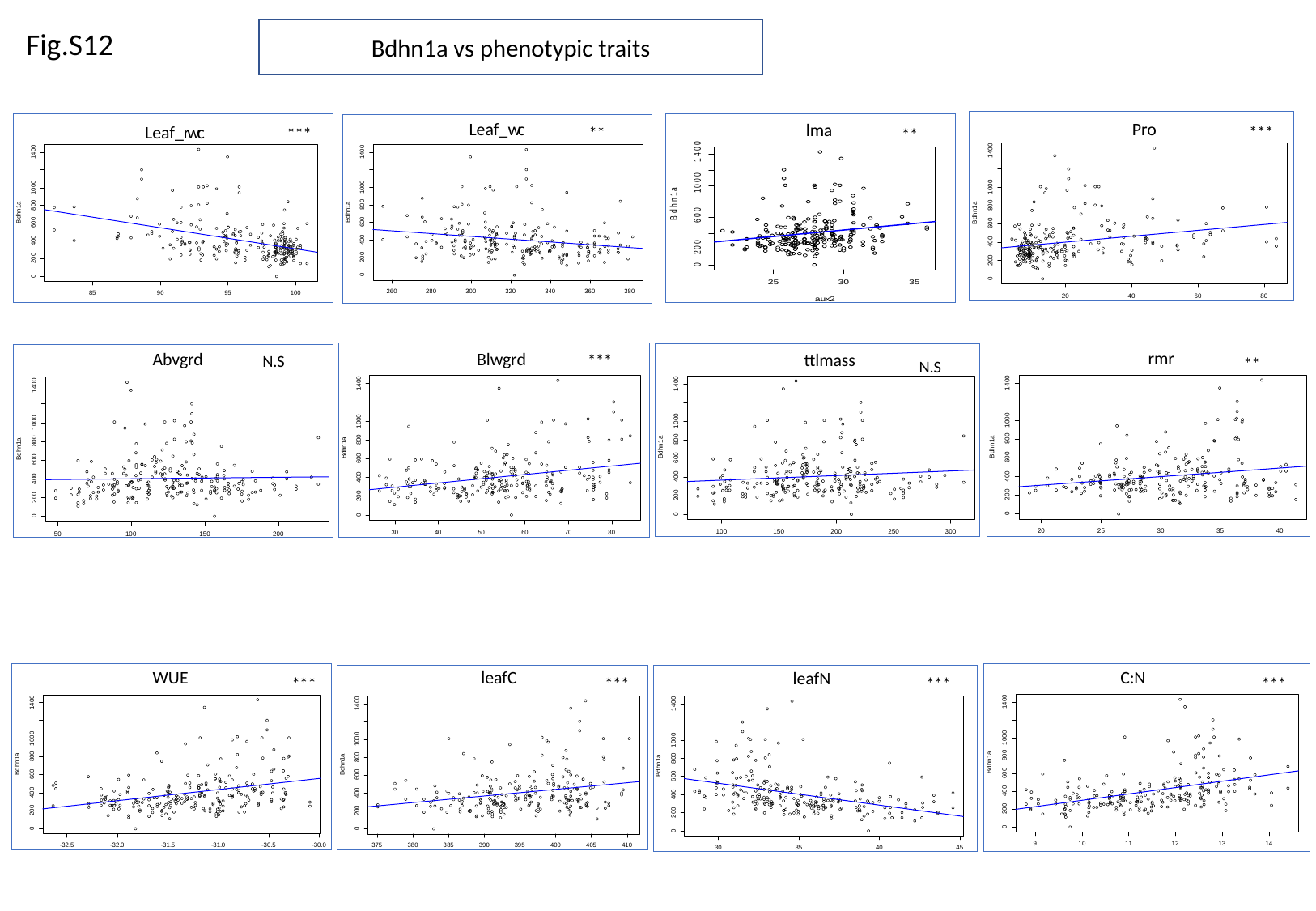

Fig.S12
Bdhn1a vs phenotypic traits
***
Pro
**
lma
***
**
Leaf_wc
Pro
Leaf_rwc
***
N.S
rmr
**
Blwgrd
Abvgrd
ttlmass
N.S
WUE
leafC
C:N
***
***
***
***
leafN

### Slide 18
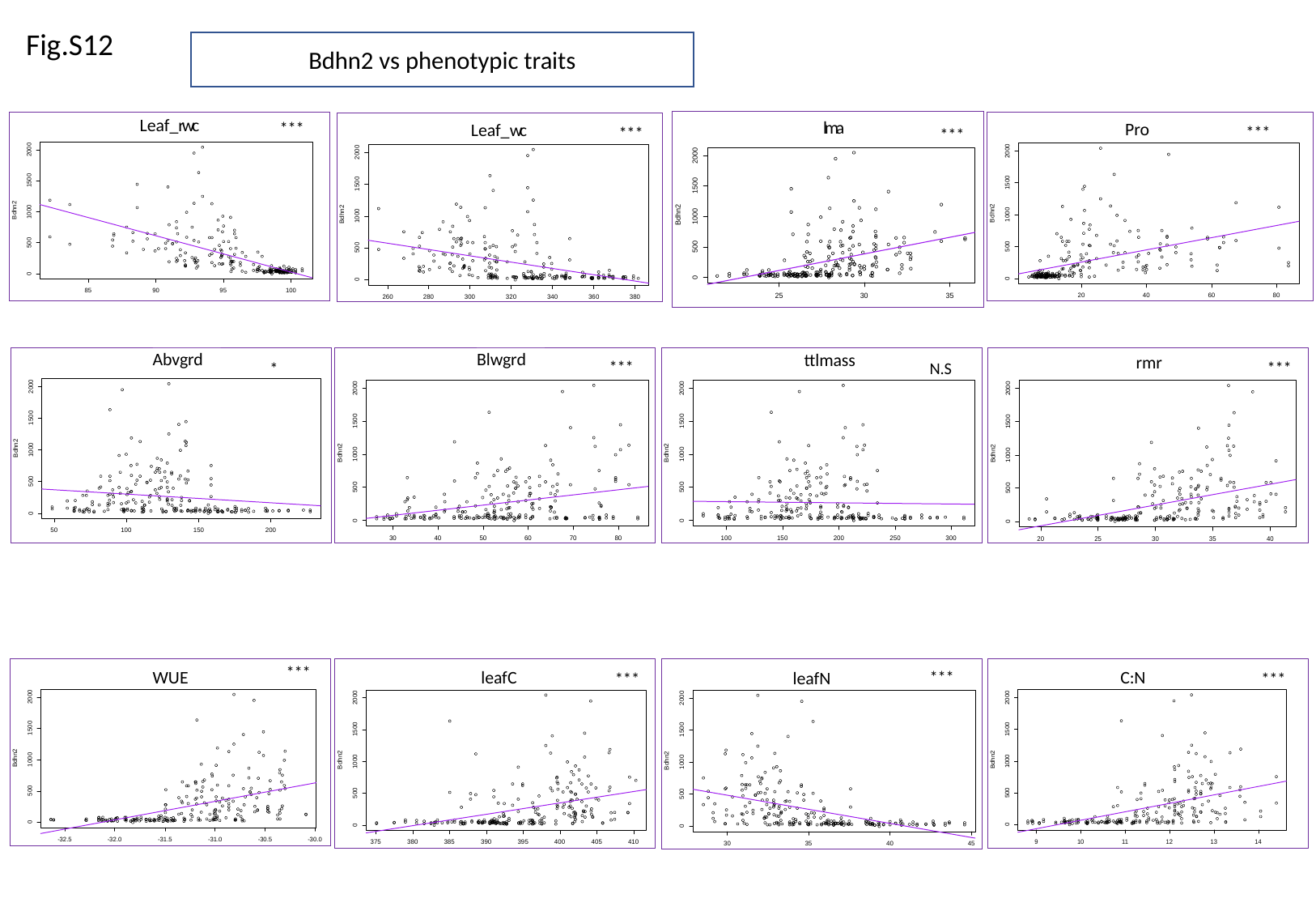

Fig.S12
Bdhn2 vs phenotypic traits
***
Leaf_rwc
***
Pro
***
Leaf_wc
lma
***
ttlmass
N.S
rmr
***
Abvgrd
*
Blwgrd
***
***
WUE
***
leafC
***
leafN
***
C:N

### Slide 19
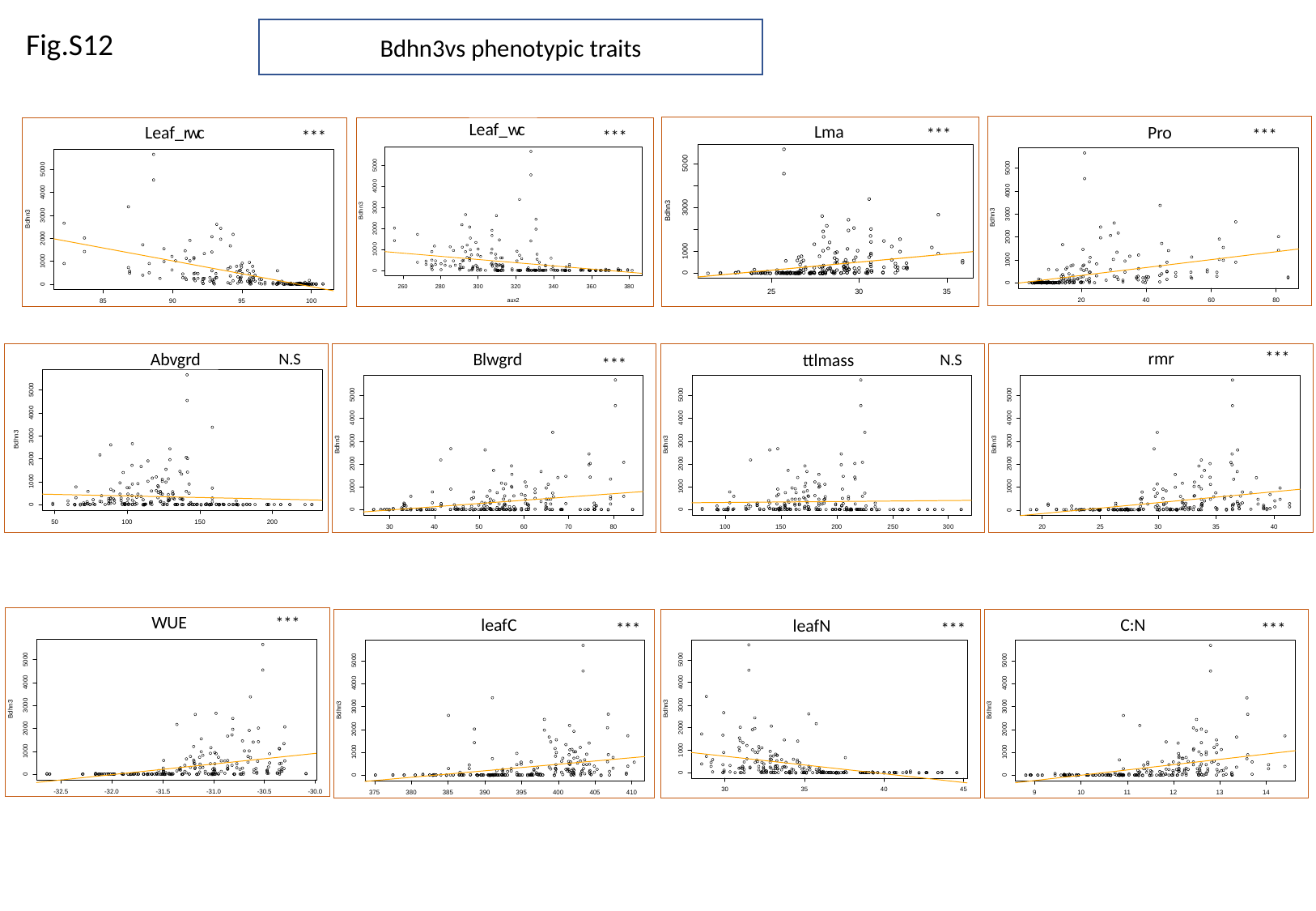

Fig.S12
Bdhn3vs phenotypic traits
***
Pro
***
Leaf_wc
***
***
Leaf_rwc
Lma
***
rmr
N.S
Abvgrd
N.S
ttlmass
***
Blwgrd
***
WUE
***
leafC
***
leafN
***
C:N

### Slide 20
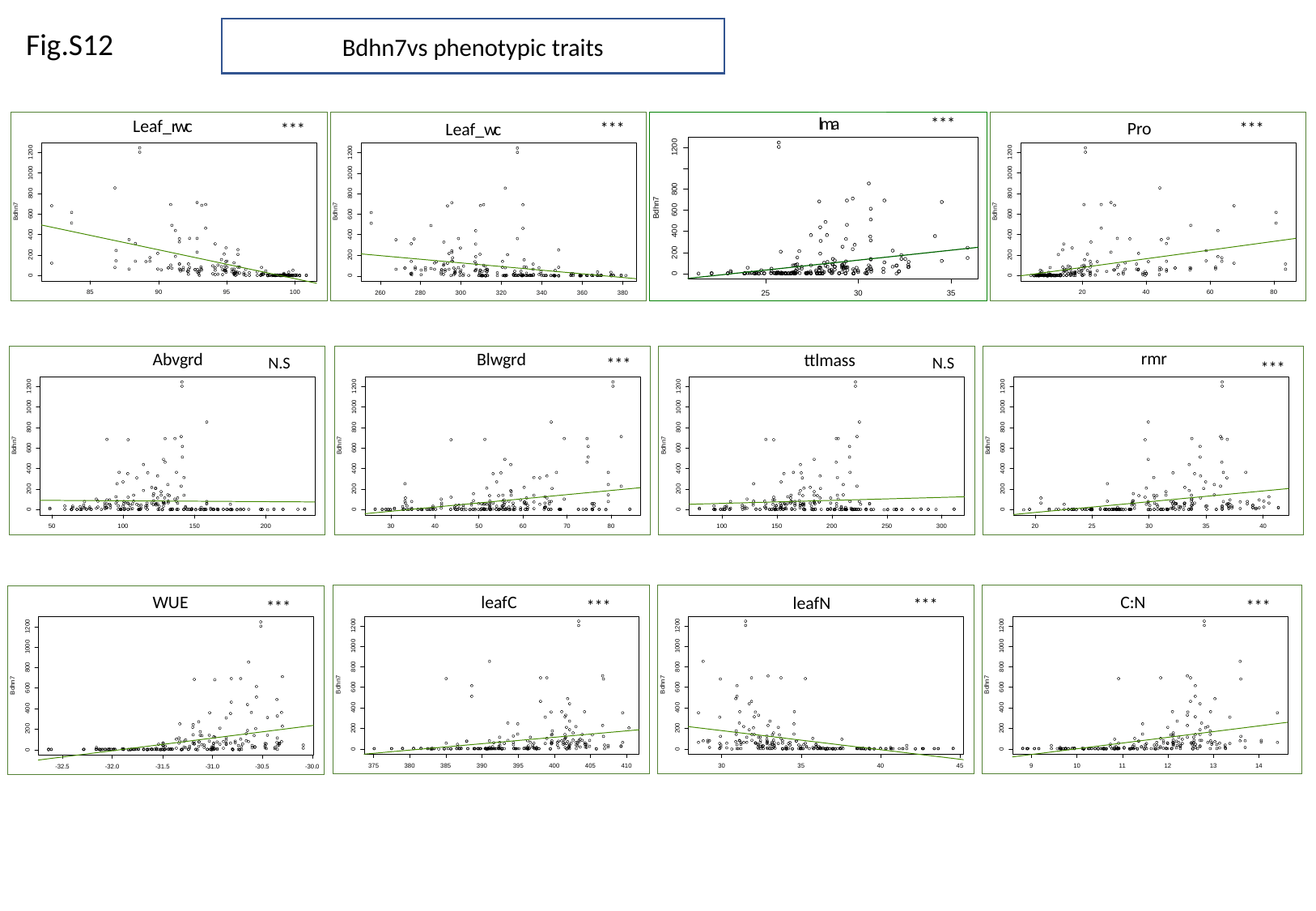

Fig.S12
Bdhn7vs phenotypic traits
***
***
Pro
***
Leaf_wc
***
Leaf_rwc
lma
Abvgrd
N.S
Blwgrd
***
ttlmass
N.S
rmr
***
***
leafC
***
leafN
***
C:N
***
WUE

### Slide 21
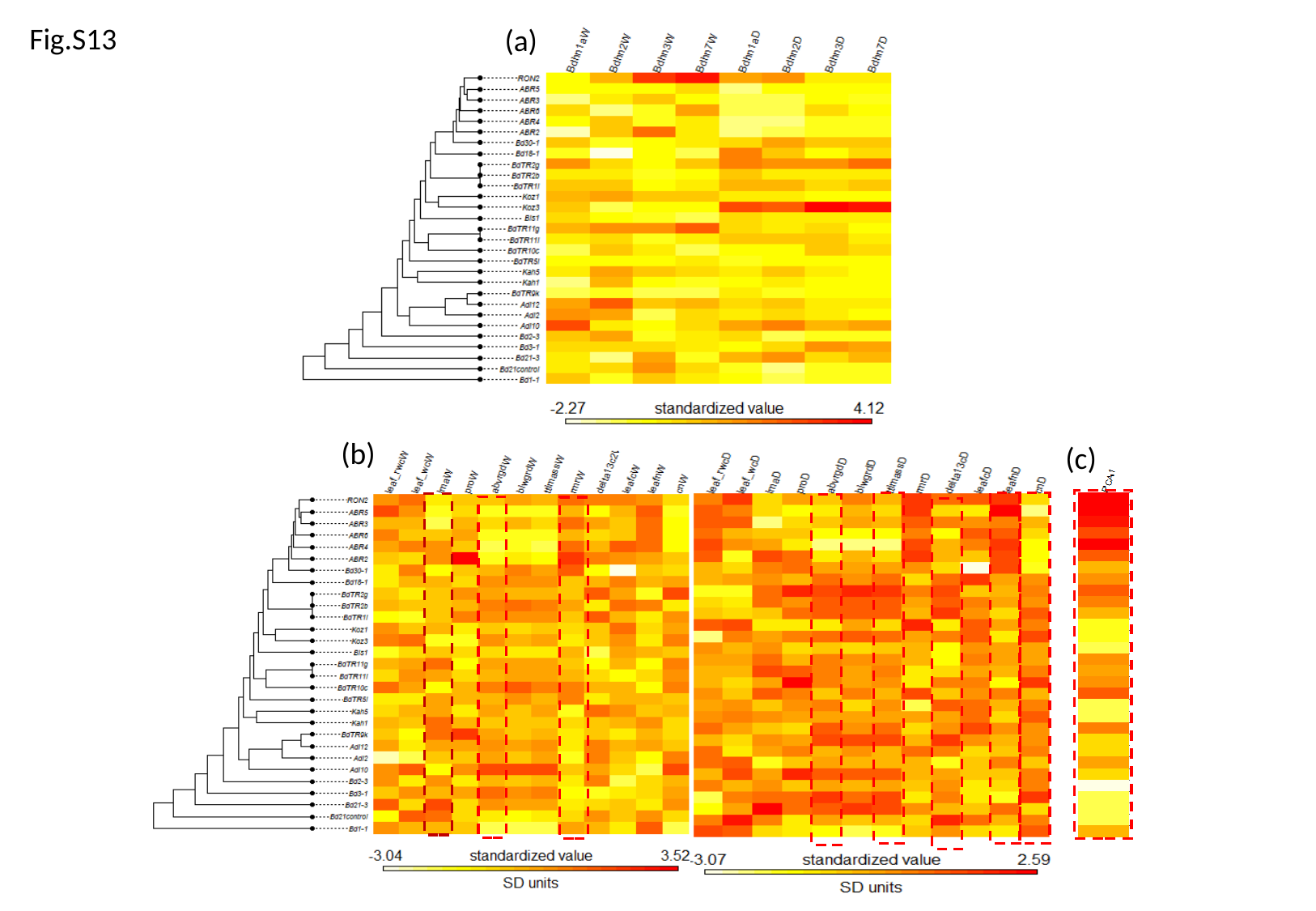

Fig.S13
(a)
(b)
(c)
